# Targeting Iron - Respiratory Reciprocity Promotes Bacterial Death

**DOI:** 10.1101/2024.03.01.582947

**Authors:** Mohammad Sharifian Gh., Fatemeh Norouzi, Mirco Sorci, Tanweer S Zaid, Gerald B. Pier, Alecia Achimovich, George M. Ongwae, Binyong Liang, Margaret Ryan, Michael Lemke, Georges Belfort, Mihaela Gadjeva, Andreas Gahlmann, Marcos M. Pires, Henrietta Venter, Thurl E. Harris, Gordon W. Laurie

**Author notes:** Equal contribution. Correspondence: Mohammad Sharifian Gh Gordon Laurie. Contact author: Gordon Laurie.

## Abstract

Discovering new bacterial signaling pathways offers unique antibiotic strategies. Here, through an unbiased resistance screen of 3,884 gene knockout strains, we uncovered a previously unknown non-lytic bactericidal mechanism that sequentially couples three transporters and downstream transcription to lethally suppress respiration of the highly virulent *P. aeruginosa* strain PA14 - one of three species on the WHO’s ‘Priority 1: Critical’ list. By targeting outer membrane YaiW, cationic lacritin peptide ‘N-104’ translocates into the periplasm where it ligates outer loops 4 and 2 of the inner membrane transporters FeoB and PotH, respectively, to suppress both ferrous iron and polyamine uptake. This broadly shuts down transcription of many biofilm-associated genes, including ferrous iron-dependent TauD and ExbB1. The mechanism is innate to the surface of the eye and is enhanced by synergistic coupling with thrombin peptide GKY20. This is the first example of an inhibitor of multiple bacterial transporters.

## INTRODUCTION

Prokaryotic respiration is largely the responsibility of evolutionarily ancient multisubunit transporters in their outer and/or inner membranes ^1^. Manipulation of transporters to limit bacterial respiration, and thereby pathogenesis, has been a decades long quest - particularly those that competitively acquire extracellular iron essential for energy production, oxygen transport, gene regulation and virulence. ^2^ Other targets involve polyamine uptake for growth, biofilm formation, swarming and siderophore synthesis ^3^, and transporters capable of antibiotic expulsion that in turn can enhance resistance.

Yet, among current preclinical anti-bacterial approaches few directly address transporters, including PHT-427 for Gram-positive and Gram-negative ferrous iron uptake ^4^ or GW3965·HCl only for Gram-positive ferrous iron uptake ^5^ as well as fluorophenylalkyl-substituted cyanoguanidine derivatives ^6^ and others in early development ^7^ that suppress drug efflux. None directly inhibit more than one transporter. In contrast, current antibiotic classes target cell wall synthesis (β-lactams ^8^ and glycopeptides ^9^) or depolarize the inner-membrane (lipopeptides ^10^) or alter the bacterial metabolome (pyrimidines ^11^ and sulfonamides ^12^) or inhibit replication (quinolones and fluoroquinolones ^13^) or transcription pathways (rifamycins ^14^). Many obstruct translational pathways (aminoglycosides ^15^, lincosamides ^16^, macrolides ^17^, streptogramins ^18^, oxazolidinones ^19^, phenicols ^20^, and tetracyclines ^21^).

Identifying an inhibitor of multiple transporters might be transformational as virulent pathogens have evolved many elegant ways to overcome host nutritional immunity ^2^. Direct uptake of the neutral pH soluble Fe^2+^ (ferrous) form of iron is mainly by the anaerobically induced transporter FeoB under conditions of hypoxia when Fe^2+^ is the most common form of iron ^22^. Its availability enhances the virulence of the opportunistic pathogen *P. aeruginosa* in individuals with cystic fibrosis ^23^. FeoB is widely represented in the inner membrane of Gram-negative bacteria (including *Enterobacteriaceae, Acinetobacter and Pseudomonas*) and in the sole membrane of Gram-positive bacteria (including *Staphylococci*) and in archaea, with modeling suggesting a GTP gated pore structure ^24^. Unlike Fe^2+^, the oxidized Fe^3+^ (ferric) form of iron is insoluble at neutral pH, although more abundant and the primary iron source when the host environment is normoxic. Bacterial uptake is indirect through secreted chelating siderophores that displace host Fe^3+^ sequestered on ferritin-, lactoferrin- and transferrin for capture by bacterial surface receptors and transport by the TonB:ExbB:ExbD ^25^ and lipoprotein siderophore-binding protein ^26^ systems on Gram-negative and Gram-positive bacteria, respectively.

Here we provide evidence for a direct inhibitor of both FeoB and the polyamine transporter PotH that also suppresses RNA expression of *exbB1* involved in Fe^3+^ uptake and the taurine ABC transporter WP_003114312.1. The inhibitor is the cationic lacritin peptide ‘N-104’ ^27^ that is innate to the surface of the eye and enhanced by synergistic coupling with thrombin peptide GKY20.

## RESULTS

### Lacritin ‘N-104’ does not promote bacterial cell lysis

We previously discovered ‘lacritin’, a pleiotropic tear, salivary, plasma and CSF protein, out of an unbiased biochemical screen to address the biological basis of the most common ocular surface disease ^28^. Lacritin transiently stimulates autophagy in the presence of inflammatory stress to restore oxidative phosphorylation ^29^ and promote both epithelial ^30^ and neuronal regeneration ^31^.

Interestingly, a bactericidal activity is potentiated by a lacritin cleavage product centered on the endogenous 15 amino acid cationic tear fragment ‘N-104’ ^27^ (or slightly smaller ‘N-107’ ^32^) that appears to play a central role in the innate defense of the eye ^27^. Most often, cationic antimicrobial peptides are considered to be bacterial lytic, with potential for more complex mechanisms ^33^. N-104 (pI 10.5) is contained within the 54 amino acid recombinant fragment ‘N-65’ (**Figure 1A**). N-65 (pI 9.9) allowed entry of membrane impermeable SYTOX green into *E. coli*, although metabolomic changes were incompatible with lysis ^27^. Curious about this observation and in particular the bactericidal mechanism, we reexamined membrane permeabilization first by monitoring surface plasmon resonance (SPR) as N-104 (2 - 25 µM) flowed for 100 s over two different chip-supported model membranes (**Figures 1B-1F and S1A**). Monitoring was continued after a buffer wash whereby disruptive peptides remove membrane lipid causing the SPR response to fall below the membrane-alone baseline by 600 s. SPR sensitivity is as little as 10 pg/ml ^34^ of dry mass. However, no significant disruption of the model membranes was detected (**Figures 1B-1C and S1A**). Although the model ‘Gram-positive’ membrane appeared to show some slight disruption at two lower N-104 concentrations, it was not confirmed at the highest concentration (**Figure S1A**). The experiment was then repeated using quartz crystal microbalance with dissipation monitoring (QCM-D) in which a decrease of frequency is proportional to increase in wet mass ^35^. This suggested a slight decrease in mass (**Figures S1B and S1C**) at levels not membrane disruptive. Thus, the bactericidal mechanism of the N-104 cleavage product is non-lytic.

**Figure 1.**
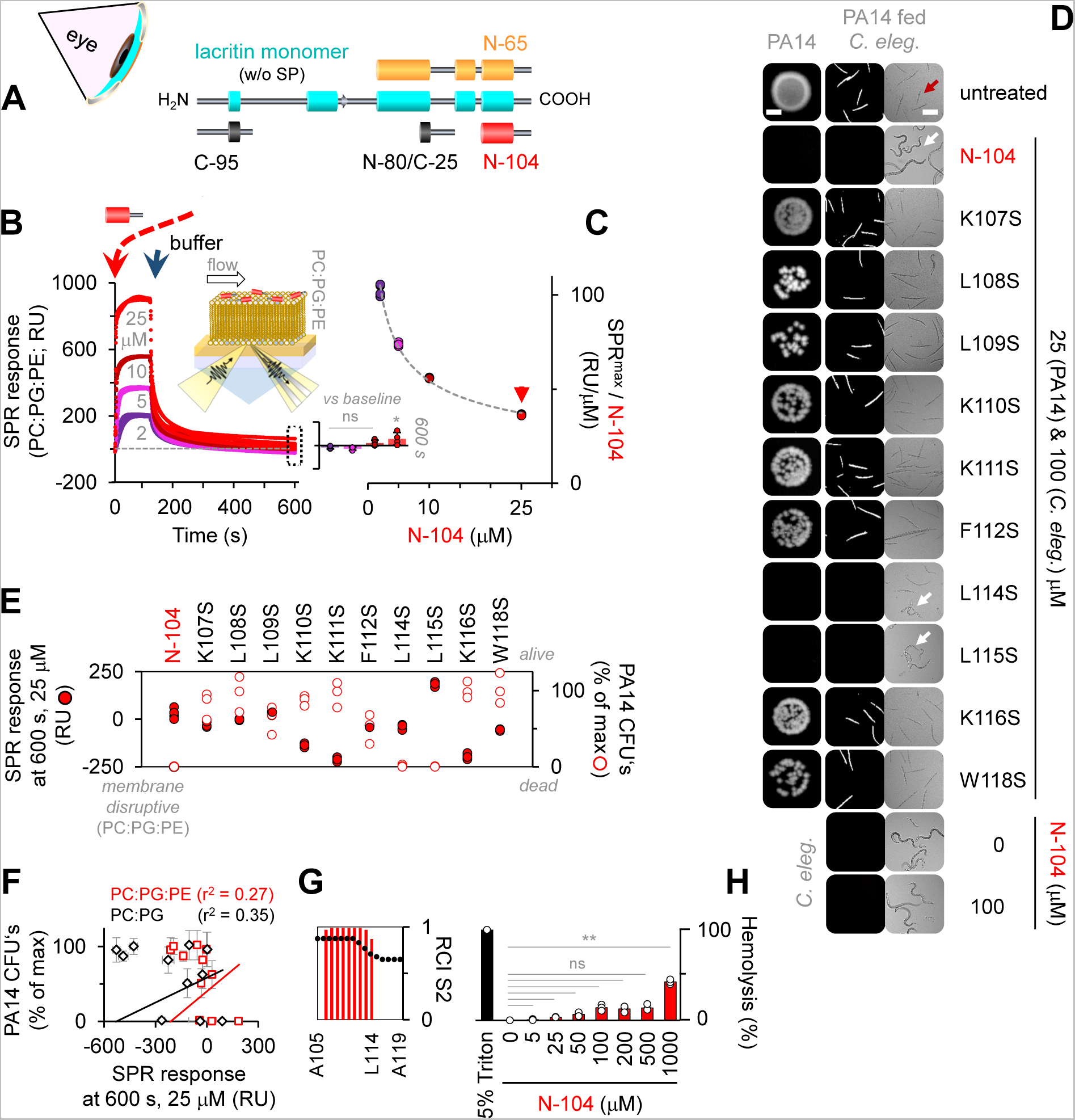
Lysis-free ‘N-104’ killing of virulent and multi-drug resistant *P. aeruginosa* strain ‘PA14’. (A) Diagram of lacritin with bioactive synthetic peptide ‘N-104’, recombinant ‘N-65’ and inactive ‘N-80/C-25’ and ‘C-95’. (B) N-104 interaction with Gram-negative model membrane as monitored by surface plasmon resonance (SPR). Inserts: diagram of SPR with N-104 on model membrane and SPR response at 600 s. Two-way ANOVA with Tukey’s multiple comparison test (n = 4 experiments). (C) Maximum SPR as a function of N-104 concentration. (D) N-104 amino acids necessary for killing determined by colony forming unit (CFU) and *C. elegans* pathogenesis assays. Scale bars are 2 mm and 200 μm, respectively. (E, F) Correlation assay of N-104 or N-104 analog bactericidal activity versus membrane disruption. (SPR: n = 4 experiments, CFU: n = 3 experiments). (G) ^1^H-NMR of N-104. (H) Assay of N-104 hemolytic activity versus positive control 5% Triton X-100. Nonparametric Friedman test with Dunn’s multiple comparisons test (n = 3 experiments). Data represent the mean ± SD, **p<0.01, *p<0.05, ns, non-significant. See also **Figure S1**.

Some non-lytic antimicrobial peptides utilize multiple tryptophans or both arginines and prolines or a single helix disruptive proline to gain bacterial entry ^36^. N-104 lacks arginine and contains only a single proline and tryptophan, both in a coiled-coil domain C-terminal to an amphipathic α-helix (**Figures 1G and S1F, S1J-K**). We synthesized ten N-104 analogs, each with a single serine substitution for lysines, leucines, phenylalanine and tryptophan. Most membrane disruptive by SPR were N-104:K111S and N-104:K116S, but neither substantially diminished growth of the highly virulent, multi-drug resistant PA14 strain ^37^ of *P. aeruginosa* as examined by colony forming unit (CFU) (**Figures 1D and 1E**) and *C. elegans* feeding assays (**Figures 1D and S1G-I**). Comparison of all ten analogs revealed no correlation between diminished CFU’s and either SPR (**Figure 1F**) or QCM-D response (**Figures S1D and S1E**). Only N-104:L114S and N-104:L115S retained full bactericidal activity at a minimum inhibitory concentration (MIC) of 2 μM (3.6 µg/ml). Thus, all amino acids with basic side chains and most with nonpolar side chains contribute to the N-104 activity that is also not hemolytic (**Figure 1H**). Curious about resistance that often develops by 10 generations ^38^ - particularly when antibiotic exposure is below the minimal inhibitory concentration - we tested N-104 on PA14 through 30 generations. No stable resistance was apparent (**Figure S2**).

### Mutants lacking transporters FeoB (Fe^2+^) and PotH (polyamine) are N-104 resistant

Despite newly apparent transcriptional effects downstream of deleted genes in bacterial libraries of single gene knockout strains ^39^, unbiased screening of such libraries has proven to be powerfully insightful ^40,41^. Accordingly, we screened the Keio *E. coli* K12 collection of 3,884 single gene knockout strains ^42^ for resistance to N-104 (**Figure 2A**) using the reduction of resazurin in Alamar Blue as a measure of bacterial metabolic activity. Recognizing that the metabolic activity of each knockout strain may differ, a screening strategy was developed that compared the five hour Alamar Blue detectable metabolic activity of each before and after N-104 treatment (**Figure 2B**). A treated:untreated metabolic slope ratio of 0.75 or greater was considered to be indicative of N-104 resistance (**Figure 2C**) at a relatively high 100 µM (180 µg/ml) dose to minimize ambiguous outcomes. As respective negative and positive controls we took advantage of the 14-mer ‘N-80/C-25’ from an inactive region of lacritin (**Figure 1A**) and the beta-lactam antibiotic ampicillin, both at 100 µM (**Figure 2B**). Duplicate N-104 screens of the entire collection yielded 122 resistant candidates (**Figure 2D**). These were further screened twice (**Figures 2E and F**) thereby narrowing the list to 10 (**Figure 2G**) - each with a Z-score ≥ 1.85. Resistance was attributable to absence of: (i) the inner membrane ferrous iron transporter FeoB, (ii) the inner membrane polyamine transporter subunit PotH, (iii) the colanic acid polymerase WcaD involved in biofilm formation, (iv) N-acetyl transferase YjhQ, (v) ‘putative glucose transporter regulator’ YeeI (pseudogene), (vi) DUF986 protein YobD, or to one of four regulatory transcription factors: (vii) SgrR family member YbaE, (viii) ParB-like nuclease domain containing YbdM, (ix) GntR family member RluE (YmfC), or (x) hypothetical YhfZ. BLAST of each against the non-redundant protein sequences database (**Table S1**) revealed a common distribution among *Enterobacteriaceae*, and some also in *Acinetobacter baumannii (*FeoB, PotH, YobD, YbdM, RluE, YhfZ) and *P. aeruginosa* (FeoB, PotH, WcaD, YbdM, RluE) - all Gram-negative opportunistic pathogens. Heightened antibiotic development against multi-drug resistant strains of each is a WHO-recognized worldwide need. We chose to focus our efforts on FeoB and PotH since both are common to all three pathogen groups and together with RluE, a 23S rRNA pseudouridine synthase (**Table S1**), are the only three expressed in multi-drug resistant *P. aeruginosa* PA14 of the ten. We therefore tested PA14 transposon mutants ^43^ of FeoB and PotH (**Figure 2H**) and observed similar N-104 resistance (**Figures 2I and S2D**). Thus inner membrane proteins FeoB and PotH mediate N-104 bactericidal activity. Curiously, numerous inner membrane proteins with or without a periplasmic domain are suggested (BioGRID and/or STRING) to bind *E. coli* FeoB and PotH (**Table S2**) for which PA14 orthologs for the most part exist. Does N-104 act directly by gaining access to the periplasm, the ~25 nm wide protein and peptidoglycan rich region between the outer and inner membranes of Gram-negative bacteria? Or does it act indirectly? We first explore access and then the interaction mechanism and subsequently the consequences.

**Figure 2.**
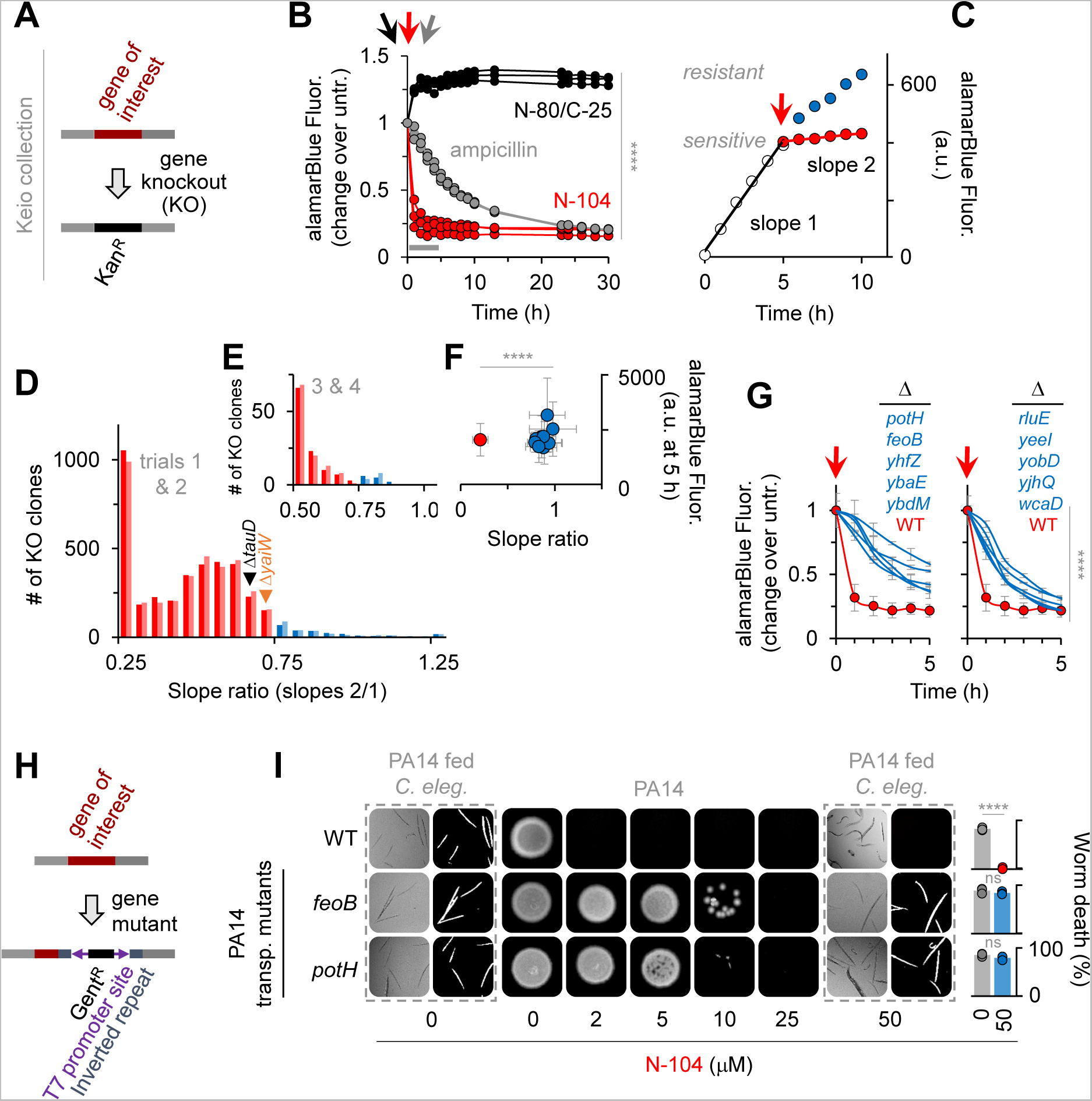
N-104 killing involves both an iron and a polyamine transporter. (A) Keio *E. coli* K12 gene knockout library. (B) *E. coli* K12 viability after treatment with N-104, N-80/C-25 (negative control) or ampicillin (positive control). Two-way ANOVA with Dunnett’s multiple comparisons test (n = 3 experiments). (C) Resistance screen relative slope ratio before (slope 1) and after (slope 2) N-104 treatment (model example). (D) 120 mutant strains display a slope ratio ≥ 0.75 indicative of N-104 resistance, identified from duplicate screens of the entire Keio library. (E) Resistant mutant strains from (D) narrowed by duplicate screens. (F) Combined slope ratios versus alamarBlue fluorescence at five hours of (E; blue) resistant compared to wild type (red) strains. One-way ANOVA (n = 4 experiments). (G) Viability assays of the 10 resistant mutants, inclusive of *E. coli* K12 lacking ferrous iron transporter FeoB and the PotH subunit of the PotFGHI polyamine transporter. Two-way ANOVA (n = 3 experiments). (H) *P. aeruginosa* PA14 Transposon Insertion Mutant Library. (I) Analogous *feoB* and *potH* PA14 mutants are also N-104 resistant. Two-way ANOVA with Šidák’s multiple comparisons test (n = 3 experiments). Data represent the mean ± SD, ****p<0.0001, ns, non-significant. See also **Figure S2** and **Table S1**.

### N-104 outer membrane translocation is facilitated by transporter YaiW

PA14 cells were monitored by imaging flow cytometry one hour after treatment with FITC-N-104 in the absence or presence of ethanol permeabilization, or with negative control FITC-’C-95’, a 24-mer synthetic peptide (**Figures 1A**) from lacritin’s inactive N-terminus. In contrast to FITC-C-95, substantial intracellular FITC-N-104 fluorescence was apparent at either 100 (**Figures 3A and 3B**) or 10 (**Figure 3B**) µM concentrations, although less than with positive control ethanol permeabilization. Since N-104 was incapable of membrane disruption (**Figure 1**) and FITC was not membrane permeable, we explored the possible role of an outer membrane transporter. From PA14 genome screening, ABC transporters of peptide-opine-nickel, di- and tripeptides and of peptidyl nucleoside antibiotics have been discovered; however, no transport of four tested antimicrobial peptides was apparent ^44^. We therefore considered outer membrane protein biotinylation for attempted N-104 affinity purification followed by tandem mass spectrometry. However, this approach is complicated by biotin permeation through outer membrane porin channels ^45^. There is also the possibility that biotinylation could alter binding properties ^46^. Precleared whole cell PA14 octylglucoside lysates were therefore incubated overnight with bead immobilized N-104 (versus C-95) under physiological conditions (**Figure 3C**). Following washes and progressive step gradient elution through 50, 75, 100, 125, 150, 300, 500 mM KCl for stringency, the 500 mM KCl eluate was subjected to tandem mass spectrometry (**Table S3**). The only outer membrane protein identified in the 500 mM KCl eluate off N-104 was ‘YaiW’ (**Figures 3C-D and S3A-B**), a lipoprotein. In *E. coli*, YaiW together with inner-membrane SbmA facilitates the non-lytic entry of arginine and proline-rich bovine cathelicidin antimicrobial peptide (Bac7(1-35)) ^47^. Our *E. coli* resistance screens show both ΔYaiW (0.70) and ΔSbmA (0.69) cells to be just below the arbitrary resistance cutoff of 0.75 (**Figure 2D**), however SbmA is absent from PA14 and BLAST failed to detect any homologs. Nonetheless, the PA14 transposon mutant lacking YaiW is partially resistant (**Figures 3E and S3B**). Thus YaiW facilitates N-104 transport across the outer membrane.

**Figure 3.**
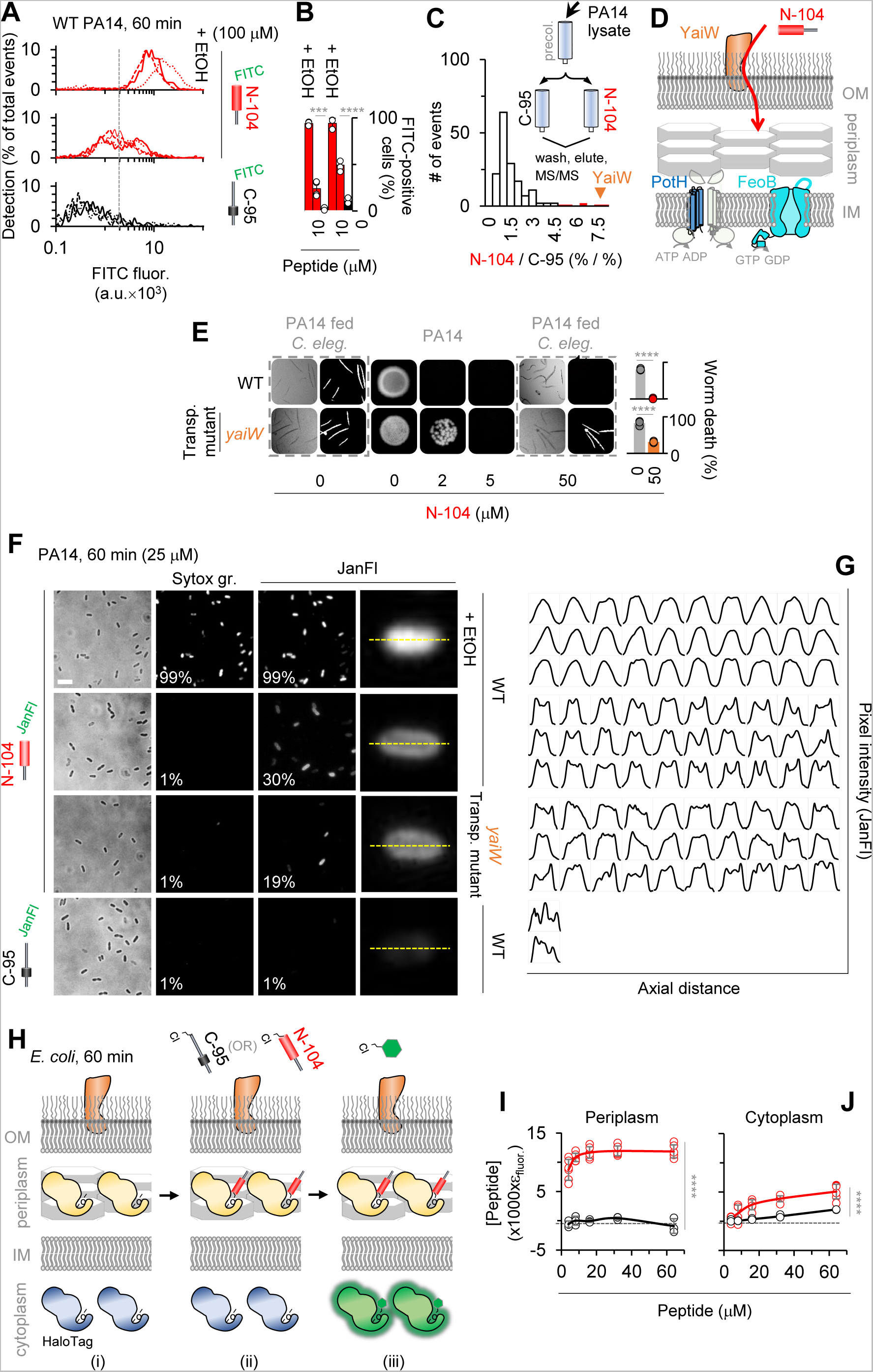
N-104 killing requires N-104 transport across the outer membrane into the periplasm. (A) Imaging flow cytometry of FITC-N-104 and FITC-C-95 (negative control) treated PA14 cells versus positive control ethanol permeabilized cells. (B) Dose dependence of (A). Two-way ANOVA with Šidák’s multiple comparisons test (n = 3 experiments). (C) Selective affinity of N-104 with tandem mass spectometry identified outer membrane lipoprotein ‘YaiW’, previously known to transport Bac7(1-35) peptide ^47^. (D) Hypothetical YaiW-mediated translocation of N-104 through the outer membrane to reach inner membrane FeoB and PotH. (E) PA14 YaiW transposon mutant is N-104 resistant. Two-way ANOVA with Šidák’s multiple comparisons test (n = 3 experiments). (F) Wide-field fluorescence imaging of Janelia Fluor 549-N-104 and -C-95 (negative control) treated PA14 or PA14 YaiW transposon mutant cells. SYTOX green controls for Janelia Fluor 549 permeability. (G) Single cell axial pixel intensity tracings of cells in (F). (H) Schematic of Bacterial ChloroAlkane Penetration Assay in *E. coli* in which chloroalkane tagged N-104 and C-95 (negative control) compete with chloroalkane modified fluorescent dye for cells expressing either HaloTag periplasmic or cytoplasmic marker. (I) Results of with HaloTag periplasmic marker. Two-way ANOVA with Šidák’s multiple comparisons test (n = 3 experiments). (J) Results with HaloTag cytoplasmic marker. Two-way ANOVA with uncorrected Fisher’s least significant difference test (n = 3 experiments). Data represent the mean ± SD, ****p<0.0001, ***p<0.001. See also **Figure S3 and Table S3**.

In enteric bacteria, SbmA’s contribution is the periplasm-to-cytoplasm translocation of Bac7(1-35) and other antimicrobial peptides ^47^. Lacking SbmA, N-104 might therefore accumulate in the periplasm. PA14 cells were imaged by wide-field fluorescence microscopy one hour after treatment with Janelia Fluor 549 conjugated N-104 in the absence or presence of ethanol permeabilization, or instead with Janelia Fluor 549 conjugated-C-95 (controlling for labeling effects in **Figure S3C**). We observed an internal ring of Janelia Fluor-N-104 staining (**Figures 3F**) as documented by pixel intensity measurements along with axial distance of individual cells (**Figure 3G**). This pattern is similar to that observed for GFP targeted to the periplasm of *E. coli* by a TorA signal sequence ^48^. Internal to the ring (presumably the cytoplasm) fluorescence was apparent, but less. In the absence of YaiW, the frequency of PA14 cell staining was substantially reduced. Janelia Fluor-C-95 cells lacked staining and with ethanol permeabilization ^49^ Janelia Fluor-N-104 was almost entirely cytoplasmic (**Figures 3F and 3G**). Since the Janelia Fluor 549 label is membrane permeable, we also monitored PA14 for SYTOX green uptake. None was detected except in the presence of ethanol (**Figure 3F**), thereby further validating the lack of N-104-dependent membrane disruption as per SPR (**Figure 1**).

The internal ring of Janelia Fluor-N-104 staining was suggestive of periplasm accumulation. To explore this using a different approach, advantage was taken of the newly developed Bacterial ChloroAlkane Penetration Assay (BaCAPA), currently available only in *E. coli* ^50^. Competition of chloroalkane tagged N-104 (or as negative control C-95) with chloroalkane modified fluorescent dye in the presence of cells expressing a HaloTag periplasmic marker or alternatively of other cells expressing a HaloTag cytoplasmic marker is evidence of location (**Figure 3H**). Chloroalkane tagged N-104 (but not C-95) treatment of *E. coli* expressing a HaloTag periplasmic marker successfully outcompeted the chloroalkane modified fluorescent dye to bind the HaloTag periplasmic marker - an outcome indicative of a periplasmic localization (**Figures 3I and top row of S3D**). Competition of tagged N-104 (but largely not C-95) for the HaloTag cytoplasmic marker was considerably less, suggesting some cytoplasmic entry (**Figures 3J and top row of S3E**). As a positive control for both periplasmic and cytoplasmic markers, a ChloroAlkane tagged ligand was included (**Figures S3D-S3E, bottom row**). The analytical approach underlying **Figures 3I and 3J** is provided in the Supplementary Information.

### N-104 ligates FeoB and PotH ion channel-associated periplasmic loops

FeoB is predicted to extend five (’TMHMM v 2’) or four (’DeepTMHMM 1.0.24’; lacks loop 2) loops into the periplasm anchored respectively by ten or eight transmembrane domains forming an Fe^2+^ transport channel with cytoplasmic G-protein and GDP dissociation inhibitor domains. The PotH subunit contributes to a polyamine channel through six transmembrane domains connected to three predicted (’DeepTMHMM 1.0.24) periplasmic loops. The channel transports primarily spermidine in *P. aeruginosa* and putrescine in *E. coli*. To ask whether periplasmic N-104 targets each directly, advantage was taken of the same immobilized N-104 versus C-95 affinity approach to which was applied lysates of overexpressed recombinant FeoB (PAO1 strain of *P. aeruginosa*) or PotH (*E. coli*), both with His tags (**Figures 4A-4D**). Identity to PA14 orthologs are respectively 99% and 65%. Incubation was under physiological conditions. After extensive washing, both FeoB and PotH eluted off N-104 with 0.5 and 1 M KCl, but not with 0.3 M KCl, indicative of high affinity ligation (**Figures 4E-4H and S4**). Neither bound to C-95. Single loop deletion mutants (**Table S4**) were then developed from the TMHMM v 2 predictions (**Figures S4D and S4P**). For FeoB, expression of each proved problematic (**Figure S4E**) with the exception of deleted outer loop 4, although with a lower yield. Column loading was thereby compensated. Loop 4 comprises one of two predicted FeoB ‘gate motifs’ that channel Fe^2+^ across the inner membrane ^24,51^. Deletion of loop 4 completely abrogated affinity for N-104, as confirmed by substitution of outer loop 4 with outer loops 2, 3 or 5 (**Figures 4E[i]-4F and S4F-S4I**). Although synthesis of a 131 amino acid outer loop 4 peptide was not attempted, we next generated synthetic peptides corresponding to outer loop 1 (INIGGALQP), outer loop 2 (LEDSGYMARAAFVMDRLMQ), outer loop 3 (GAFFGQGGA) and outer loop 5 (ATFAA). Each was then evaluated at increasing concentrations as competitive inhibitors of N-104 - FeoB targeting. However, outer loop 1 and 3 peptides were not water, DMSO, methanol, isopropanol nor acetonitrile soluble. Solubility of outer loop 1 peptide in formic or acetic acid, and outer loop 3 in ammonia water and acetic acid was identified, however N-104 - FeoB ligation was not possible in each. Interestingly, the lacritin targeting ‘GAGAL’ domain in our human syndecan-1 synthetic peptide ‘pep19-30’ ^52^ is similar to respective hydrophobic outer loop 1 and 3 sequences ‘GGAL’ and ‘GA(LA)’ with ‘LA’ membrane associated. It was therefore assayed, however neither outer loops 2 nor 5, nor pep19-30 were inhibitory, as per lack effect on 1 M KCl elutable FeoB from N-104 beads (**Figures 4E[ii]-4F and S4J-S4L**). FeoB’s outer loop 4 thus appears to be the target of N-104.

**Figure 4.**
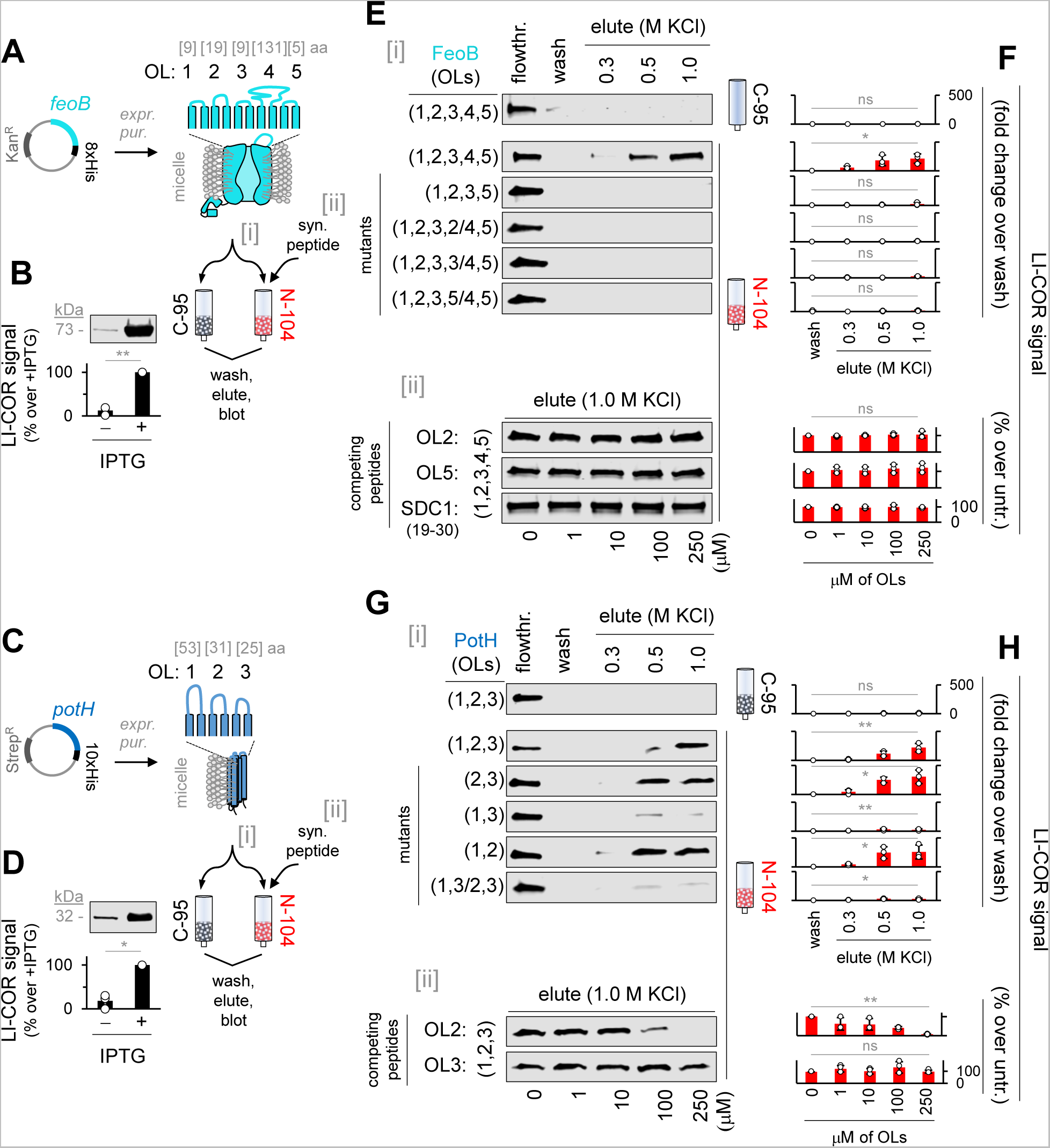
N-104 targets periplasmic loops 4 and 2 respectively of FeoB and PotH. (A) FeoB expression construct and TMHMM (v 2) predicted membrane structure. (B) Western blot of IPTG-induced FeoB and passage over N-104 vs C-95 (negative control) columns, the former in the absence or presence of FeoB synthetic peptides corresponding to periplasmic loops. Unpaired t-test with Welch’s correction (n = 3 experiments). (C) PotH expression construct and TMHMM (v 2)/Deep TMHMM (1.0.24) predicted membrane structure. (D) Western blot of IPTG-induced PotH and passage over N-104 vs C-95 (negative control) columns, the former in the absence or presence of PotH synthetic peptides corresponding to periplasmic loops. Unpaired t-test with Welch’s correction (n = 3 experiments). (E) Western blots of flowthrough, final wash fraction or KCl eluants from N-104 or C-95 columns of (i) FeoB or FeoB deletion/substitution mutants, or (ii) FeoB with periplasmic (outer) loop synthetic peptides or SDC1 peptide. (F) Analysis of (E). Friedman ANOVA (n = 3 experiments). (G) Western blots of flowthrough, final wash fraction or KCl eluants from N-104 or C-95 columns of (i) PotH or PotH deletion/substitution mutants, or (ii) PotH with periplasmic (outer) loop synthetic peptides. (H) Analysis of (G). Friedman ANOVA (C-95; 2,3; 1,3; 1,2; 1,3/2,3), Kruskal-Wallis ANOVA (OL2; OL3) (n = 3 experiments). Data represent the mean ± SD, **p<0.01, *p<0.05, ns, non-significant. See also **Figure S4 and Table S4**.

Among PotH’s three predicted periplasmic loops, deletions suggest that only loop 2 is targeted by N-104 as confirmed by substitution of outer loop 2 with outer loop 3 (**Figures 4G[i]-4H and S4M-S4T**). Further, soluble outer loop 2 but not outer loop 3 synthetic peptides (respectively WMGILKNNGVLNNFLLWLGVIDQPLTILHTN and ELLGGPDSIMIGRVLWQEFFNNRDW) competitively inhibited the capacity of N-104 to ligate PotH (**Figures 4G[ii]-4H and S4U-S4V**). PotH together with PotI constitute the transmembrane polyamine channel of the ABC transporter PotFGHI ^53^. In aggregate, N-104 uniquely targets the periplasmic portion of the FeoB Fe^2+^ and PotH/PotI polyamine channels.

### N-104 ligation suppresses Fe^2+^ and polyamine uptake

Current antibiotics do not address transporters, nor do any in preclinical development with the exception of GW3965·HCl that inhibits the GTPase and ATPase activity of *S. aureus* FeoB and growth of several other Gram-positive but not Gram-negative bacteria ^5^, and PHT-427 for Gram-positive and Gram-negative ferrous iron uptake ^4^. Others suppress drug efflux ^7^. To explore whether ligation is negatively coupled with transport (**Figures 5A-5F**), PA14 iron ^54^ and both spermidine and putrescine uptake assays were initiated. 2,2’-bipyridyl depletion of intracellular iron facilitated measurement of exogenous Fe^2+^ uptake through the chelation-dependent colorimetric change in ferrozine ^55^. Uptake was (0.55 ± 0.03) × 10^6^ without and (2.49 ± 0.19) × 10^6^ Fe^2+^ ions per cell with exogenous Fe^2+^ (**Figures 5B and S5A**). 10 and 25 µM N-104 inhibited exogenous Fe^2+^ uptake respectively by 47 and 61% whereas negative control C-95 was ineffective (**Figure 5C**). We also tested negative control peptide N-80/C-25 representing a lacritin region immediately N-terminal to N-104 (**Figure 1A**) that like N-104 is cationic. N-80/C-25 was partially inhibitory but not in a dose dependent manner (**Figure 5C**), and displayed almost no effect in CFU assays (**Figures 5G and S5B**). PA14 lacking FeoB displayed a 37% decrease in Fe^2+^ uptake (**Figure S5C**) in keeping with compensation by alternate transport systems. Similar uptake studies were performed with ^14^C-spermidine and -putrescine. Uptake of ^14^C-spermidine in PA14 did not differ from that of ^14^C-putrescine (**Figures 5E and S5D-S5E**). Like Fe^2+^, pretreatment with 10 and 25 µM N-104, but not C-95, (**Figure 5F**) inhibited uptake of ^14^C-spermidine by 54 and 66%. Cationic N-80/C-25 showed almost similar inhibitory effect (**Figure 5F**). In eukaryotic cells, polyamines appear to be capable of chelating iron for uptake - possibly by a polyamine transporter ^56^ such as ATP13A3 ^57^. Limited exploration of this concept confirmed that spermidine but not putrescine chelates Fe^2+^ (as also recently observed ^58^), but not Mg^2+^, Ca^2+^, Co^2+^, Na^+^ nor K^+^ (**Figures S5F-S5H**). Moreover, exogenous spermidine, but not putrescine, inhibited N-104 bactericidal activity in a concentration-dependent manner (**Figures 5H-5I and S5I**) at or below 100 µM (**Figures S5J-S5K**). Collectively, the process of N-104 binding to FeoB and PotH impedes the cellular uptake of Fe^2+^ and spermidine, thereby promoting bacterial death.

**Figure 5.**
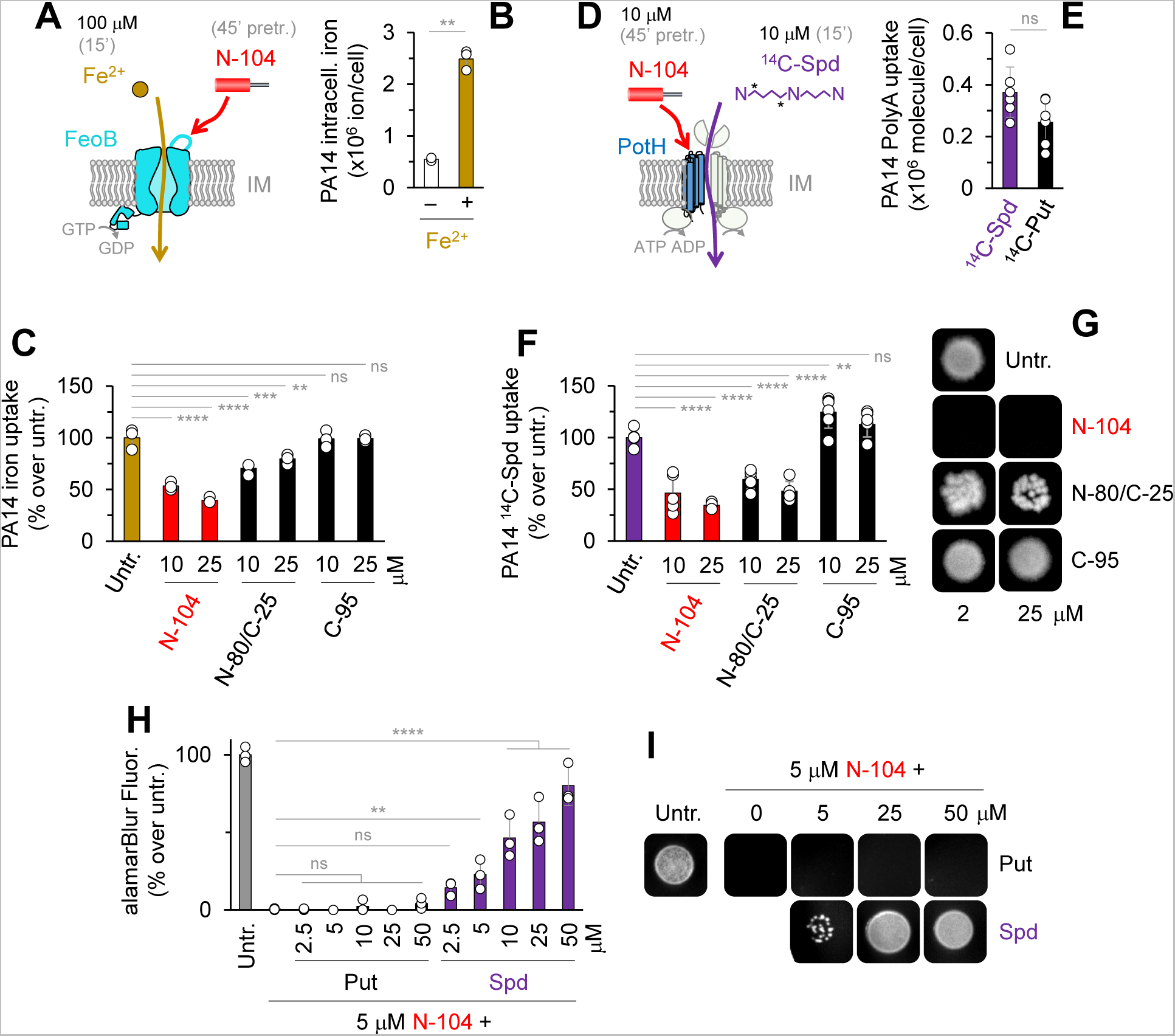
By targeting FeoB and PotH, N-104 suppresses ferrous iron and spermidine uptake. (A) Pretreatment with N-104 (or N-80/C-25 or C-95 as negative controls) prior to Fe^2+^ supplementation. (B) PA14 intracellular iron in the absence and presence of supplemented Fe^2+^. Unpaired t-test with Welch’s correction (n = 3 experiments). (C) Comparative iron uptake by untreated, N-104, or negative control N-80/C-25 or C-95 treated PA14 cells. Two-way ANOVA with Dunnett’s multiple comparisons test (n = 3 experiments). (D) Pretreatment with N-104 (or N-80/C-25 or C-95 as negative controls) prior to ^14^C-spermidine (^14^C-Spd) supplementation. (E) PA14 uptake of supplemented ^14^C-spermidine (^14^C-Spd) and ^14^C-putrescine (^14^C-Put). Unpaired t-test (n = 3 experiments). (F) Comparative ^14^C-spermidine uptake by untreated, N-104, or negative control N-80/C-25 or C-95 treated PA14 cells. Two-way ANOVA with Dunnett’s multiple comparisons test (n = 3 experiments). (G) N-104, but not N-80/C-25 nor C-95, killing of PA14. (H) N-104 killing of PA14 in the presence of excess putrescine, but not spermidine. Two-way ANOVA with Dunnett’s multiple comparisons test (n = 3 experiments). (I) N-104 killing of PA14 in the presence of excess putrescine, but not spermidine. Data represent the mean ± SD. ****p<0.0001, ***p<0.001, **p<0.01, ns, non-significant. See also **Figure S5**.

### N-104 synergizes with thrombin antimicrobial peptide ‘GKY20’

Whereas N-104 appears to play a central role in the innate defense of the eye as a cleavage-potentiated lacritin fragment ^27^, it likely does not act in isolation. Synergism among antimicrobial peptides, some with remarkable functional specificity, is well established ^59^ and at least twenty-two other putative or demonstrated antimicrobial proteins have been detected in human tears ^60^. It is also apparent that N-104’s efficacy diminishes as osmolarity increases (**Figures 6A and S6A**) despite its high affinity interactions with YaiW, FeoB and PotH (fully elutable only at 500 mM KCl); and earlier evidence ^27^. To explore possible synergies, we subjected precleared human tears to N-104 versus negative control C-95 affinity columns. Following extensive PBS washing, 500 mM KCl eluants were subjected to tandem mass spectrometry (**Figures 6B and 6C**) with specificity explored after subtraction of C-95 interacting proteins from those bound to N-104 (**Table S5**). Affinity with 10 proteins reported to have antimicrobial activity was detected. The literature was most convincing for the GKY20 (GKYGFYTHVFRLKKWIQKVI) peptide of thrombin, F2 ^61^ (**Figures 6D and S6B**), the QRLFQVKGRR peptide of gelsolin (GSN ^62^) and the SSSKEENRIIPGGI peptide of cystatin S (CST4 ^63^). GKY20 overlaps with antimicrobial FYT21 released from thrombin by *P. aeruginosa* elastase ^64^. Each synthetic peptide was synthesized for incubation with N-104 in a modified checkerboard challenge of PA14. Osmolarity was 300 mOsm/l. Only GKY20 was synergistic (**Figures 6E and S6C, S6D**), in keeping with its carpet-like mechanism of lipid disruption as studied on model membranes ^65^. N-104 binds thrombin (**Figure 6D**). Does it also directly interact with GKY20 in thrombin’s C-terminus? To address this question, GKY20 was immobilized on nitrocellulose for incubation with N-104 versus C-95 (300 mOsm/l). Only N-104 bound (**Figures 6F and S6E**). This assay was repeated with each of the ten N-104 serine-substituted analogs from **Figure 1D and 1E**. Single serine substitution was ineffective at disturbing the interaction (**Figures 6G and S6F**). Partially effective was multiple serine substitution of L108, L109, L114, L115, F112 and W118 (’LFW:S’; **Figures 6H and S6G-S6H**) however it alone was not bactericidal (**Figure S6I**).

**Figure 6.**
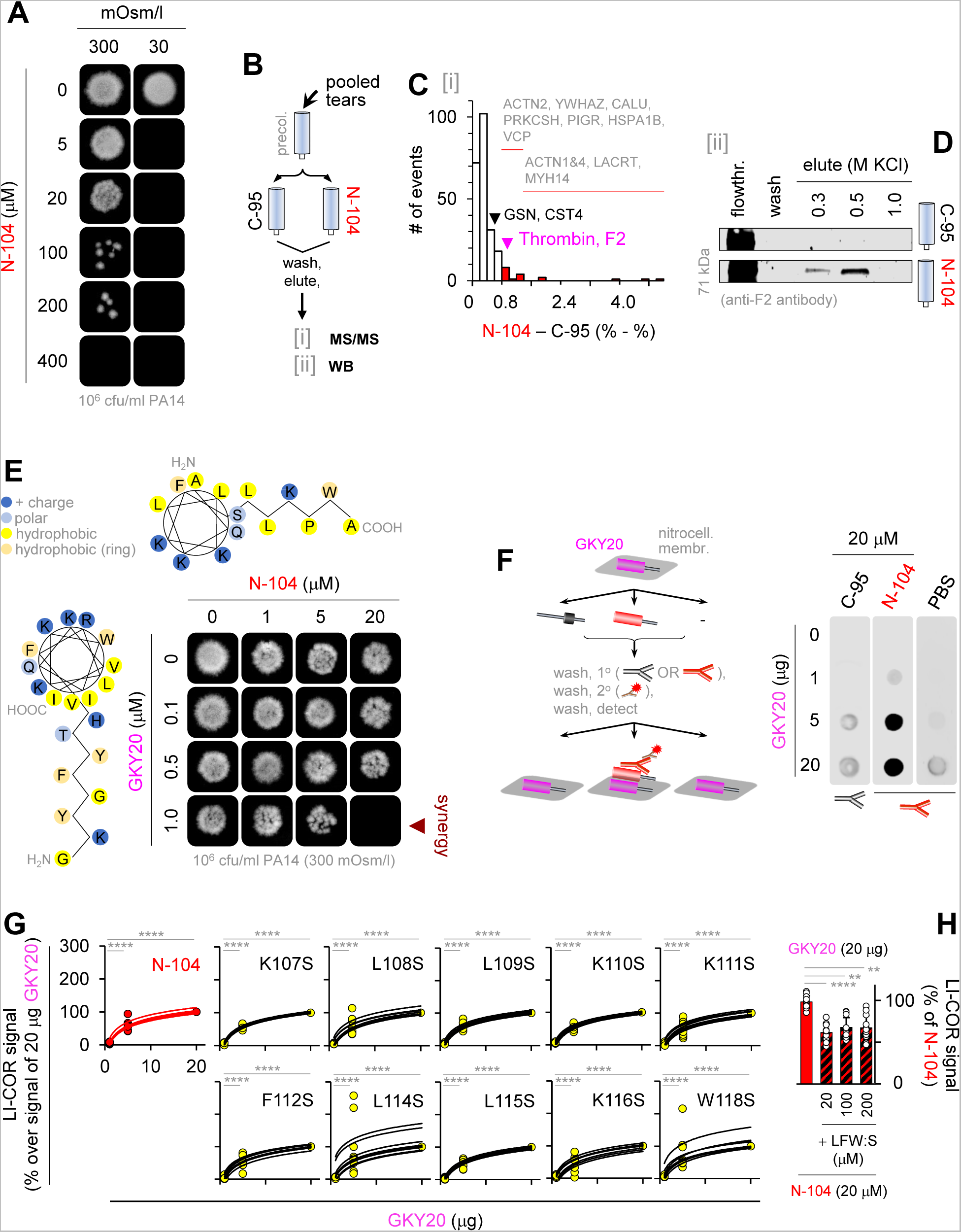
N-104 killing synergizes with thrombin C-terminal ‘GKY20’ peptide. (A) N-104 killing of PA14 at two different osmolalities. (B) N-104 affinity screen of antimicrobial-rich human tears by: (i) tandem mass spectrometry (MS/MS) and (ii) western blotting (WB). (C) Thrombin identified as a N-104 ligand, among others. (n = 3 experiments). (D) Thrombin western blots of human tear flowthrough, final wash fraction or KCl eluants off N-104 or negative control C-95 columns. (E) N-104 plus thrombin peptide GKY20 checkerboard killing assay. Both share an amphipathic α-helix and coiled-coiled secondary structure. (F) N-104 (but not C-95) ligation of immobilized GKY20 as respectively detected by ab-C-term and ab-N-term lacritin antibodies (n = 5 experiments). (G) Affinity of GKY20 for N-104 is similar to that for (Figure 1) N-104 analogs. Two-way ANOVA with Dunnett’s multiple comparisons test (n = 3 experiments). (H) GKY20 ligation of N-104 L108S/L109S/F112S/L114S/L115S/W118S (’LFW:S’) is less. Friedman ANOVA with Dunn’s multiple comparisons test (n = 3 experiments). Data represent the mean ± SD, ****p<0.0001, **p<0.01. See also **Figure S6 and Table S5**.

We next tested seven different clinical isolates of *P. aeruginosa*, as well as one each of *Serratia marcescens* and *Stenotrophomonas maltophilia* - all opportunistic Gram-negative bacteria that are resistant to bacitracin, vancomycin, cefazolin, and sulfasoxazole. Additional resistance is to polymyxin B (*Serratia marcescens*), gentamicin and tobramycin (*Stenotrophomonas maltophilia*), and ofloxacin and moxifloxacin (*P. aeruginosa* K2414). Resistance was determined by the University of Pittsburgh’s Charles T. Campbell Ophthalmic Microbiology Laboratory. In the synergistic 20 µM (N-104):1 µM (GKY20) mixture, growth of all but three (10^3^ cfu/ml) or four (10^5^ cfu/ml) clinical isolates of *P. aeruginosa* and both *S. marcescens* and *S. maltophilia* strains was prevented (**Figures 7A and S7A-S7B**). Hemolytic activity of N-104 (**Figure 1H**) and GKY20 (**Figure 7B**) alone or together up to a 500 µM N-104:25 µM GKY20 mixture was low. We next tested the 20 µM (N-104):1 µM (GKY20) mixture in a well-established mouse model of PA14 keratitis ^66^ in which corneal scratches are introduced in anesthetized mice to facilitate seeding of PA14 four and eight hours before topical treatment with the peptide mixture (**Figure 7C**). Twenty four hours later, we assessed the level of keratitis and of CFU’s. Treatment significantly diminished the level of ocular opacity by ~2 fold (**Figure 7D**) and CFU infectivity by ~2.8 fold (**Figures 7E-7G**), each with a strong effect size estimate (d>1.4). Taken together, N-104 binds thrombin and its C-terminal GKY20 peptide. Also, N-104 with GKY20 act synergistically in their joint capacity to kill multidrug resistant *P. aeruginosa*, *S. marcescens* and *S. maltophilia in vitro* and PA14 in infected mice.

**Figure 7.**
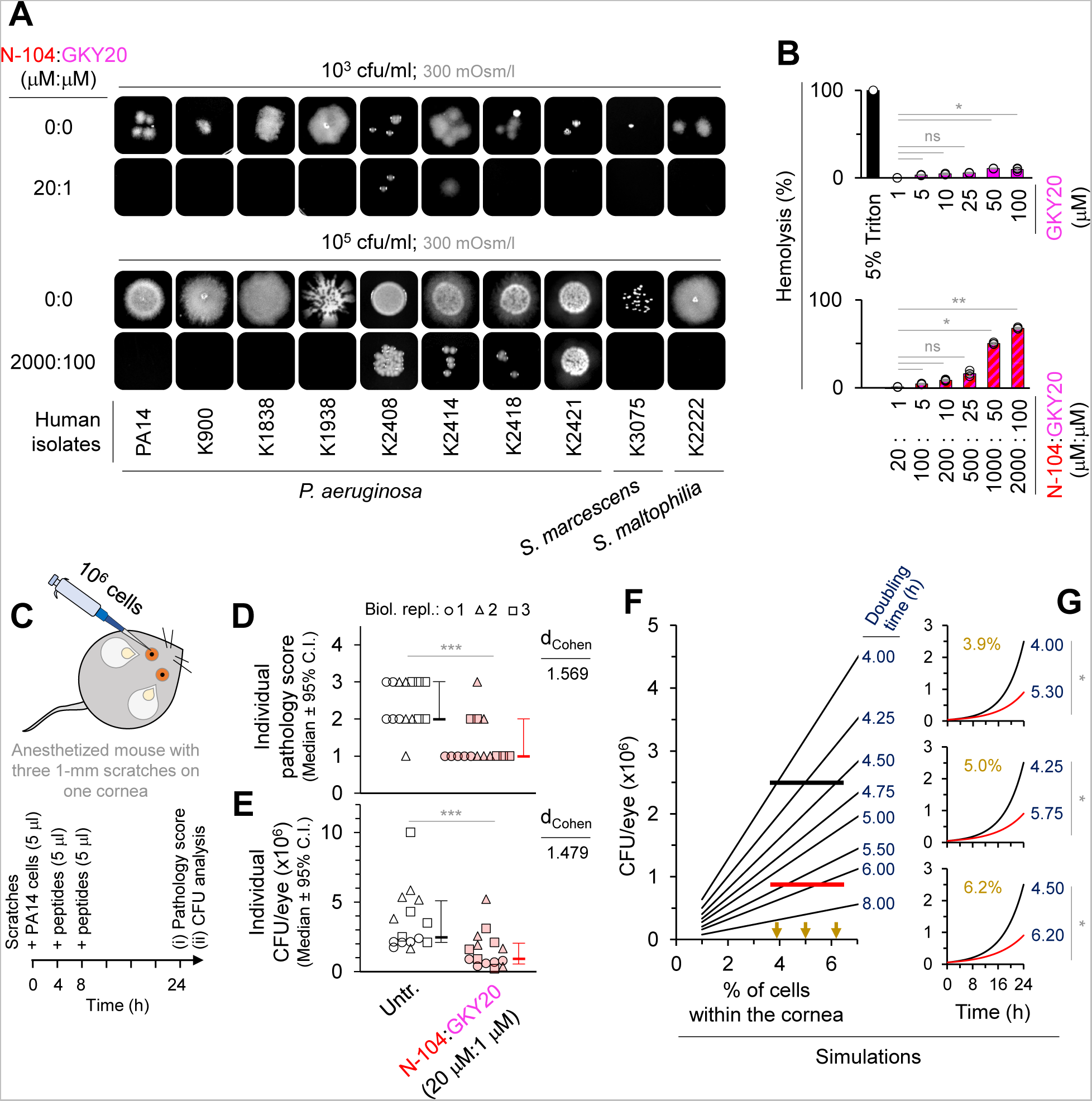
N-104 plus GKY20 kill multidrug resistant clinical isolates as well as PA14 in infected mice. (A) N-104 plus GKY20 killing of clinical isolates. (B) Test of N-104 plus GKY20 and GKY20 alone for hemolysis activity (see Figure 1H for N-104 hemolysis assay). Friedman ANOVA with Dunn’s multiple comparisons test (n = 3 experiments). (C) PA14 infection of mouse cornea after scratching ^66^ without or with N-104 plus GKY20. (D) Corneal surface pathology scores 24 hours after infection without or with N-104 plus GKY20 treatment (0, eye macroscopically identical to the uninfected contralateral control eye; 1, faint opacity partially covering the pupil; 2, dense opacity covering the pupil; 3, dense opacity covering the entire anterior segment; and 4, perforation of the cornea and/or phthisis bulbi). Mann-Whitney test and effect size estimate for non-parametric tests from Psychometrica ‘Computation of Effect Sizes’ (n = 3 experiments). (E) CFU infectivity remaining in the cornea without or with N-104 plus GKY20 treatment. Mann-Whitney test and effect size estimate for non-parametric tests from Psychometrica ‘Computation of Effect Sizes’ (n = 3 experiments). (F) Simulation of reduced CFU infectivity as a consequence of increased PA14 doubling time. (G) Simulated doubling time and CFU infectivity when the density of PA14 is 3.9, 5 or 6.2% (arrows in (F)) of all corneal cells. Unpaired t test (one simulation over 24 time points). All bar graphs represent the mean ± SD. (D) and (E) are represented as the median ± 95% confidence interval. ***p<0.001, **p<0.01, *p<0.05, ns, nonsignificant. See also **Figure S7**.

### N-104-dependent interplay between anaerobic respiration and the acquisition of iron

Although FeoB is the most common Fe^2+^ uptake mechanism ^22^ and multisubunit PotFGHI containing PotH is one of two main polyamine transporters in bacteria ^53^, other Fe^2+^ (for example hypothetical CIA_05411, ABJ14631), and polyamine transporters may be available. There are also elaborate Fe^3+^ acquisition mechanisms involving chelating siderophores, capture by bacterial surface receptors and inner membrane transport by TonB:ExbB:ExbD ^25^. Toward a broader exploration of N-104 triggered changes, RNA-seq was performed (**Figure S8A-SG**). Differential gene expression following 5 or 20 µM N-104 treatment of PA14 versus untreated (**Figure S8A (i),(ii)**) or 20 µM N-104 versus negative control 20 µM N-80/C-25 (**Figure S8A (iii)**) was assessed using established thresholds (adjusted *p*-value of < 0.001; fold change of ≥ 2.83; **Table S6**). Volcano plots revealed no N-104-dependent RNA regulation of YaiW, FeoB and PotH, perhaps in keeping with the proximal role of each as mediators of N-104 activity (**Figure S8A**). We initially focused on N-104 vs N-80/C-25 (**Figures S8B-8C**). 118 genes were significantly downregulated and 44 upregulated, of which the protein products of 72 down- and 22 up-regulated were hypothetical or poorly characterized. Downregulated genes include those coding for proteins involved in: Fe^3+^ transport (*exbB1* codes for ExbB ^25^ (**Figure S9**); FbpA transports Fe^3+^ across the inner membrane ^67^, and both taurine uptake (WP_003114312.1, PA14_12490) and release of sulfite from taurine (TauD) when sulfate is limiting ^68^. TauD is complexed to Fe^2+^ ^69^ and is just below the threshold of our Keio library resistance screen (**Figure 2D**). Like *tauD*, downregulated *ssuF*, *ssuD*, *atsK*, *atsA* and PA14_02435 (**Figure S9**) code for cytoplasmic desulfonating enzymes ^69,70^. Other downregulated categories include protein products necessary for electron transport or another part of the respiratory chain (CoxA, CoxB, EtfA, EtfB, FadH1, PA14_02460, PA14_06640, PA14_25840), fatty acid oxidation (FoaA, FoaB), antioxidants (GpO), methionine import (MetN1, MetQ-I), Mg^2+^ transport (MgtA), molybdate ion transport (ModA), cell adhesion (Pa1L), biofilm formation (PfpL, PA14_34050), transcriptional regulation (PsrA, PA14_27530, PA14_40380), universal stress proteins (UspA, PA14_21220 [UspK], PA14_66460) and small molecule transport (PA14_34770). Among upregulated categories were genes coding for proteins necessary for amino acid transport (AroP2), sulfate transport (CysT; **Figure S9**), glutamine-rich toxic protein (GrtA; inhibits *E. coli* growth ^71^), molybdopterin cofactor biosynthesis (MoeA1, MoaB1), respiratory nitrate reductase (NarG, NarH, NarI, NarJ; **Figure S9**) responsible - together with molybdopterin cofactor - for anaerobic growth of *P. aeruginosa* in cystic fibrosis ^72^, anaerobic regulation of denitrification (NosR; **Figure S9**), multidrug efflux (OprJ), cell wall biosynthesis (PbpA), sulfate binding or metabolism (SbP, SsuE; **Figure S9**), a sulfonate transporter (SsuA), a serine endopeptidase (PA14_24360; **Figure S9**), and a putative beta-lactamase (PA14_41280) that cleaves beta-lactam antibiotics to promote resistance ^73^. We next compared (i-iii) data (**Figure S8C**) to appreciate changes common to both 5 and 20 µM N-104. Three (*exbB1*, WP_003114312.1 and *tauD*; **Figure S9**) were downregulated and three (PA14_24360, PA14_41280, PA14_41290) upregulated. Curious whether any of these as *E. coli* orthologs may interact with YaiW, FeoB and/or PotH, we explored BioGRID (**Table S2**). The only predicted protein interaction was that of FeoB with ExbD (which binds ExbB; **Figure S9**) and YidC (membrane insertase; up-regulated only in (ii)). Predicted genetic interactions include *feoB* with *napA* (nitrate reductase; down-regulated only in (ii)) and *potH* with both *htpG* (heat shock protein; down-regulated only in (iii)) and *rho* (transcription termination factor; up-regulated only in (ii)). Thus, N-104 acts broadly. Perhaps most striking is the shutdown of genes necessary for aerobic respiration pointing to a likely switch from aerobic to anaerobic respiration that favors N-104’s antibiotic activity.

## DISCUSSION

Manipulation of transporters to limit bacterial respiration and thereby pathogenesis, has been a decades long quest towards addressing multi-drug resistance. Here, using unbiased forward genetic screens of single gene knockout strains, protein affinity screens, mutational and synergy analyses and RNAseq, we report lacritin N-104 as the first example of an inhibitor of multiple bacterial transporters through direct binding or gene downregulation that in turn profoundly affects respiratory capacity. By gaining periplasmic access via outer membrane lipoprotein YaiW, the mechanism hinges on N-104 ligation of predicted periplasmic loops 4 and 2 respectively of the inner membrane transporters FeoB (a well-known virulence factor) and PotH (of the PotFGHI complex) to respectively suppress both ferrous iron and polyamine uptake. It also directs the transcriptional shut down of Fe^3+^, taurine, methionine, Mg^2+^, molybdite and small molecule transport machinery, as well as multiple elements of the respiratory chain.

N-104 is one of over 40 different proteoforms from lacritin’s pleiotropic C-terminus ^32^ whose cleavage-dependent bactericidal activity was first identified by progressive recombinant protein and then synthetic peptide screening initially in *E. coli* and culminating with *P. aeruginosa* ^27^. Later validation by ‘Random Forests’, ‘Support Vector Machines’ and ‘Discriminant Analysis’ of human tear proteoforms focused on slightly smaller, but equally active ‘N-106’ and ‘N-107’ on *E. coli*, *P. aeruginosa*, and on Gram-positive *L. monocytogenes* and *S. aureus* ^32^. Other lacritin proteoforms overlap with N-104. These include epithelial ^30,74,75^ and neuronal ^31^ regenerative, autophagic ^29^ and secretory ^28,76–78^ agonists. Some also act as surfactants ^79^. All center on lacritin’s C-terminal amphipathic α-helix that serves as the ligation surface for its co-receptor ^52^ and possibly for outer membrane YaiW. The latter is suggested by diminished bactericidal activity of N-104 L108S, L109S and F112S within the α-helix, and by less N-104 periplasmic entry in the absence of YaiW. N-104 and overlapping proteoforms are thus uniquely capable of both innate defense and subsequent epithelial and neuronal regenerative repair.

Targeting periplasmic constituents or the periplasmic domain of inner membrane proteins is a relatively new strategy to address drug resistance ^80^. Although none suppress transport of iron nor multiple transporters, targets include: (i) AcrA necessary for assembly of the drug efflux complex AcrAB-TolC ^81^, (ii) the virulence-associated thiol:oxidoreductase DsbA necessary for intrachain disulfide bonding in the elaboration of type III and VI secretion systems respectively in *P. aeruginosa* ^82^ and *S. marcescens* ^83^ and in other proteins, (iii) the peptidyl-prolyl isomerase/chaperone SurA required for periplasmic transport and folding of outer membrane proteins/virulence factors although not necessary in a multidrug resistant strain of *A. baumannii* ^84^, (iv) LolA that carries lipoproteins from the inner to outer membrane as part of the LolABCDE system ^85^, (v) inner to outer membrane lipopolysaccharide transport family member LptA ^86^, (vi) periplasmic ZnuA that delivers zinc to the inner membrane ZnuBC transporter ^87^, and (vii) the abundant periplasmic TolB protein of the TolQ-TolR-TolA-TolB-Pal complex together necessary for the joint integrity of inner and outer membranes and peptidoglycan layer, as well as translocation of the colicin family of protein antibiotics ^88^. With 76 and 7 proteins respectively detected in the *P. aeruginosa* periplasm and inner membrane ^89^, the likelihood that AcrA, DsbA, SurA, LolA, LptA, ZnuA or TolB might interact with YaiW, FeoB or PotH is low. This appears to be the case for YaiW and PotH. However, high throughput bait/prey capture in *E. coli* for mass spectrometry detected FeoB complexed with TolB as well as TolQ of the same complex ^90^. TolB and TolQ are widely expressed among *Enterobacteriaceae, Acinetobacter and Pseudomonas* and are regulated by iron respectively late or throughout growth ^91^.

At least nineteen other periplasmic or inner membrane proteins are targeted by FeoB ^90^. Notable are those involved in bacterial respiration including: (i) NuoE, cytoplasmic subunit E of the mitochondrion-like proton pump NADH:quinone oxidoreductase (complex I) ^92^; the periplasmic, inner membrane anchored thiol:disulfide oxidoreductase CcmG (DsbE) thought to be involved in the maturation of cytochrome C ^93^; cytosolic TdcD - a propionate kinase that catalyses propionyl phosphate with ADP to propionate and ATP ^94^; and the NAD(P)H: quinone oxidoreductase WrbA suggests an expanded role of FeoB in bacterial respiration. Coupled with N-104 transcriptional downregulation of ten other electron transport or respiratory chain or fatty acid oxidation elements, the apparent suppression of aerobic respiration and inhibition of polyamine transporter PotH is profound. Indeed, in the anoxic Fe^2+^-rich environment of the earth’s Paleoarchean period, polyamines are thought to have nourished LUCA – the hypothetical last universal common ancestor ^3^ from which bacteria, archaea and eukarya evolved. N-104 thus in effect reverses evolutionary time to an anaerobic Fe^2+^- and polyamine-dependent era.

## METHODS

Bacterial strains, clinical isolates, plasmids, primers, recombinant proteins, synthetic peptides, chemicals, antibodies, software and algorithms, and deposited data. See **Key Resources Table**.

## EXPERIMENTAL MODEL AND STUDY PARTICIPANT DETAILS

### Cells

*E. coli* K12 (GC616) and the Keio collection of *E. coli* K12 single gene knockouts (JW1023-KC) ^42^ were purchased from the National Institute of Genetics’ National BioRource Project (Mishima, Japan). Competent *E. coli* BL21 (DE3; C2527l) was purchased from New England BioLabs (Ipswich, MA). Sheep red blood cells (0855876) were purchased from MP Biomedicals (Irvine CA). Seven different clinical isolates of *P. aeruginosa*, as well as one each of *Serratia marcescens* and *Stenotrophomonas maltophilia* were provided by Robert Shanks of the University of Pittsburgh’s Charles T. Campbell Ophthalmic Microbiology Laboratory. *E. coli* OP50 and *C. elegans* N2 were purchased from *Caenorhabditis* Genetics Center (CGC), University of Minnesota. *P. aeruginosa* PA14 and PA14 *feoB*, *potH (spuG)* and *yaiW* transposon mutants were provided by Mihaela Gadjeva from Harvard University’s PA14 Transposon Insertion Mutant Library ^43^. *E. coli* and *P. aeruginosa* were expanded in LB broth. *C. elegans* L1 stage worms were cultivated at room temperature on Nematode Growth Medium (NGM) plates, with *E. coli* OP50 as food source, until they matured into sterile young adults.

Sample size estimation: Sample sizes were not chosen based on a pre-specified effect size, thus sample sizes were not estimated at the outset. Rather, multiple independent, biological replicate experiments were performed as specified in the figure legends.

### Synthetic peptide design

We ^27^ and others ^32^ had previously narrowed lacritin’s (LACRT) cleavage potentiated C-terminal bactericidal activity to ‘N-104’ (AQKLLKKFSLLKPWA; numbering indicates a synthetic peptide lacking 104 N-terminal lacritin amino acids excluding the signal peptide). Lacritin ‘N-80/C-25’ (lacking 80 N-terminal amino acids without signal peptide and 25 C-terminal amino acids; AKAGKGMHGGVPGG) is inactive ^27^. As an additional negative control, we tested ‘C-95’ (lacking 95 C-terminal amino acids; EDASSDSTGADPAQEAGTSKPNEE). These and other peptides were synthesized as a trifluoracetic acid (TFA) salt (≥ 95% purity; HPLC) by GenScript (Piscataway NJ) with N-terminal acetylation (N-104, N-80/C-25) and C-terminal amidation (N-80/C-25, C-95). In *S. aureus*, several TFA peptides were superior to the same peptides as acetate or chloride salts with less hemolysis ^96^. To assay the respective contribution of amino acids with nonpolar or basic side chains, a panel of N-104 analogs was generated by substituting serine for a single leucine (‘L108S’, ‘L109S’, ‘L114S’, ‘L115S’), lysine (‘K107S’, ‘K110S’, ‘K111S’, ‘K116S’), phenylalanine (‘F112S’), or tryptophan (‘W118S’).

For imaging flow cytometry of N-104 (vs negative control C-95) entry into PA14 cells, we respectively synthesized ‘FITC-N-104’ and ‘FITC-C-95’ with N-terminal added FITC. Similarly, for higher resolution wide-field fluorescence of the same process, we synthesized Cys(N)-N-104 and Cys(N)-C-95 with N-terminal added cysteine to which was conjugated Janelia Fluor 549 maleimide. Conjugation was initiated in 50 mM Tris (pH 8.5), 5 mM EDTA at a 10:1 fluorophore to peptide molar ratio for 3 h at room temperature in the dark. Separation of conjugated peptides from free fluorophore was by Sephadex^TM^ G-10 size exclusion chromatography followed by MALDI-TOF analysis of the conjugate fraction. Validation of these experiments took advantage of the Bacterial ChloroAlkane Penetration Assay for which chloroalkane tagged N-104 and C-95 were synthesized as previously described ^50^.

To screen for proteins in PA14 lysates and later human tear proteins with affinity for N-104, Cys(N)-N-104 (or negative control Cys(N)-C-95) was immobilized on SulfoLink® Coupling resin. Bactericidal peptides from identified tear N-104 binding proteins were then synthesized including thrombin (F2) C-terminal peptide GKY20 (GKYGFYTHVFRLKKWIQKVI), gelsolin (GSN) N-terminal peptide QRLFQVKGRR, and N-terminal cystatin S peptide (CST4) SSSKEENRIIPGGI.

Guided by ‘TMHMM v 2’ and ‘DeepTMHMM 1.0.24’, we synthesized peptides corresponding to FeoB’s predicted outer loop1 (INIGGALQP), 2 (LEDSGYMARAAFVMDRLMQ), 3 (GAFFGQGGA) and 5 (ATFAA), as well as PotH’s predicted outer loop 2 (WMGILKNNGVLNNFLLWLGVIDQPLTILHTN) and 3 (ELLGGPDSIMIGRVLWQEFFNNRDW). Also synthesized was syndecan-1 pep19-30 (DNFSGSGAGAL) ^52^. All were acetylated and amidated, respectively on amino and carboxy termini. HPLC analyses confirmed the purity (≥ 95%) of all synthetic peptides (GenScript, Piscataway NJ), which were aliquotted and stored lyophilized at −80°C in a dry environment.

### Surface Plasmon Resonance (SPR)

A lipid bilayer supported on lipophilic SPR sensor chip L1 (BR100558; GE Healthcare Life Sciences) was prepared using a 59:21:20 mixture of 1-palmitoyl-2-oleoyl-sn-glycero-3-phosphocholine:1-palmitoyl-2-oleoyl-sn-glycero-3-phospho-(1’-rac-glycerol):1-palmitoyl-2-oleoyl-sn-glycero-3-phosphoethanolamine (PC:PG:PE) with some similarity to the outer Gram-negative bacterial membrane although lacking lipopolysaccharide ^97^. Another lipid bilayer was formed from PC:PG (75:25) as a model for Gram-positive bacteria ^98^. The lipid bilayers were formed in 8.3 mM phosphate buffer and 10 mM NaCl (pH 7.2). N-104 or N-104 analogs were then introduced to a Biocore 3000 (Cytiva, Marlborough MA) and their time-resolved SPR responses were recorded at a flow rate of 30 μl/min for 100 s, followed by a buffer wash at 30 μl/min for 500 s. The use of a high flow rate minimized mass transfer limitations and resulted in good agreement among the four replicates of each experiment. The experiment was terminated with the addition of 20 mM 3-[(Cholamidopropyl) dimethylammonio]-1-propanesulfonate (CHAPS) to remove lipids and peptides from the sensor chip for reuse. CHAPS was introduced at a flow rate of 5 μl/min for 1 min.

### Quartz Crystal Microbalance with Dissipation Monitoring (QCM-D)

The same two supported lipid bilayers on a silicon dioxide sensor chip (QSX 303; NanoScience Instruments, Phoenix AZ) in a QSense Explorer (Biolin Scientific, Stockholm Sweden) exhibited characteristic negative frequency shifts (ΔF = −25 Hz) indicative of positive mass as well as a dissipation shift (ΔD = (0.1 - 0.2) × 10^−6^) denoting a rigid layer. N-104 or N-104 analogs were introduced for 10 minutes at a flow rate of 100 μl/min, followed by a 10 min buffer wash. The changes in mass and rigidity (viscoelasticity) of the lipid/peptide layers were determined by analyzing the 5th, 7th, and 9th overtones and calculating the ΔΔF and ΔΔD, respectively.

### Colony forming unit (CFU) assay

100 μl aliquots of wild type *E. coli* or *E. coli* gene deletion strain, or wild type or transposon mutant PA14 overnight cultures were suspended in 1 mM Na_2_HPO_4_, 0.18 mM KH_2_PO_4_, 0.27 mM KCl, 13.7 mM NaCl, pH 7.2 or in 10 mM Na_2_HPO_4_, 1.8 mM KH_2_PO_4_, 2.7 mM KCl, 137 mM NaCl, pH 7.2, at a final density of 10^6^ cfu/ml. Each was treated with N-104, N-80/C-25, C-95, N-104 analogs, N-104 together with thrombin peptide GKY20, or left untreated for ~8 hours at 35°C (the temperature at the eye surface), and then was used to each (2 µl) inoculate an Luria-Bertani agar plate with or without relevant antibiotic supplementation. After overnight incubation at 37°C, plates were analyzed on a ChemiDoc XRS (Bio-Rad, Hercules CA).

### *C. elegans* – PA14 liquid infection assay

L1 stage worms on Nematode Growth Medium (NGM) plates with *E. coli* OP50 as food source were cultivated at room temperature in a humidified chamber until maturation into sterile young adults ^99^, carefully washed off with M9 minimal salts and then collected by pelleting (1200×g, 2 min). ~50 worms per well in sterile black 96-well plates containing 150 μl of 90% M9 minimal salts and 10% LB broth were then incubated aerobically with an equal volume of cultures of PA14 or of PA14 transposon mutants of *feoB* (PA14_56680), *PotH* (PA14_03950), or *yaiW* (PA14_29270), or alternatively with *E. coli* OP50 as negative control (diluted to 10^7^ cfu/ml in 1 mM Na_2_HPO_4_, 0.18 mM KH_2_PO_4_, 0.27 mM KCl, 13.7 mM NaCl, pH 7.2) that had been previously treated with N-104 or N-104 analogs (50 µM of each) for 5 h at 35°C. After 3 days, bacteria and debris were removed leaving 50 μl per well of worms to which was added 50 μl of 6 μM SYTOX Orange in M9 minimal salts. The next day, images were captured using both transmitted and epi-fluorescence (excitation/detection = 535/610 nm) on an AMG EVOS FL AMF-4301 fluorescence microscope (Thermo Fisher Scientific, Waltham MA).

### Nuclear magnetic resonance and circular dichroism analyses

Two N-104 samples were dissolved in 10 mM Na_2_HPO_4_, 1.8 mM KH_2_PO_4_, 2.7 mM KCl, 137 mM NaCl, pH 7.2 with or without the addition of 150 mM dodecylphosphocholine-d_38_ (DPC-d_38_), respectively. NMR spectra were collected at 25°C at an N-104 concentration of 1 mM. For both samples, ^1^H-^1^H TOCSY and ^1^H-^1^H NOESY spectra were collected at various mixing times to achieve the best resolved spectra for spectral assignments. The chemical shifts showed that N-104 in buffer without dodecylphosphocholine-d_38_ exists only in a random coil structure. In contrast, in dodecylphosphocholine-d_38_ N-104 showed characteristic helical chemical shifts so that we were able to walk through the resolved linkages in the spectra to assign sequence specific chemical shifts. The combination of ^1^H-^1^H TOCSY with a mixing time of 60 ms and ^1^H-^1^H NOESY with a mixing time of 80 ms resulted in the proton chemical shifts as assigned in **Figures S1J-S1K**. Subsequently, ^13^C-^1^H and ^15^N-^1^H HSQC spectra were collected by leveraging the natural abundant ^13^C and ^15^N isotopes in the peptide samples. The near complete backbone chemical shifts (**Figures S1J-S1K**) allowed us to use TALOS+ ^100^ to predict the existence of helical structure from residue 1 to 10 in N-104 (**Figure 1G**). For validation of NMR, N-104 in 10 mM Na_2_HPO_4_, 1.8 mM KH_2_PO_4_, 2.7 mM KCl, 137 mM NaCl, pH 7.2 without or with 10 mM dodecylphosphocholine was assayed by circular dichroism in a J-1500 Circular Dichoism Spectropolarimeter (JASCO, Easton MD) over a 190-240 nm range.

### Hemolysis assay

The pellet resulting from 300×g (5 min) centrifugation of 500 µl of stock sheep red blood cells (55876; MP Biomedicals) was resuspended in an equal volume of 10 mM Na_2_HPO_4_, 1.8 mM KH_2_PO_4_, 2.7 mM KCl, 137 mM NaCl, pH 7.2, of which 50 μl aliquots were treated at 37°C for 2 h with 1, 25, 50, or 100 μM of N-104 or GKY20 or both (N-104:GKY20 [µM each]: 20:1, 100:5, 200:10, 500:25, 1000:50, 2000:100) or as positive control with 5% Triton X-100 in a final volume of 100 μl. After centrifugation at 300×g for 5 min, the supernatant absorbance at 540 nm of the supernatant minus the absorbance of the untreated sample was recorded and normalized versus the Triton X-100 sample.

### Forward genetic screens of Keio *E. coli K12* single gene knockout collection

Advantage was taken of the Alamar Blue viability assay ^101^. Briefly, wells of deep 96-well plates (1896-2000, USA Scientific) each containing 0.8 ml of LB broth with 50 μg/ml of kanamycin (BP906-5, Fisher Scientific) were singly innoculated with 10 μl aliquots of each of 3,884 knockout strains for 16 hours at 37°C on a rotator (160 rpm). 200 µl of a 1:1000 dilution of each (~10^6^ cfu/ml) in 1 mM Na_2_HPO_4_, 0.18 mM KH_2_PO_4_, 0.27 mM KCl, 13.7 mM NaCl, pH 7.2 containing 10% Alamar Blue (DAL1100, Invitrogen) in wells of black 96-well plates were monitored hourly for five hours in a SpectraMax M3 plate reader (Molecular Devices; San Jose, CA) with 3 s agitation before reading simultaneously from the well bottoms at medium PMT voltage with a 590 nm cutoff filter and six flashes per read for the Alamar Blue fluorescence signal (excitation/detection = 560/590 nm). The same hourly monitoring continued for 30 hours after the addition of N-104, N-80/C-25 (negative control), or ampicillin (positive control; BP1760-5; Fisher Scientific) at a final concentration each of 100 µM. Data for each of 3,884 strains were normalized to untreated with 10% Alamar Blue background subtracted as the ratio of the viability slope during the first five hours of N-104 treatment versus that five hours previous.

### Imaging flow cytometry

0.5 ml of 10^6^ cfu/ml PA14 cells in 1 mM Na_2_HPO_4_, 0.18 mM KH_2_PO_4_, 0.27 mM KCl, 13.7 mM NaCl, pH 7.2 were incubated aerobically for one hour at 35°C with 10 or 100 μM FITC-N-104 (without, or as positive control with 70% ethanol) or with negative control FITC-C-95. Cells were subsequently washed in the same buffer, pelleted, treated with 100 μl of 0.25% trypsin EDTA (Gibco, 25200-056) for 5 min at 35°C to remove surface attached peptide, further washed and pelleted, resuspended in 100 μl in the same buffer for fixation with an equal volume of 8% formaldehyde (18814, Polysciences Inc.) and then imaged using an Amnis Imagestream X Mark II imaging flow cytometer. 500-1000 counts were analyzed per replicate. To ensure the exclusion of debris and out-of-focus cells, ‘Gradient-RMS’ (’average slope spanning three pixels in an image’; 0-100) versus Area (’number of pixels in an image reported in square microns; 1-200) of channel 1 was plotted for each experiment with over 85% of the data falling within the ‘cells-in-focus’ gate. The fluorescence intensity values of FITC-positive cells (Intensity-MC, channel 2) were analyzed using the IDEAS version 6.2 program (Amnis, Seattle WA) and plotted as histograms with a bin number of 100. FITC-positive cells were selected based on histograms generated from the positive control sample (≥ 0.002).

### Wide-field fluorescence imaging

Janelia Fluor 549-N-104 and -C-95 (each 25 µg) in 20 µl of M9 minimal salts was incubated with ~3.75×10^9^ cfu of pelleted PA14 cells for 1 hour at 35°C, washed four times with filtered 10 mM Na_2_HPO_4_, 1.8 mM KH_2_PO_4_, 2.7 mM KCl, 137 mM NaCl, pH 7.2, and then treated with 5 µM SYTOX green to assess membrane integrity. Stained cells (1 µl) were immobilized on cover slip supported 1.5% agarose pads for imaging on a custom-built inverted fluorescence microscope, using 1 W/cm^2^ of 561-nm laser light (Genesis MX561 MTM, Coherent) to excite Janelia Fluor 549 Maleimide and 4 mW/cm^2^ of 488-nm laser light (Coherent) to excite SYTOX green. Emitted light collected using an oil immersion objective (PLSAPO, 60×, 1.4 NA) was passed through a filter set (zt440/488/561rpc, Chroma Technology) to block excitation light with separation of Janelia Fluor 549 Maleimide and SYTOX green emission using a diochroic beam splitter (T5g0lpxr-uf3, Chroma Technology) as collected on separate sCMOS cameras (ORCA-Flash 4.0 V2, Hamamatsu Photonics). Brightfield images were collected to obtain a total count of cells in each field of view.

### Bacterial ChloroAlkane Penetration Assay (BaCAPA)

The Bacterial ChloroAlkane Penetration Assay was performed as previously described ^50^. Briefly, overnight cultures of *E. coli* Lemo21 (DE3) transfected with pET-21 b (+) containing the cyto_Halo or peri_Halo cDNA insert (respectively for cytoplasm versus periplasm expression) were induced for 2 hours with 1 mM isopropyl β-D-1-thiogalactopyranoside. Cultures were pelleted, washed three times with 1 mM Na_2_HPO_4_, 0.18 mM KH_2_PO_4_, 0.27 mM KCl, 13.7 mM NaCl, pH 7.2 and then resuspended in the same buffer at a density of ~4×10^9^ cfu/ml. Subsequently, 50 µl aliquots cells were treated with 0, 4, 8, 16, 32 or 64 µM chloroalkane tagged N-104 (or C-95) for 30 minutes at 37°C with agitation, washed with 1 mM Na_2_HPO_4_, 0.18 mM KH_2_PO_4_, 0.27 mM KCl, 13.7 mM NaCl, pH 7.2, exposed to 5 μM of rhodamine chloroalkane (G3221, Promega) for 30 minutes at 37°C with shaking, washed with 1 mM Na_2_HPO_4_, 0.18 mM KH_2_PO_4_, 0.27 mM KCl, 13.7 mM NaCl, pH 7.2, fixed for 30 minutes at room temperature with 2% formaldehyde solution, and then subjected to flow cytometry (Attune NxT Acoustic Focusing Cytometer, Invitrogen) with blue laser excitation (530 nm).

### N-104 affinity purification

For generation of N-104 and negative control C-95 affinity columns, 2 mg of Cys(N)-N-104 or Cys(N)-C-95 in 2 ml of 50 mM Tris (pH 8.5), 5 mM EDTA coupling buffer were incubated with an equal volume of suspended SulfoLink® Coupling Resin (20401, Thermo Scientific) with mixing for 3 hours at room temperature. Columns were subsequently washed with 3 ml of coupling buffer followed by 2 ml of freshly prepared 50 mM L-cysteine (A10435, Alfa Aesar) for 2 hours at room temperature for quenching and then washed with 10 ml of 1 M NaCl prior to use or storage in 0.05% sodium azide at 4°C. Coupling efficiency was determined using fluorescence from N-104’s penultimate C-terminal tryptophan as 0.65 ± 0.04 μmole per ml of resin.

Screening for PA14 proteins with N-104 affinity columns was initiated from pelleted 2.5 liter overnight cultures that were washed with ice-cold 10 mM Na_2_HPO_4_, 1.8 mM KH_2_PO_4_, 2.7 mM KCl, 137 mM NaCl, pH 7.2 and lysed by sonication in 25 ml of 50 mM Tris (pH 7.5), 1 mM MgCl_2_, 200 mM *n*-octyl-β-D-glucopyranoside (O8001, Millipore Sigma) with protease inhibitors (11836153001, Roche). Sonication involved five cycles of 15 s with 45 s rest on ice followed by vortexing on ice for 30 min before being rocked end-over-end overnight at 4°C. After centrifugation, the supernatant was subjected to end-over-end mixing with a 3-ml pre-column of Pierce Agarose (26150, Fisher Scientific) followed by equal division by protein concentration for incubation end-over-end with 1 ml N-104 or C-95 columns at 4°C. Columns were washed with 25 ml of 50 mM Tris (pH 7.5), 50 mM *n*-octyl-β-D-glucopyranoside, 20 mM KCl containing the same protease inhibitors, followed by elution using KCl (50, 75, 100, 125, 150, 300, 500 and 1000 mM) in the same buffer. The 500 mM KCl eluate was submitted for mass spectrometric sequencing. Mass spectrometric sequencing was performed by UVA’s Biomolecular Analysis Facility. Briefly, eluted samples reduced in 0.1 M NH_4_HCO_3_ with 10 mM dithiolthreitol (BP172-25, Fisher Scientific) for 30 min at room temperature were alkylated with 50 mM iodoacetamide in the same buffer (30 min, room temperature), digested with magnetically immobilized trypsin overnight at 37°C, and then desalted and concentrated on reverse-phase C18 tips. Samples were analyzed with a Thermo Orbitrap Exploris 480 mass spectrometer. In total, approximately 25,000 MS/MS spectra were generated, with ions ranging in abundance over several orders of magnitude. The acquired data was analyzed by database searching using the Sequest search algorithm against Uniprot *P. aeruginosa* PA14.

N-104 (vs C-95) affinity purification also explored whether there was affinity for recombinant FeoB and PotH, and for other tear bactericidal proteins (see below).

### FeoB and PotH recombinant expression and purification

Advantage was taken of the pFeoB plasmid ^24^ with *feoB* insert from *P. aeruginosa* PAO1 in pET41-a(+) with C-terminal 8-His tag and kanamycin-resistant cassette, and *E. coli* EcCD00397460 plasmid (DNASU Plasmid Repository, Tempe Arizona) with *potH* insert in PCDF Bravo with C-terminal 10-His tag and streptomycin-resistant cassette. Plasmids of each were generated using standard procedures. From FeoB each ‘TMHMM v 2’ predicted outer loop 1 (INIGGALQP), 2 (LEDSGYMARAAFVMDRLMQ), 3 (GAFFGQGGA), 4 (131 residues) and 5 (ATFAA) were individually deleted using site-directed mutagenesis and, outer loop 4 was substituted for outer loops 2, 3 or 5. Similarly for PotH, ‘TMHMM v 2’ predicted outer loops 1 (FKISLAEMARAIPPYTELMEWADGQLSITLNLGNFLQLTDDPLYFDAYLQSLQ), 2 (WMGILKNNGVLNNFLLWLGVIDQPLTILHTN) and 3 (ELLGGPDSIMIGRVLWQEFFNNRDW) were individually deleted, and outer loop 2 was replaced with 3. Changes were validated by sequencing (Eurofins Genomics LLC). Cells overexpressing wild type and mutated constructs were washed with ice-cold 10 mM Na_2_HPO_4_, 1.8 mM KH_2_PO_4_, 2.7 mM KCl, 137 mM NaCl, pH 7.2 and lysed by sonication in 20% glycerol, 0.5 M NaCl, 10 mM MgSO_4_, 10 mM imidazole, 2% Dodecyl-β-D-maltopyranoside ^102^ (J66869, Alfa Aesar) with protease inhibitors (11836153001, Roche). Sonication involved five cycles of 15 s with 45 s rest on ice followed by vortexing on ice for 30 min before being rocked end-over-end overnight at 4°C. His-tagged FeoB or PotH, wild type or mutated proteins in the supernatant were captured on Ni-Sepharose (1 ml; 17-3712-01, GE Healthcare) for 4 hours with end-over-end rotation at 4°C, washed with 10 mM HEPES potassium salt, 10% glycerol, 0.5 M NaCl, 10 mM MgSO_4_, 10 mM imidazole, and 0.05% dodecyl-β-D-maltopyranoside and eluted with the same wash buffer but containing 0.2 M NaCl and 0.5 M imidazole. Eluted FeoB, PotH and mutated proteins were concentrated into 10 mM HEPES potassium salt, 10% glycerol, 10 mM MgSO_4_, 0.05% dodecyl-β-D-maltopyranoside, and 0.15 M NaCl respectively on Amicon Ultra-0.5 Centrifugal UFC505024 (Amicon), UFC503024 (Amicon) and UFC900596 (Amicon) filter units.

### FeoB and PotH ligation of N-104

N-104 and C-95 columns (1 ml each) equilibrated in 10 mM HEPES potassium salt, 10% glycerol, 10 mM MgSO_4_, 0.05% dodecyl-β-d-maltopyranoside, and 0.15 M NaCl were individually mixed at 4°C overnight with purified FeoB, PotH and mutated proteins. After collecting the flowthrough, columns were washed with 25 ml of 10 mM HEPES, 0.05% dodecyl-β-D-maltopyranoside, and 0.15 M KCl, and were subjected to stepwise elution using 1 ml of the same buffer containing progressively increasing concentrations of KCl (0.3, 0.5, and 1.0 M). Equal volumes of the flow through, final wash fraction and KCl elution fractions were then denatured in 4x Laemmli buffer containing 10% β-mercaptoethanol at room temperature for 15 - 30 minutes without boiling to avoid aggregation, separated by SDS-PAGE on 4 - 20% gradient gels (4561094, Bio-Rad), transferred to nitrocellulose (10600001, GE Healthcare Life Sciences), blocked with 1% fish gelatin (G7765, Sigma) in 10 mM Na_2_HPO_4_, 1.8 mM KH_2_PO_4_, 2.7 mM KCl, 137 mM NaCl, pH 7.2 supplemented with 0.1% Tween-20 (1706531, Bio-Rad), and incubated overnight (4°C) with 10 ml of 1 μg/ml of rabbit anti-His tag (Cell Signaling, 2365S). After three washes in 10 mM Na_2_HPO_4_, 1.8 mM KH_2_PO_4_, 2.7 mM KCl, 137 mM NaCl, pH 7.2 supplemented with 0.1% Tween-20, blots were incubated with 0.1 µg/ml of IRDye 680RD labeled donkey antirabbit IgG (926-68073, LI-COR) at room temperature for 75 minutes. Primary and secondary antibodies were diluted in 10 mM Na_2_HPO_4_, 1.8 mM KH_2_PO_4_, 2.7 mM KCl, 137 mM NaCl, pH 7.2 containing 0.1% Tween-20 and 0.05% sodium azide, with LI-COR Odyssey CLx detection.

In other experiments, 1 ml of ~160 and 80 µg respectively each of FeoB and PotH incubated with 200 µl N-104 column aliquots (final FeoB and PotH concentration of ~2.5 µM) were subjected to increasing concentrations (0, 1, 10, 100 or 250 µM) of FeoB outer loop 2 or 5 synthetic peptides, SDC1 synthetic peptide 19-30, or PotH outer loop synthetic peptides 2 or 3 with end-over-end rocking overnight at 4°C. Different solvents were required for dissolution. DMSO was the solvent for FeoB outer loop 2, and PotH outer loop 2 and 3. Water was the solvent for FeoB outer loop 5 and SDC1 (19-30). After flowthrough collection, columns were washed with 5 ml of 10 mM HEPES, 0.05% dodecyl-β-d-maltopyranoside, 0.15 M KCl, and then were subjected to elution with 200 μl of the same buffer containing 1.0 M KCl. 1.0 M elutions were examined separated by SDS-PAGE and Western blotted as per above.

### Iron transport assay

Iron depletion was induced in PA14 cells (5×10^9^ cfu in 10 ml 10 mM Na_2_HPO_4_, 1.8 mM KH_2_PO_4_, 2.7 mM KCl, 137 mM NaCl, pH 7.2) by treatment with the chelator 1 mM 2,2’-bipyridyl (D216305, Sigma) at room temperature for an hour. After washing twice with 0.9% NaCl, cells were suspended in 1 ml of 1 mM Na_2_HPO_4_, 0.18 mM KH_2_PO_4_, 0.27 mM KCl, 13.7 mM NaCl, for treatment without or with 10, 25 or 100 μM of N-104 (or negative control N-80/C-25 or C-95) for 45 min at 35°C, incubated at the same temperature for 15 min with 100 μM FeSO_4_ (215422, Sigma) and 13 mM ascorbic acid, washed twice with 1 mM Na_2_HPO_4_, 0.18 mM KH_2_PO_4_, 0.27 mM KCl, 13.7 mM NaCl and pelleted at 12,000xg. The ferrozine assay ^55^ was subsequently employed. Briefly, cells in pellets were lysed with 20 μl of 50 mM NaOH, partially neutralized with 20 μl of 10 mM HCl, subjected to iron release with 20 μl of acidified 0.29 M KMnO_4_ for 30 minutes at 80°C, cooled to room temperature, incubated with 7 μl of 30 mM ferrozine (160601, Sigma), 1.1 μl of 180 mM neocuproine (N1501, Sigma), 15 μl of 5 M ammonium acetate (A639, Fisher Scientific) and 15 μl of 1 M ascorbic acid for 30 minutes at room temperature, and then centrifuged at 21,000xg for 10 min. Cellular iron uptake could then be assessed from the supernatant OD_565_. Single molecule transportation per cell was estimated using a standard curve.

### Polyamine transport assay

100 μl aliquots (0.8×10^8^ cfu) of PA14 cell cultures in 1 mM Na_2_HPO_4_, 0.18 mM KH_2_PO_4_, 0.27 mM KCl, 13.7 mM NaCl without or with 10 or 25 µM N-104 (or N-80/C-25 or C-95) for 45 min at 35°C were incubated with 1 µl of 10 mM [1,4-^14^C]putrescine (0.1 mCi/ml; ARC-0245-250, American Radiolabeled Chemicals) or [1,4-^14^C]spermidine (0.1 mCi/ml; ARC-3138-50, American Radiolabeled Chemicals) for 15 minutes at 35°C. Cell pellets were washed four times with 100 μl of 10 mM Na_2_HPO_4_, 1.8 mM KH_2_PO_4_, 2.7 mM KCl, 137 mM NaCl, pH 7.2 and then transferred to 4 ml of scintillation fluid (1200.437, PerkinElmer OptiPhase HiSafe 3) for analysis in a Beckman Coulter LS6500. Also assessed was radioactivity from the last wash. Single molecule transportation per cell was estimated using standard curves for each radiolabeled polyamine.

### Screen for proteins or peptides in human tears that are synergistic with N-104

Human basal tears collected for 5 minutes onto filter paper-like Schirmer strips after 0.5% proparacaine anesthesia and that wicked for at least 12 mm, were subjected to 4°C elution with 10 mM Na_2_HPO_4_, 1.8 mM KH_2_PO_4_, 2.7 mM KCl, 137 mM NaCl, pH 7.2 for a total pooled tear volume of 170 µl. The pool was diluted five-fold in 10 mM Na_2_HPO_4_, 1.8 mM KH_2_PO_4_, 2.7 mM KCl, 137 mM NaCl, pH 7.2 with protease inhibitors (11836153001, Roche) and subjected to end-over-end mixing with a 100 µl pre-column of Pierce Agarose (26150, Fisher Scientific) followed by equal division by protein concentration with each of 100 µl of N-104 or C-95 columns. Incubation with mixing was for 5 hours at (4°C). Columns were subject to washing with 10 mM Na_2_HPO_4_, 1.8 mM KH_2_PO_4_, 2.7 mM KCl, 137 mM NaCl, pH 7.2 containing the same protease inhibitors followed by elution using KCl (0.3, 0.5, and 1.0 M KCl) in the same buffer. The 0.5 M KCl eluate was submitted for mass spectrometric sequencing by UVA’s Biomolecular Analysis Facility, as previously noted. The acquired data was analyzed by database searching using the Sequest search algorithm against Uniprot *H. sapiens*. Anti-thrombin western blotting of non-boiled flowthrough, final wash fraction and KCl elution fractions in 4x Laemmli buffer containing 10% β-mercaptoethanol was performed after separation on 4 - 20% gradient SDS-PAGE gels (4561094, Bio-Rad), transfer onto nitrocellulose (10600001, GE Healthcare Life Sciences), blocking with 1% fish gelatin (G7765, Sigma) in 10 mM Na_2_HPO_4_, 1.8 mM KH_2_PO_4_, 2.7 mM KCl, 137 mM NaCl, pH 7.2 supplemented with 0.1% Tween-20 (1706531, Bio-Rad), and then incubation overnight (4°C) with 10 ml of 1 μg/ml of polyclonal rabbit anti-coagulation factor II/thrombin antibody (NBP1-58268, Novus Biologicals; antigen comprises amino acids 395 - 445) in 10 mM Na_2_HPO_4_, 1.8 mM KH_2_PO_4_, 2.7 mM KCl, 137 mM NaCl, pH 7.2 supplemented with 0.1% Tween-20 (1706531, Bio-Rad) and 0.05% sodium azide. After three washes in 10 mM Na_2_HPO_4_, 1.8 mM KH_2_PO_4_, 2.7 mM KCl, 137 mM NaCl, pH 7.2 supplemented with 0.1% Tween-20, blots were incubated with 0.1 µg/ml of IRDye 680RD labeled donkey antirabbit IgG (926-68073, LI-COR) at room temperature for 75 minutes. Primary and secondary antibodies were diluted in 10 mM Na_2_HPO_4_, 1.8 mM KH_2_PO_4_, 2.7 mM KCl, 137 mM NaCl, pH 7.2 containing 0.1% Tween-20 and 0.05% sodium azide. After washing in the same way, imaging was on a LI-COR Odyssey CLx.

Synthesized peptides (see Synthetic peptide design) corresponding to identified and selected N-104 binding proteins were subjected to modified PA14 checkerboard killing assays with N-104. Sterile microtiter wells containing 25 μl of N-104 serially diluted along the ordinate and an equal volume of candidate synthetic peptide serially diluted along the abscissa were incubated aerobically with 100 μl of an overnight culture of PA14 at 10^6^ cfu/ml in 10 mM Na_2_HPO_4_, 1.8 mM KH_2_PO_4_, 2.7 mM KCl, 137 mM NaCl, pH 7.2 at 35°C for 7-8 hours after which 150 μl per well of LB broth was added for an additional 16-18 hours incubation (35°C). Well mixed 2 μl aliquots of each were then dropped onto LB plates for overnight incubation and imaging (Molecular Imager® Gel Doc™ XR System).

The interaction of N-104 with synergistic thrombin peptide GKY20 was explored by drying 1, 5, and 20 μg of GKY20 onto nitrocellulose, blocking with 1% fish gelatin in 10 mM Na_2_HPO_4_, 1.8 mM KH_2_PO_4_, 2.7 mM KCl, 137 mM NaCl, pH 7.2 for 1 hour at room temperature, and then adding 10 ml of 20 μM N-104 or of each N-104 analog, or of negative control C-95, for 2 hours at 35°C with shaking. After washing three times for 10 min, each with 10 ml of 10 mM Na_2_HPO_4_, 1.8 mM KH_2_PO_4_, 2.7 mM KCl, 137 mM NaCl, pH 7.2, nitrocellulose membranes were incubated overnight at 4°C with 1 μg/ml of either rabbit polyclonal ‘ab-C-term’ with specificity for lacritin’s C-terminal 54 amino acids ^103^ appropriate for N-104 or its analogs, or with rabbit anti-lacritin N-terminus antibody ‘ab-N-term’ appropriate for C-95. Membranes washed three times for 10 min each with 10 ml of 10 mM Na_2_HPO_4_, 1.8 mM KH_2_PO_4_, 2.7 mM KCl, 137 mM NaCl, pH 7.2, were then incubated with 0.1 µg/ml of IRDye 680RD labeled donkey antirabbit IgG (926-68073, LI-COR) as per above for Li-COR Odyssey CLx detection.

### Murine corneal infection assay

*In vivo* PA14 infection studies were performed using the murine corneal scratch-injury protocol as previously reported ^66^. Briefly, 18 hour aerobic (37°C) PA14 cultures were prepared from a stock vial inoculated onto tryptic soy agar plates with 5% sheep blood. Bacterial cells were suspended in 1% proteose peptone to an OD_650_ of 1.0, diluted to ~1 to 1.2 × 10^6^ cfu in 5 µl, to prepare the inoculum for ocular infections. Confirmation of the actual inoculum was by diluting this preparation in 1% proteose peptone containing 0.05% Triton X-100 for bacterial enumeration on agar plates. Thirty C57/BL6 mice (7 weeks old; female) anesthetized by intraperitoneal injection of ketamine (100 mg/Kg) and xylazine (10 mg/Kg) - as confirmed by absence of toe pinch response - were each subjected to three 26-gauge needle scratches running from 3 - 5 mm along the length of one cornea followed by placement of 5 μl of PA14 inoculum. Four and eight hours later, 5 μl of a mixture of 1 µM of GKY20 and 20 µM N-104 in 10 mM Na_2_HPO_4_, 1.8 mM KH_2_PO_4_, 2.7 mM KCl, 137 mM NaCl, pH 7.2 was applied to the infected eye in anesthetized mice. After 24 hours, eyes were assigned a 0 to 4 pathology score using the following scheme: 0, eye macroscopically identical to the uninfected contralateral control eye; 1, faint opacity partially covering the pupil; 2, dense opacity covering the pupil; 3, dense opacity covering the entire anterior segment; and 4, perforation of the cornea and/or phthisis bulbi (shrinkage of the globe after inflammatory disease). Mice were euthanized by an overdose of CO_2_ followed by cervical dislocation. Corneas dissected off from enucleated whole eyes were then homogenized in 500 ul of 1% proteose peptone containing 0.05% Triton X-100 for serial dilution and bacteria plating to estimate bacterial CFU infectivity. All procedures were carried out in accordance with the Association for Research in Vision and Ophthalmology resolution on the use of animals in research and were approved by the Brigham and Women’s Hospital Institutional Animal Care and Use Committee, protocol 2018N000003.

### RNAseq of N-104 treated PA14

For RNA extraction and sequencing of *P. aeruginosa*, we used diethylpyrocarbonate (AC170250250, Fisher Scientific) treated Eppendorf tubes (except for those used for synthetic peptide incubations). An aliquot of PA14 overnight culture grown for 3.5 to 4.0 hours in LB broth to OD_600_ = 0.5 (5.0 × 10^8^ cells) was gently pelleted (12,000×g, 30 s), resuspended in 0.5 ml of 1 mM Na_2_HPO_4_, 0.18 mM KH_2_PO_4_, 0.27 mM KCl, 13.7 mM NaCl, pH 7.2 and incubated with 5 or 20 μM N-104, or 20 μM N-80/C-25 for 45 min at 35°C or left untreated. After pelleting (12,000×g, 5 min), cells were resuspended in 0.5 ml of TE buffer (12-090-015, Invitrogen) to which was added 1 ml of RNA-protect Bacteria Reagent (76506, Qiagen) with 5 s of vortexing prior to incubation for 5 min at room temperature, pelleting (5,000×g, 10 min), and lysis initiated in 200 μl of 15 mg/mL lysozyme (PI89833, Fisher Scientific) together with 20 μl of 2.5 mg/mL proteinase K (EO0491, Fisher Scientific), both in TE buffer. Vortexing and agitation was followed by addition of 20 μl of 10% SDS in DEPC-treated water with repeated vortexing and agitation. RNA was then isolated using the RNeasy mini kit (74104, Qiagen) as per manufacturer’s directions. The resulting RNA samples were stored at −80°C. By gel electrophoresis, two sharp 18s and 28s rRNA peaks were apparent with an RNA Integrity Number equivalent (RINe) of 8.4-9.2 and a concentration of approximately 40 - 60 ng/μl. Library construction and sequencing were performed by UVA’s Genome Analysis and Technology Core involving mRNA enrichment via the NEBNext® rRNA Depletion Kit (Bacteria) (E7850L, New England BioLabs), and transcriptome sequencing on a Next Seq 500 Mid Output platform, with 150-bp pair-end reads. The range of raw reads for each sample varied from 8-10 million, thus ensuring an optimal depth of sequencing that was analyzed by UVA’s Bioinformatics Core facility.

## GRAPHICS

Schematic diagrams were generated in PowerPoint (Microsoft).

## QUANTIFICATION AND STATISTICAL ANALYSIS

A minimum of three biological replicates were performed for all experiments, unless explicitly specified otherwise as noted in the Figure Legends. All data are presented as the mean ± SD with single datapoints represented. After an initial normality test, significance was determined by GraphPad Prism (v 10.2) as documented in the Figure Legends. Significant difference was noted as ****p<0.0001, ***p<0.001, **p<0.01, *p<0.05.

## Materials Availability

Addgene plasmid numbers for all unique recombinant DNA generated in this study are documented in the Key Resources Table. These and all other unique reagents generated in this study are also available from the lead contact upon request.

## Data and Code Availability

RNAseq data are available in NCBI GEO DataSets: Accession ID: GSE253123. Tandem mass spectrometry data are available in the MassIVE Repository as: MassIVE MSV00009408 and MassIVE MSV000094086. All data in this paper will be shared by the lead contact upon request. This paper does not report original code. Any additional information required to reanalyze the data reported in this paper is available from the lead contact upon request.

## SUPPLEMENTAL INFORMATION

Supplemental information is attached.

## Supporting information

Key Resource Table

## ACKNOWLEDGEMENTS

We acknowledge UVA’s Nicholas E. Sherman (W.M. Keck Biomedical Mass Spectrometry Laboratory), Pankaj Kumar (Bioinformatics Core) and Michael Solga (Flow Cytometry Core Facility [RRid:SCR_017829], partially supported by P30-CA044579). We also thank Robert M.Q. Shanks (Charles T. Campbell Laboratory of Ophthalmic Microbiology, University of Pittsburgh) for clinical isolates and Robert A. Bloodgood for reviewing the manuscript. This project was supported by NIH 2R01EY026171 (GWL) and R01EY032956 (GWL) and by an unrestricted gift from TearSolutions, Inc (GWL).

## CONTRIBUTIONS

M.S. and G.B. conceived and performed the SPR and QCM-D experiments. M.S.G. and F.N. performed *in vitro* bacterial studies (aided initially by M.R.) including Keio library screens, *C. elegans* feeding assays, iron and ^14^C-polyamine uptake (with help of M.L. and T.E.H.), synergy studies and RNAseq. M.S.G. generated recombinant FeoB (plasmid and key experimental advice from H.V) and PotH, generated mutations and synthetic peptides, carried out binding and imaging flow cytometry studies and with A.A. and G.A wide-field fluorescence imaging. G.M.O. and M.M.P. completed Bacterial ChloroAlkane Penetration Assays. M.G. provided PA14 transposon mutants and offered key experimental advice. G.B.P. and T.S.Z. performed *in vivo* bacterial studies. M.S.G. and G.W.L. conceived the study and with F.N. wrote the article with help from all authors.

## KEY RESOURCES TABLE

[Dum]

## SUPPLEMENTAL FILE INFORMATION

### SUPPLEMENTARY INFORMATION

Formulaic analysis of ChloroAlkane Penetration Assay data, the consequence of which is depicted in **Figures 3I and 3J**. Raw data is presented in **Figures S3D and S3E**.

### SUPPLEMENTAL TABLES

**Supplemental Table S1. Bacterial orthologs of identified N-104 mediators, as related to Figure 2** Orthologs of Feob, PotH, RluE, YbaE, YbdM, Yeel, YhfZ, YhjQ, YobD, WcaD, TauD and YaiW among bacterial phyla, classes, orders, genera and species, as identifed by BLAST.

**Supplemental Table S2. Putative ligands of identified N-104 mediators, as related to Figure 2** Putative ligands of Feob, PotH, RluE, YbaE, YbdM, Yeel, YhfZ, YhjQ, YobD, WcaD, TauD and YaiW, as suggested by BioGRID and STRING.

**Supplemental Table S3. Unbiased MS/MS screen of putative N-104 binding proteins in PA14 lysate, as related to Figures 3 and S3**

MS/MS identified PA14 proteins KCl eluted from N-104 versus negative control C-95 columns.

**Supplemental Table S4. *feoB* and *potH* site-directed mutagenesis and sequencing primers as related to Figures 4 and S4**

*feoB* and *potH* periplasmic outer loops (OL) were subjected to mutagenesis.

**Supplemental Table S5. Unbiased MS/MS screen of putative N-104 binding proteins in human basal tears, as related to Figures 6 and S6**

MS/MS identified tear proteins KCl eluted from N-104 versus negative control C-95 columns.

**Supplemental Table S6. Altered transcription in: (i) 5 μM N-104 treated *vs.* untreated, (ii) 20 μM N-104 treated *vs.* untreated and (iii) 20 μM N-104 treated *vs.* 20 μM N-80/C-25 treated PA14, as related to Figure S8**

Changes common to (i) and (iii)* and (i) - (iii)** are noted.

### SUPPLEMENTAL FIGURE LEGENDS

**Figure S1.**
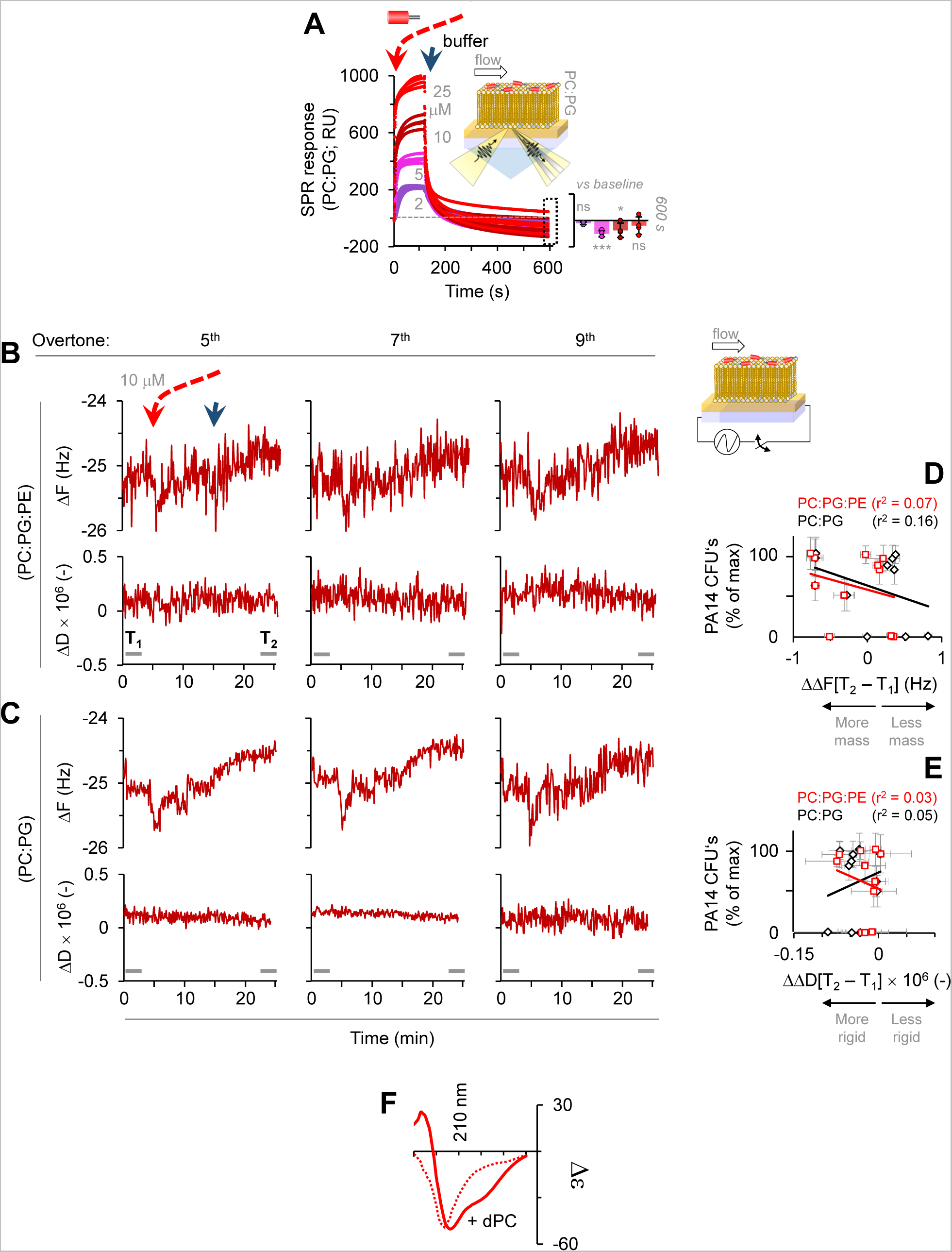
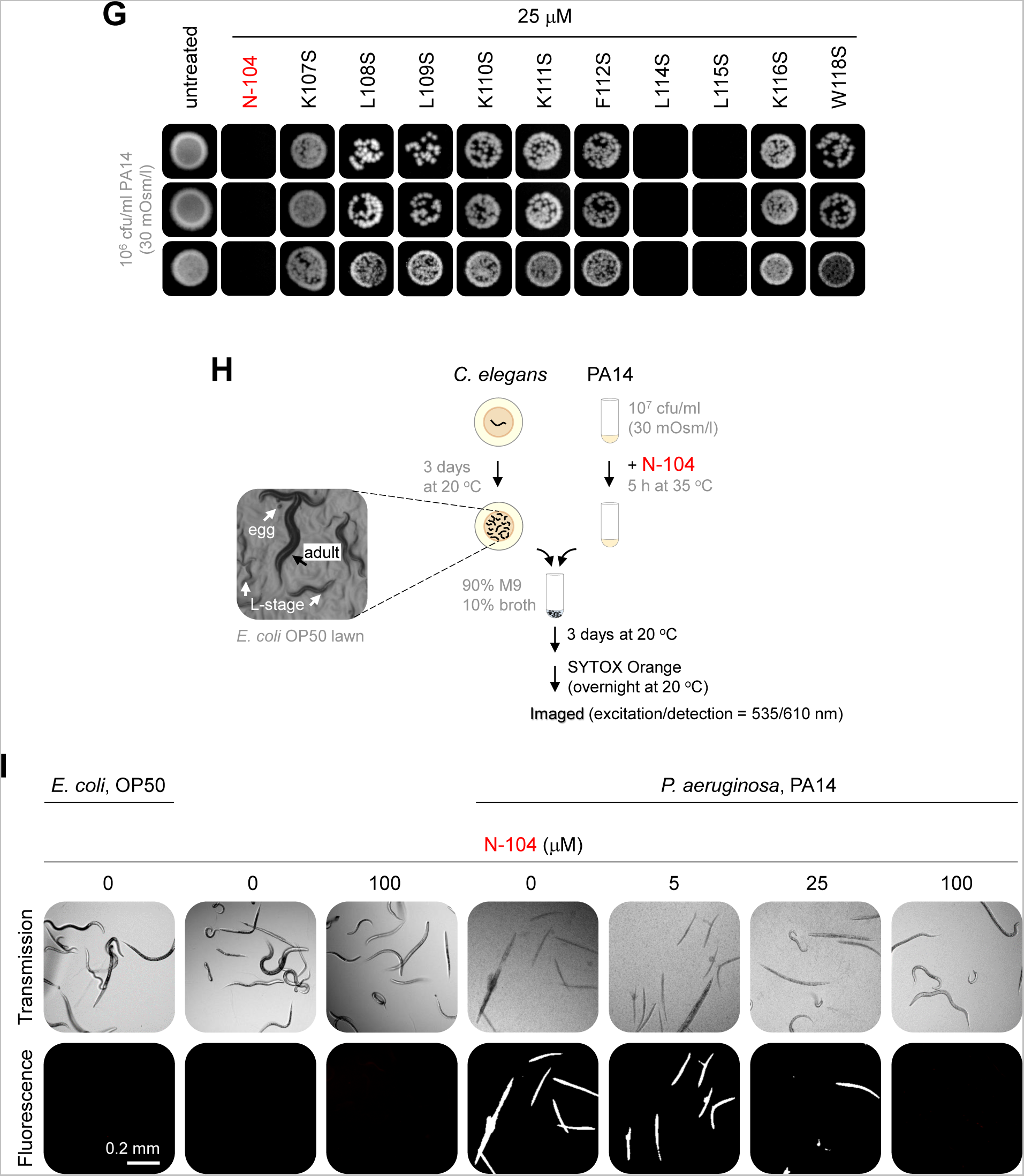
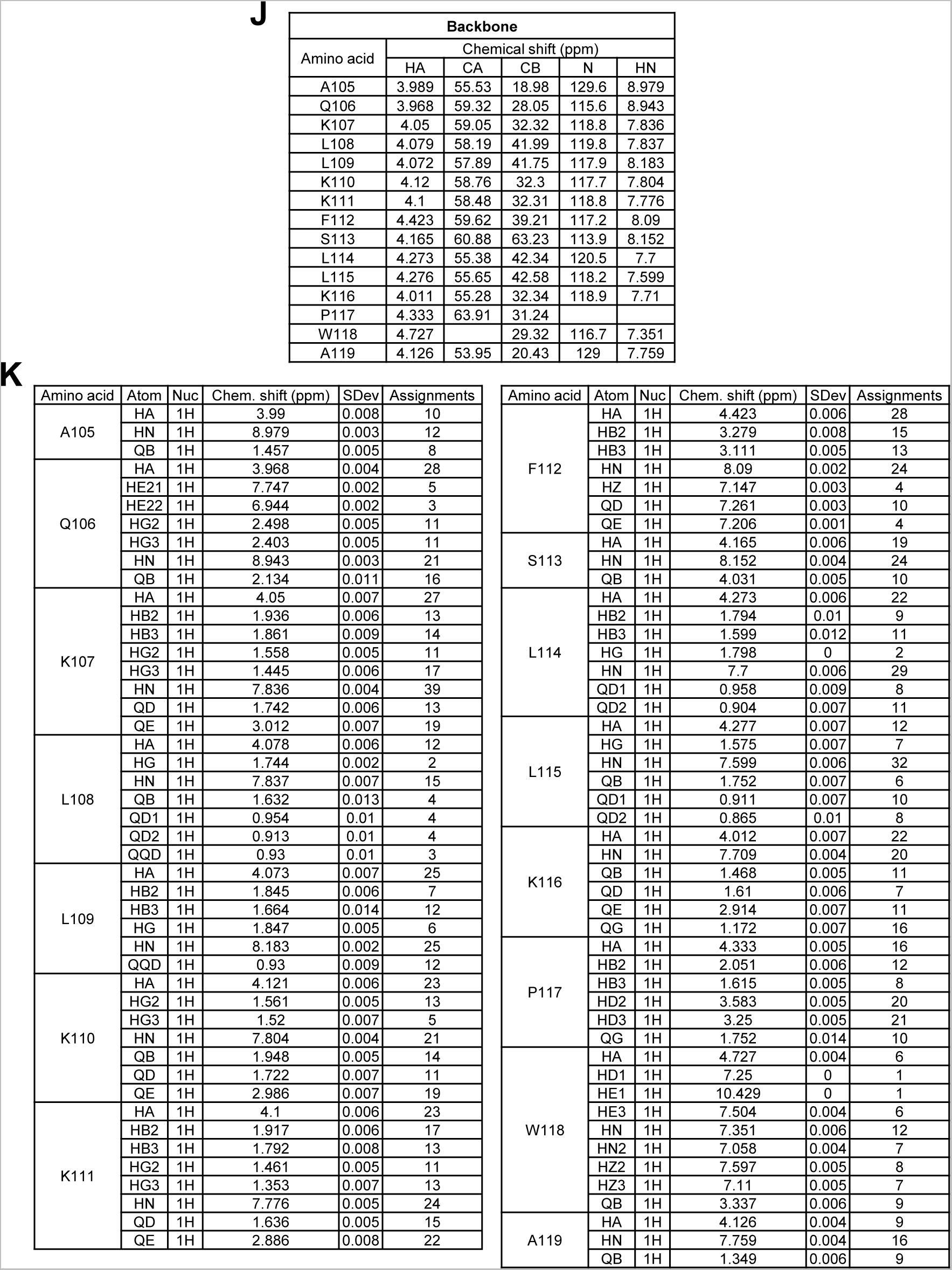
Lysis-free ‘N-104’ killing of virulent and multi-drug resistant *P. aeruginosa* strain ‘PA14’ together with biological replicates related to Figure 1. (A) N-104 interaction with Gram-positive model membrane as monitored by surface plasmon resonance (SPR). Insert: SPR response at 600 s. Two-way ANOVA with Tukey’s multiple comparisons test versus baseline for each N-104 concentration (n = 4 experiments). (B) N-104 interaction with Gram-negative model membrane as monitored by quartz crystal microbalance with dissipation monitoring (QCM-D). Real time monitoring of ΔF and ΔD of the sinusoidal waves at overtones 5, 7 and 9. (n = 3 experiments). (C) N-104 interaction with Gram-positve model membrane as monitored by QCM-D. (n = 3 experiments). (D) Lack of correlation between the interaction (ΔΔF) of N-104 and N-104 analogs with both Gram-negative and -positive model membranes and reduction of PA14 colony forming units. (n = 3 experiments). (E) Lack of correlation between the interaction (ΔΔD) of N-104 and N-104 analogs with both Gram-negative and -positive model membranes and reduction of PA14 colony forming units. (n = 3 experiments). (F) Circular dichroism of N-104 in the presence or absence of 10 mM dodecylphosphocholine. (G) Figure 1D biological replicates. (H) Schematic of *C. elegans* liquid infection assay in which dead worms are distinguished by uptake of SYTOX Orange. (I) Only *C. elegans* that have consumed N-104 killed PA14 survive (lack of SYTOX Orange fluorescence). N-104 alone has no apparent effect on *C. elegans*. (J,K) N-104 NMR assignments. Graphs represent the mean ± SD, ***p<0.001, *p<0.05, ns, nonsignificant.

**Figure S2.**
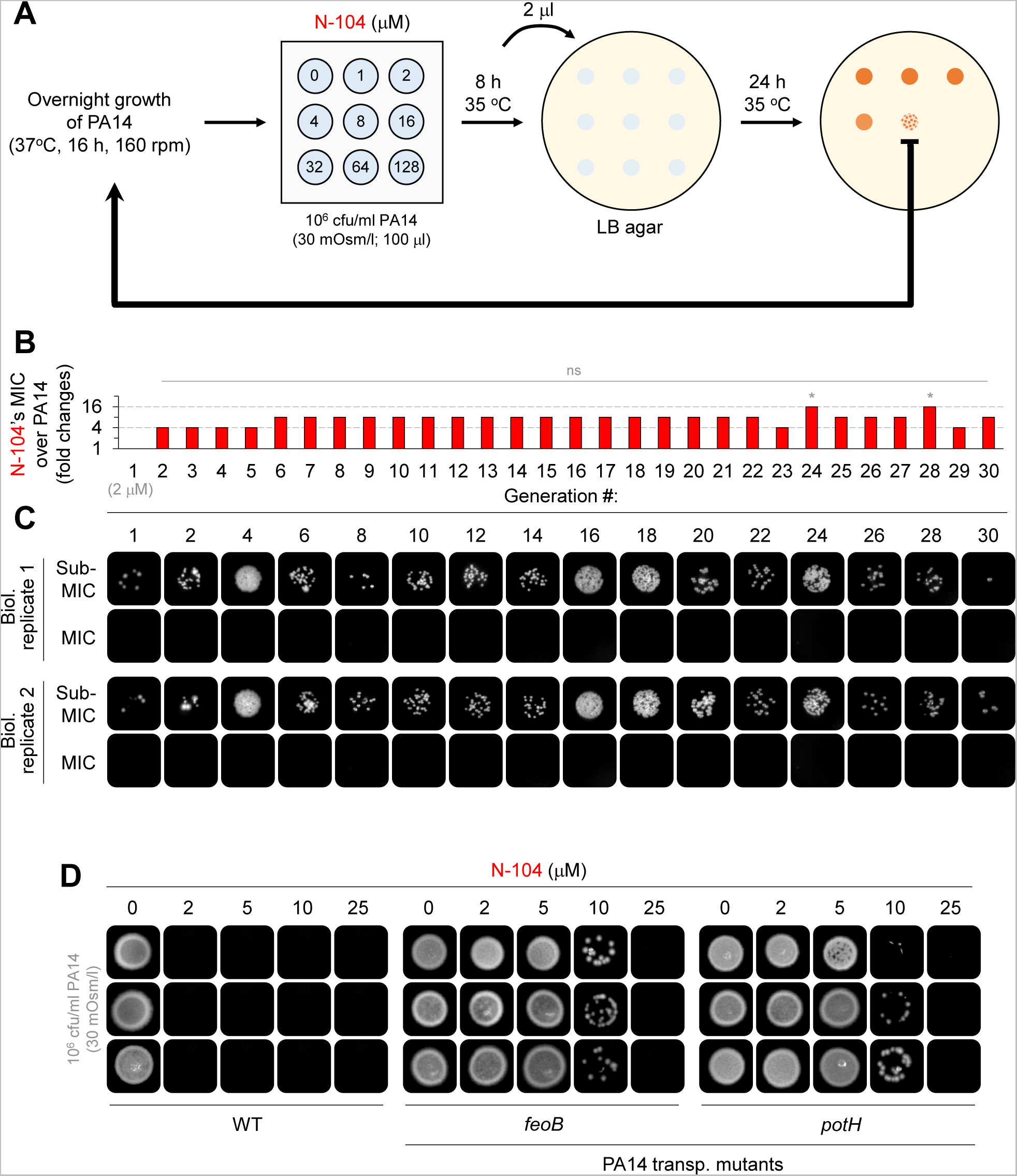
Thirty generations of PA14 fail to develop stable N-104 resistance together with biological replicates related to Figure 2. (A) Schematic of the N-104 resistance assay. (B) Despite deviations at generations 24 and 28, no PA14 resistance to N-104 developed over 30 generations. Friedman ANOVA with Dunn’s multiple comparisons test (n = 2 experiments). (C) N-104 killing assays over 30 generations of PA14. (D) Biological replicates of N-104 killing assays of PA14 wild type, or PA14 *feoB* or *potH* mutants. Graphs represent the mean ± SD, *p<0.05, ns, nonsignificant.

**Figure S3.**
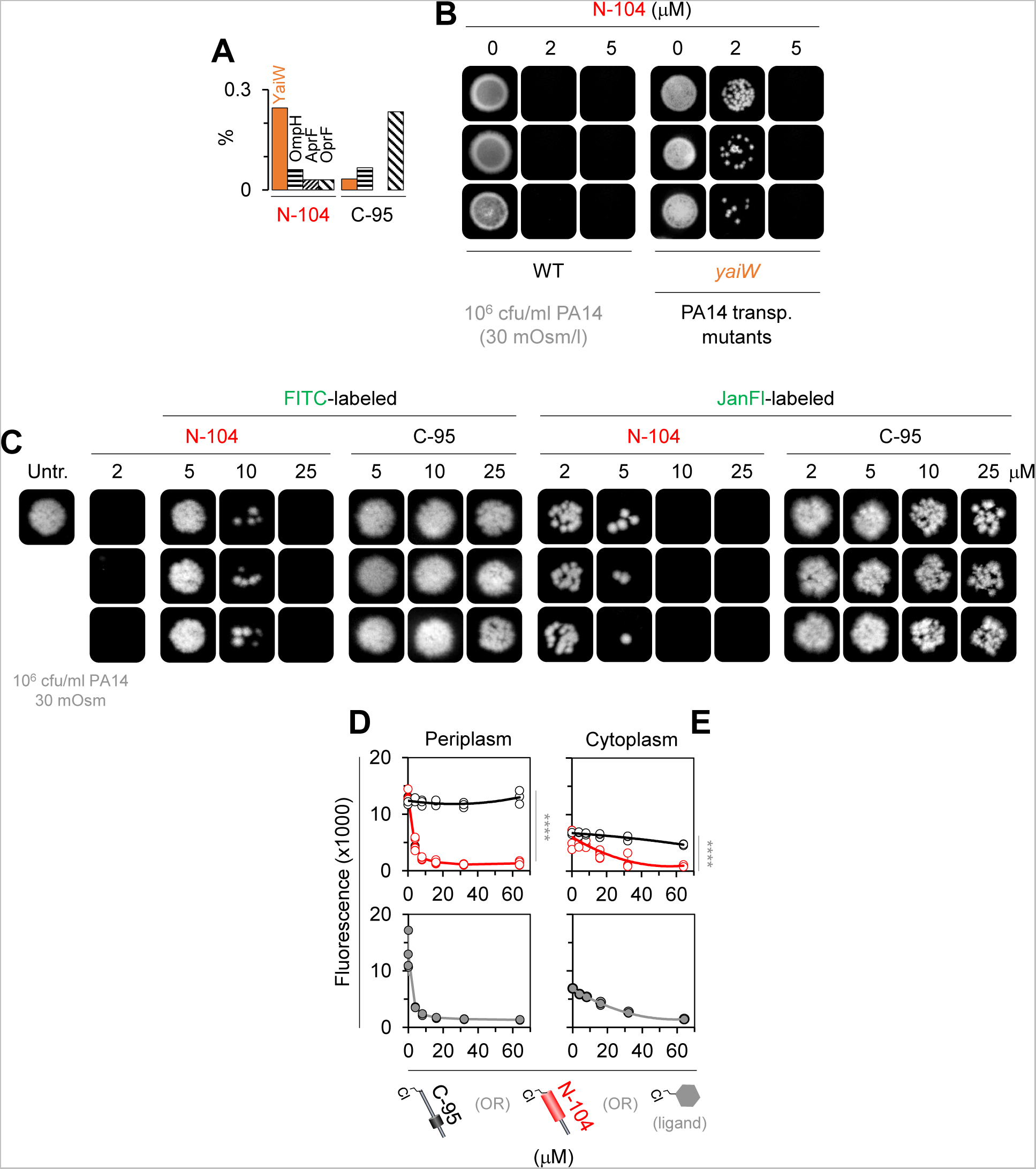
N-104 activity is partially affected by FITC and Janelia Fluor 549 tagging together with biological replicates related to Figure 3 and Table S3. (A) N-104 column enrichment of the outer membrane lipoprotein YaiW (orange bar) relative to other N-104- or C-95-enriched outer membrane proteins including OmpH, AprF and OprF. (B) Biological replicate N-104 killing assays of PA14 wild type or *yaiW* mutant cells. (C) Biological replicates of FITC- and Janelia Fluor 549-tagged killing assays of PA14. (D) Top, successful competition by chloroalkane tagged N-104 against a chloroalkane-modified fluorescent dye for a HaloTag periplasmic marker (as per the positive control (bottom)). The same is not true for negative control chloroalkane tagged C-95. Two-way ANOVA with Šidák’s multiple comparisons test (n = 3 experiments). (E) Top, slight competition of chloroalkane tagged N-104 (as per the positive control (bottom)) but not similarly tagged C-95 against chloroalkane modified fluorescent dye for HaloTag cytoplasmic marker (D, E; same original data as Figures 3I **and 3J** but here showing raw fluorescence). Two-way ANOVA with Šidák’s multiple comparisons test (n = 3 experiments). Graphs represent the mean ± SD, ****p<0.0001.

**Figure S4.**
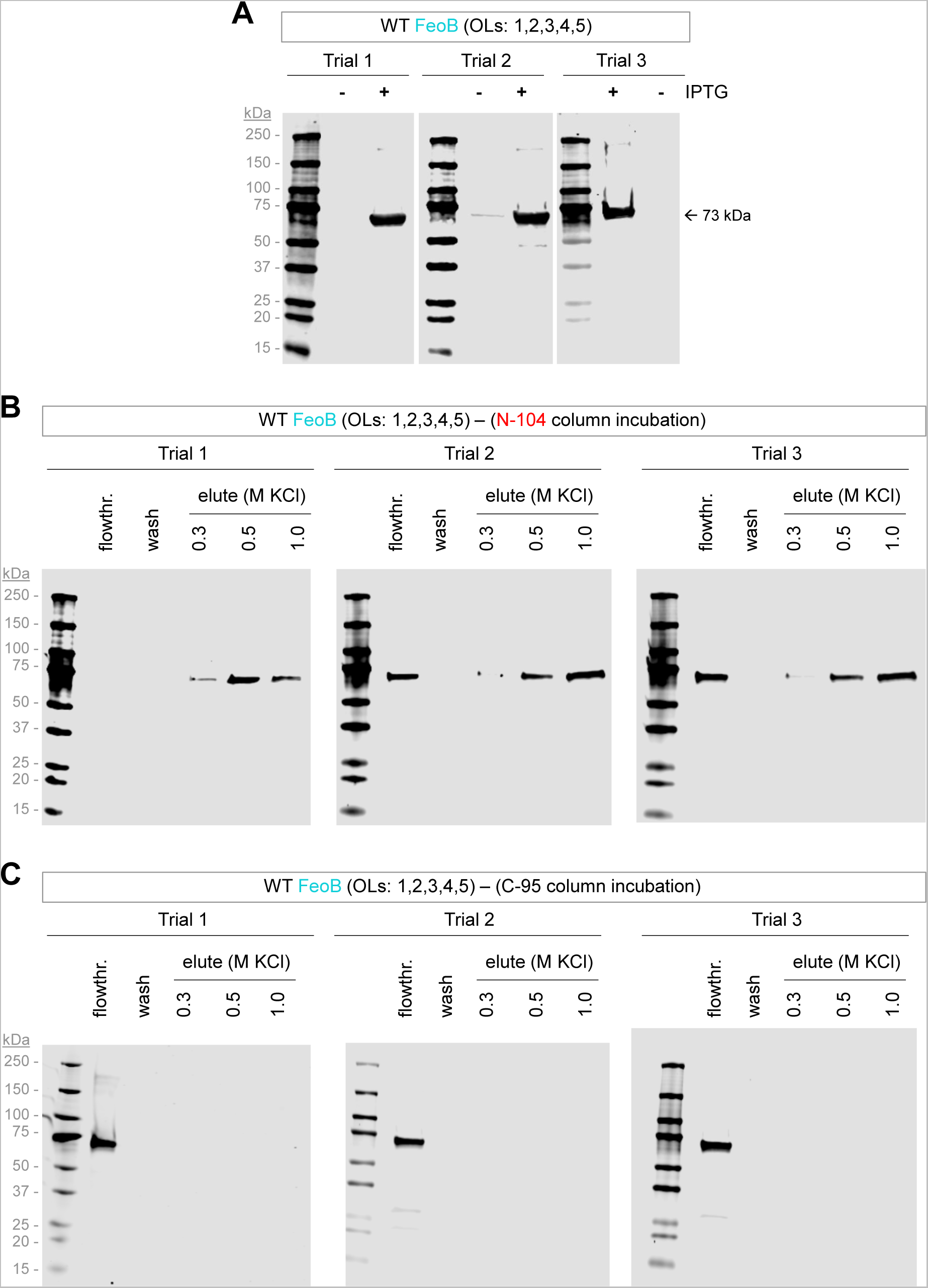
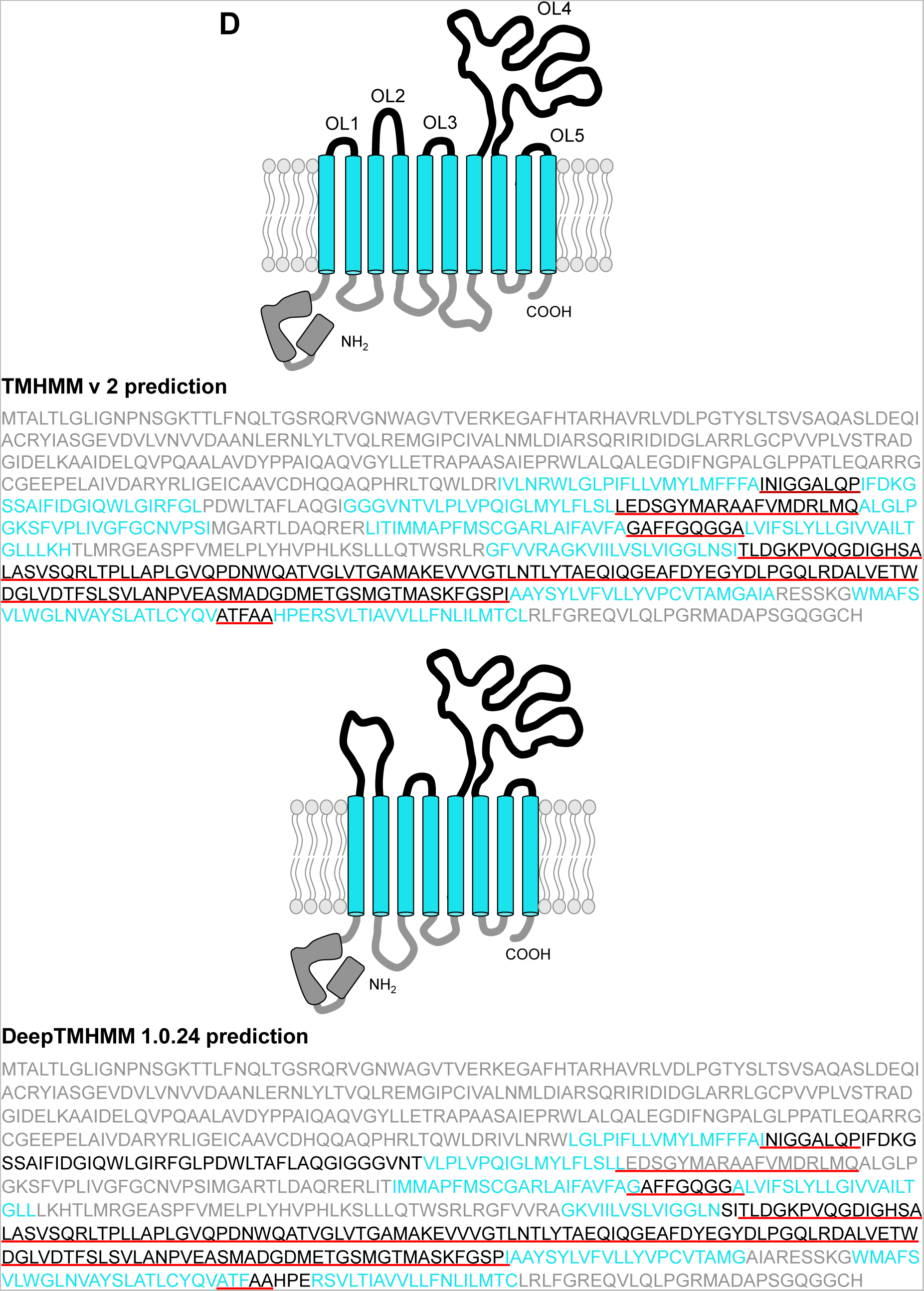
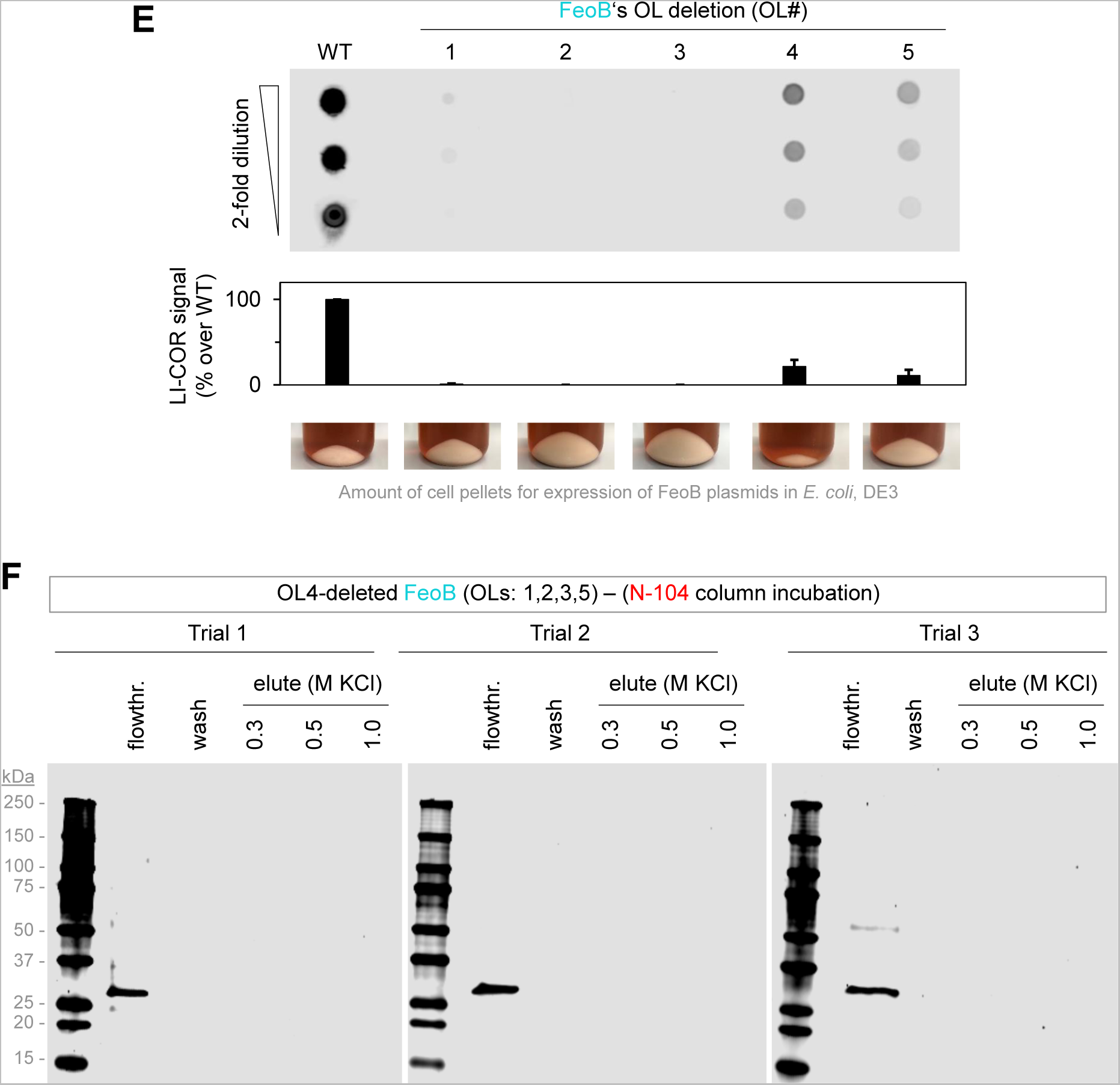
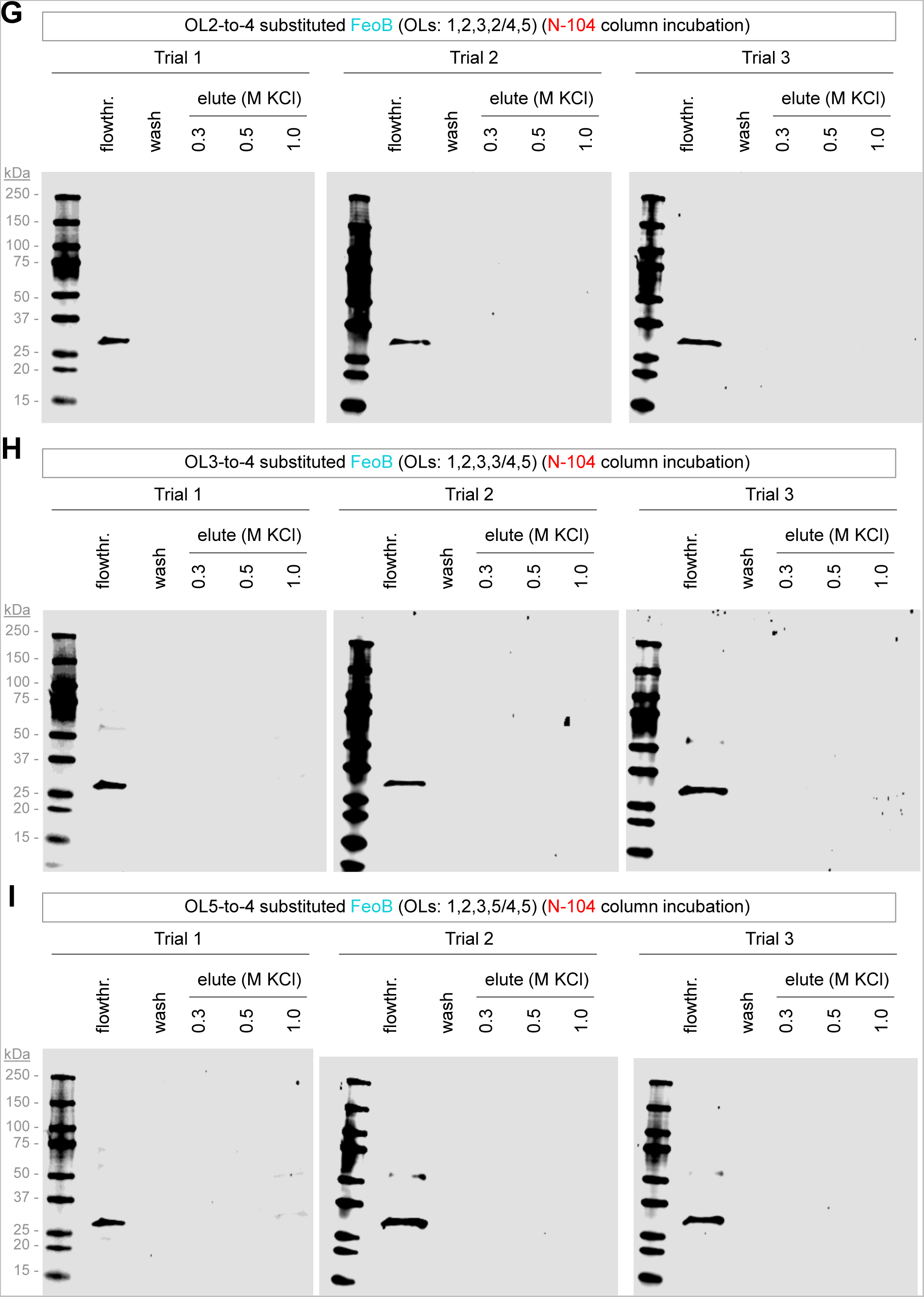
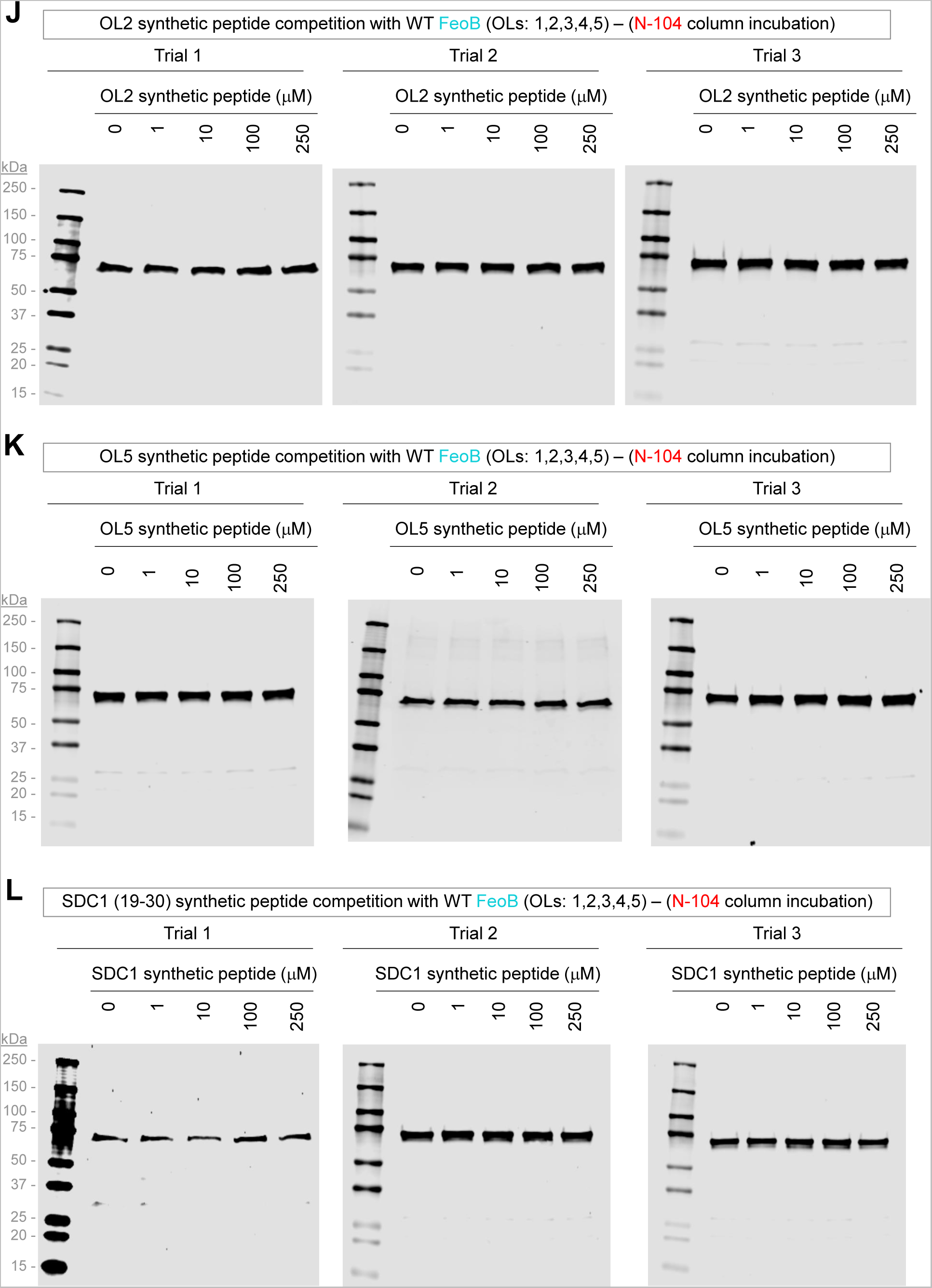
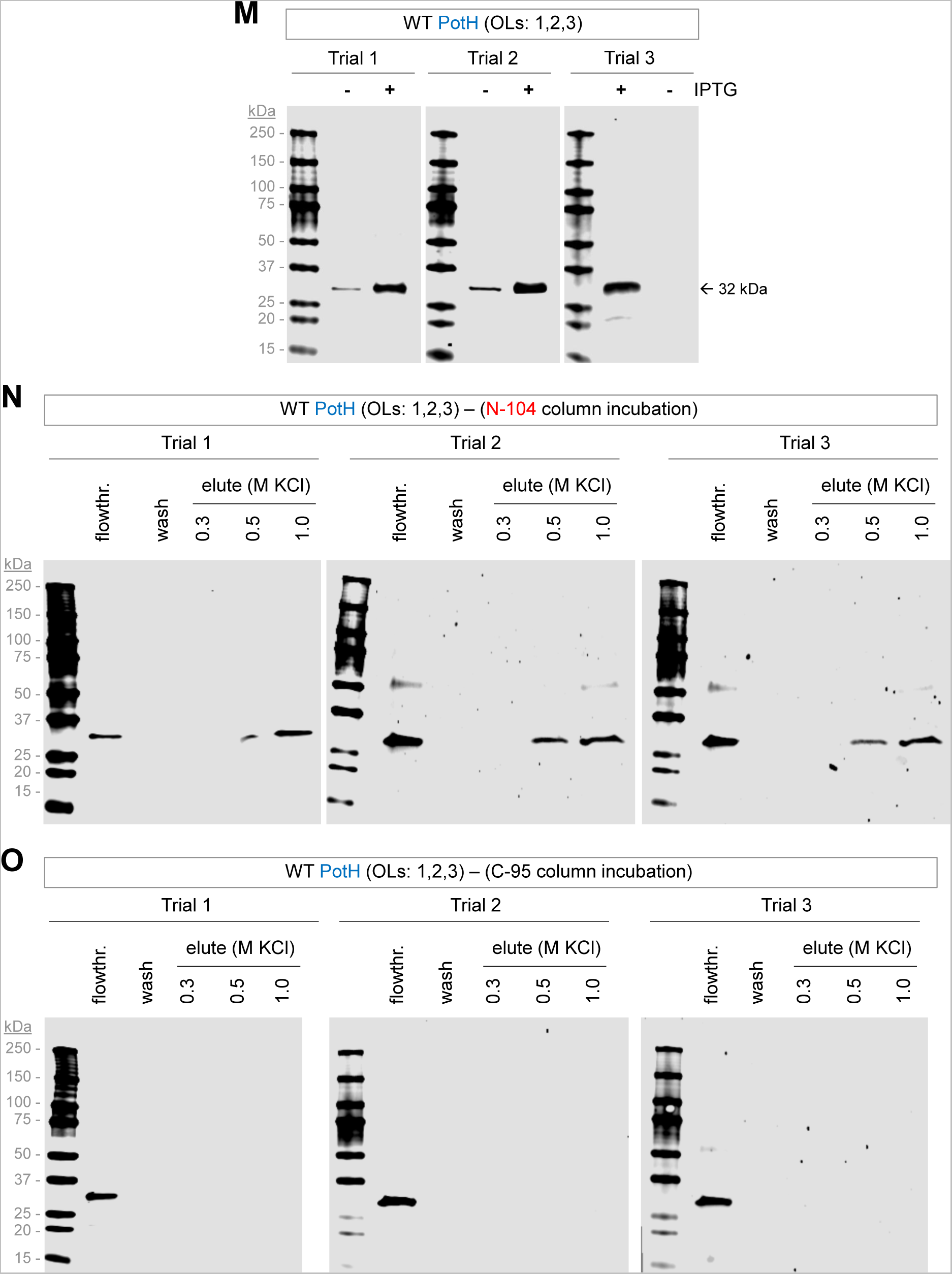
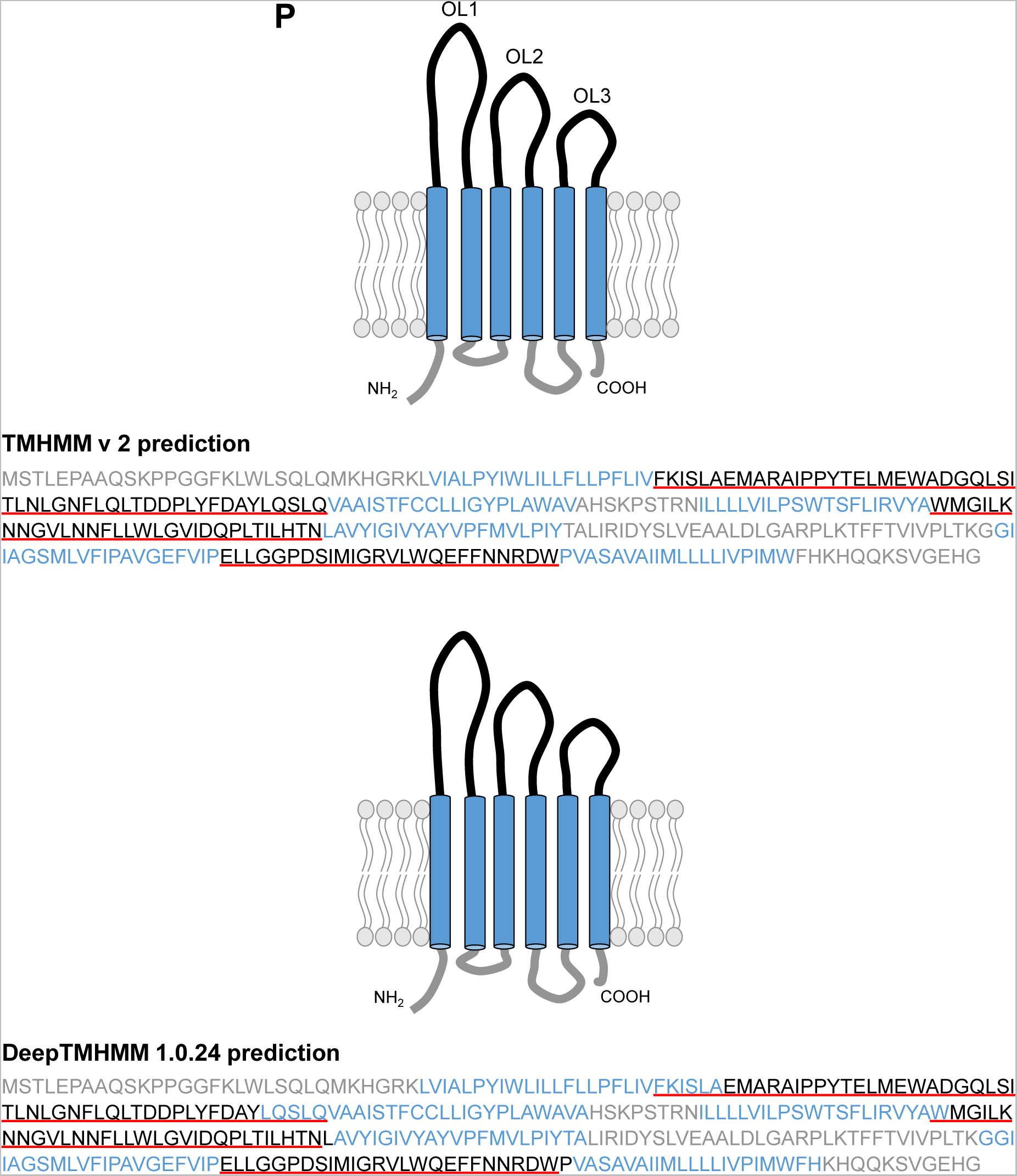
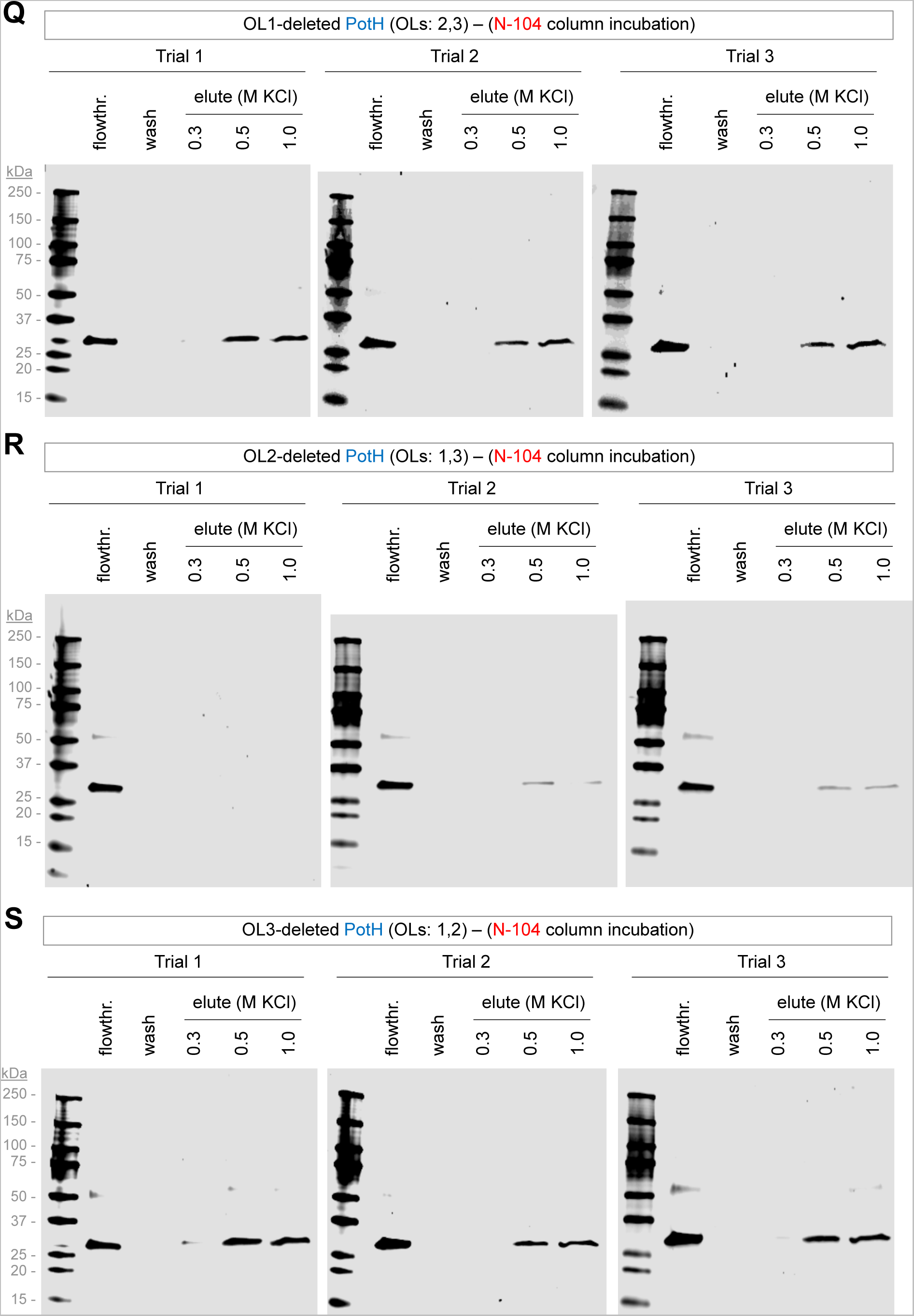
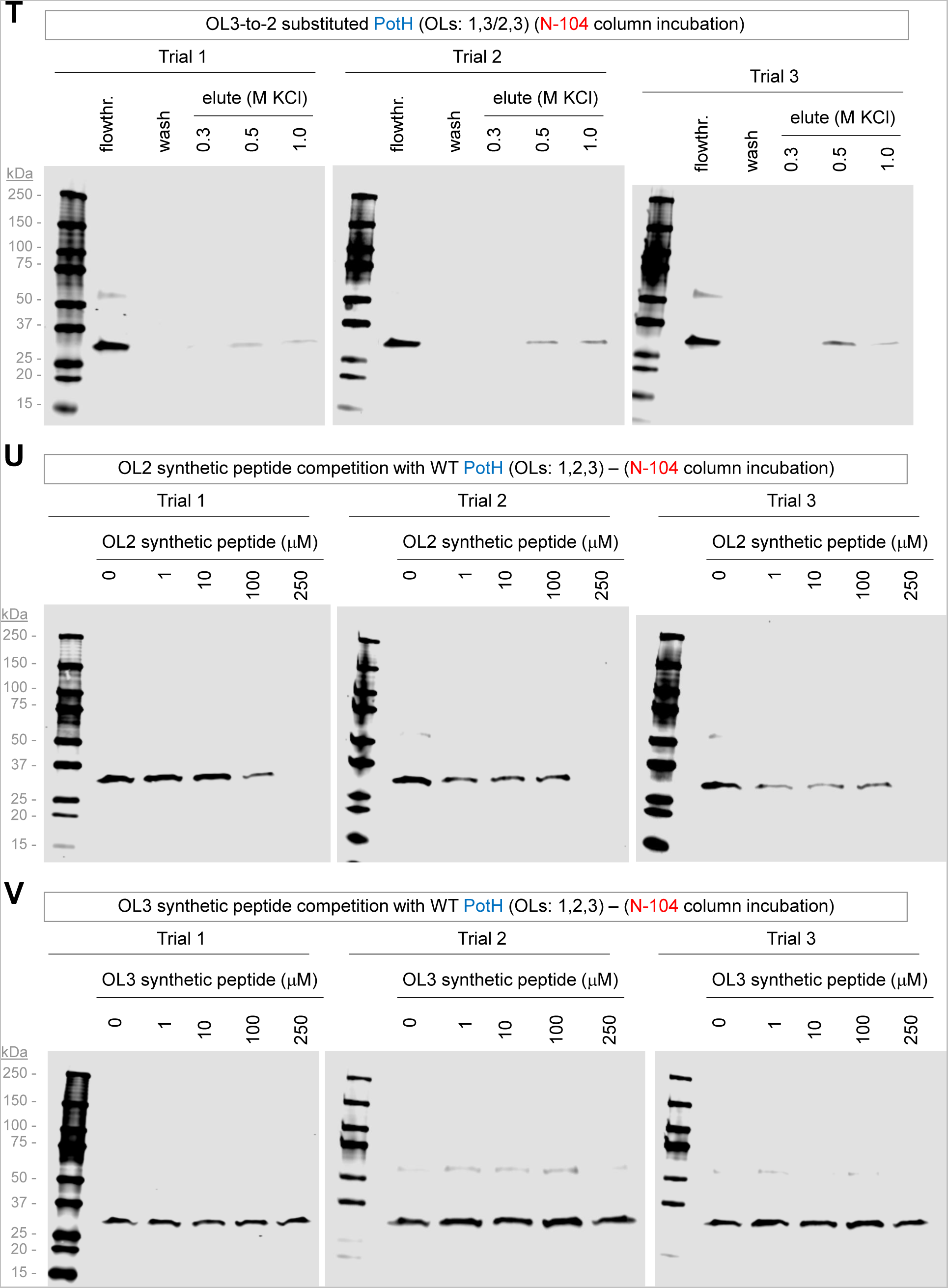
Anti-His western blot biological replicates of FeoB and PotH targeting N-104, as related to Figure 4 and Table S4. (A) IPTG-induced expression of His tagged FeoB. (B) His tagged FeoB binds column immobilized N-104, as per absence in the final wash fraction and detection in KCl elution fractions. (C) His tagged FeoB does not bind column immobilized C-95 (negative control), as suggested by lack of KCl elution. (D) TMHMM2 and Deep TMHMM predicted FeoB periplasmic loops (after ^51^). (E) Comparative recombinant production of His tagged FeoB, or His tagged FeoB lacking periplasmic outer loop (’OL’) 1, 2, 3, 4 or 5 (TMHMM2 prediction). Not detected is His tagged FeoB lacking periplasmic outer loop 2 or 3. Without outer loop 1 expression is very low. Without outer loop 4 or 5, expression is higher but nonetheless much less than wild type. (F) His tagged FeoB lacking periplasmic outer loop 4 fails to bind column immobilized N-104, as suggested by lack of KCl elution. (G) His tagged FeoB in which outer loop 4 has been replaced with outer loop 2 fails to bind column immobilized N-104, as suggested by lack of KCl elution. (H) His tagged FeoH in which outer loop 4 has been replaced with outer loop 3 fails to bind column immobilized N-104, as suggested by lack of KCl elution. (I) His tagged FeoB in which outer loop 4 has been replaced with outer loop 5 fails to bind column immobilized N-104, as suggested by lack of KCl elution. (J) FeoB outer loop 2 synthetic peptide fails to compete His tagged FeoB off N-104 columns. (K) FeoB outer loop 5 synthetic peptide fails to compete His tagged FeoB off N-104 columns. (L) SDC1 (19-30) synthetic peptide ^52^ (with similarity to FeoB outer loop 1) fails to compete His tagged FeoB off N-104 columns. (M) IPTG-induced expression of His tagged PotH. (N) His tagged PotH binds column immobilized N-104, as per absence in the final wash fraction and detection in KCl elution fractions. (O) His tagged PotH does not bind column immobilized C-95 (negative control), as suggested by lack of KCl elution. (P) TMHMM2 and Deep TMHMM predicted PotH periplasmic loops. (Q) His tagged PotH lacking periplasmic outer loop 1 binds column immobilized N-104, as per absence in the final wash fraction and detection in the KCl elution fractions. (R) His tagged PotH lacking periplasmic outer loop 2 displays no or slight binding to column immobilized N-104, as per KCl elution. (S) His tagged PotH lacking periplasmic outer loop 3 binds column immobilized N-104, as per absence in the final wash fraction and detection in KCl elution fractions. (T) His tagged PotH in which outer loop 2 has been replaced with outer loop 3 poorly binds column immobilized N-104, as suggested by minimal KCl elution. (U) PotH outer loop 2 synthetic peptide partially competes His tagged PotH off of N-104 columns. (V) PotH outer loop 3 synthetic peptide fails to compete His tagged PotH off of N-104 columns.

**Figure S5.**
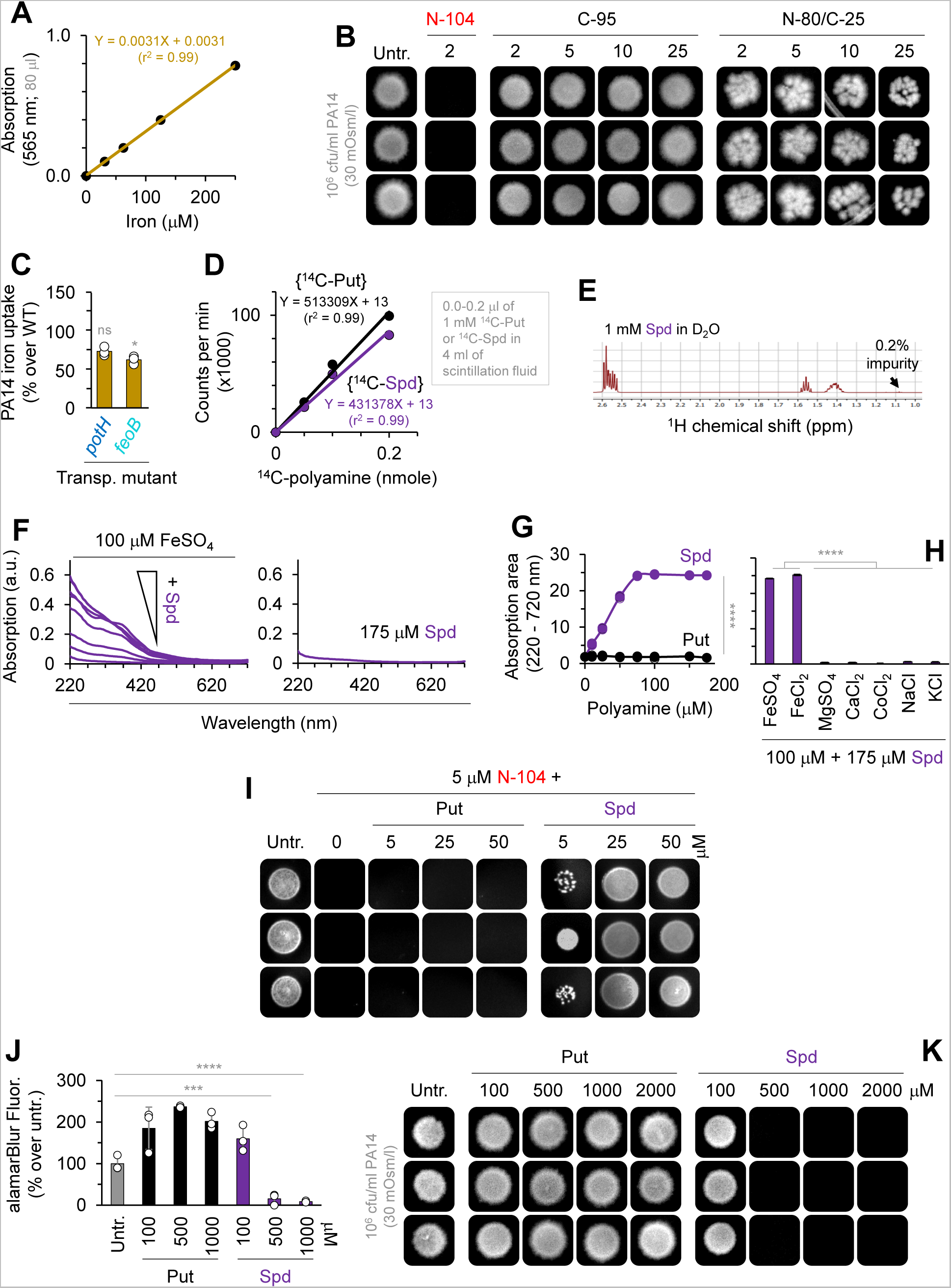
Characterization of spermidine and putrescine uptake efficiency and the affinity of the former for ferrous iron in PA14 cells, together with biological replicates related to Figure 5. (A) Absorbance at 595 nm of increasing concentrations of Fe^2+^. (B) N-104 versus negative control C-95 or N-80/C-25 in PA14 killing assays. (C) Comparative Fe^2+^ uptake in iron-depleted PA14 cells lacking *feoB* or *potH*, comparison versus wild type. Kruskal-Wallis ANOVA with Dunn’s multiple comparisons test (n = 3 experiments). (D) Scintillation counter detection of increasing concentrations of the polyamines ^14^C-spermidine (’^14^C-Spd’) or ^14^C-putrescine (^14^C-Put). (E) 99.8% purity of of ^14^C-spermidine, as per ^1^H-NMR spectra. (F) Left, increasing absorbance of FeSO_4_ with increasing concentrations (0, 10, 25, 50, 75, 100, 150, 175 µM) of spermidine. Right, lack of absorbance of spermidine alone. (G) Binding of spermidine but not putrescine to 100 µM FeSO_4_. Two-way ANOVA (n = 3 experiments). (H) Spermidine bind FeSO_4_ and FeCl_2_, but not MgSO_4_, CaCl_2_, CoCl_2_, NaCl nor KCl. One-way ANOVA with Tukey’s multiple comparisons test (n = 3 experiments). (I) Spermidine, but not putrescine, inhibits N-104 bactericidal activity in a concentration-dependent manner. (J) At concentrations greater than 100 µM, spermidine (but not putrescine) is toxic for PA14. Two-way ANOVA with uncorrected Fisher’s least significant difference test (n = 3 experiments). (K) High concentration spermidine killing assay in PA14. Graphs represent the mean ± SD, ****p<0.0001, ***p<0.001, *p<0.05, ns, nonsignificant.

**Figure S6.**
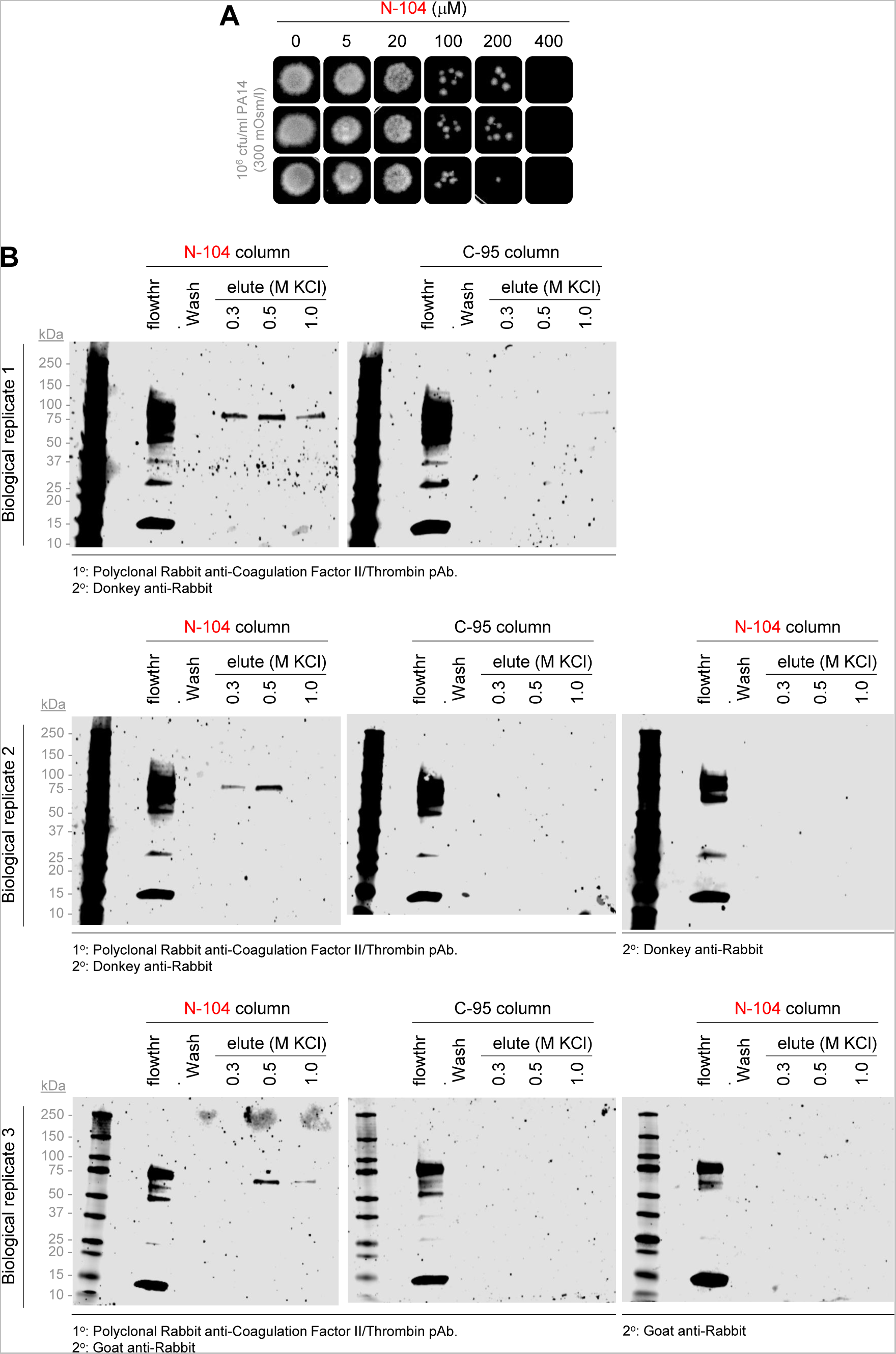
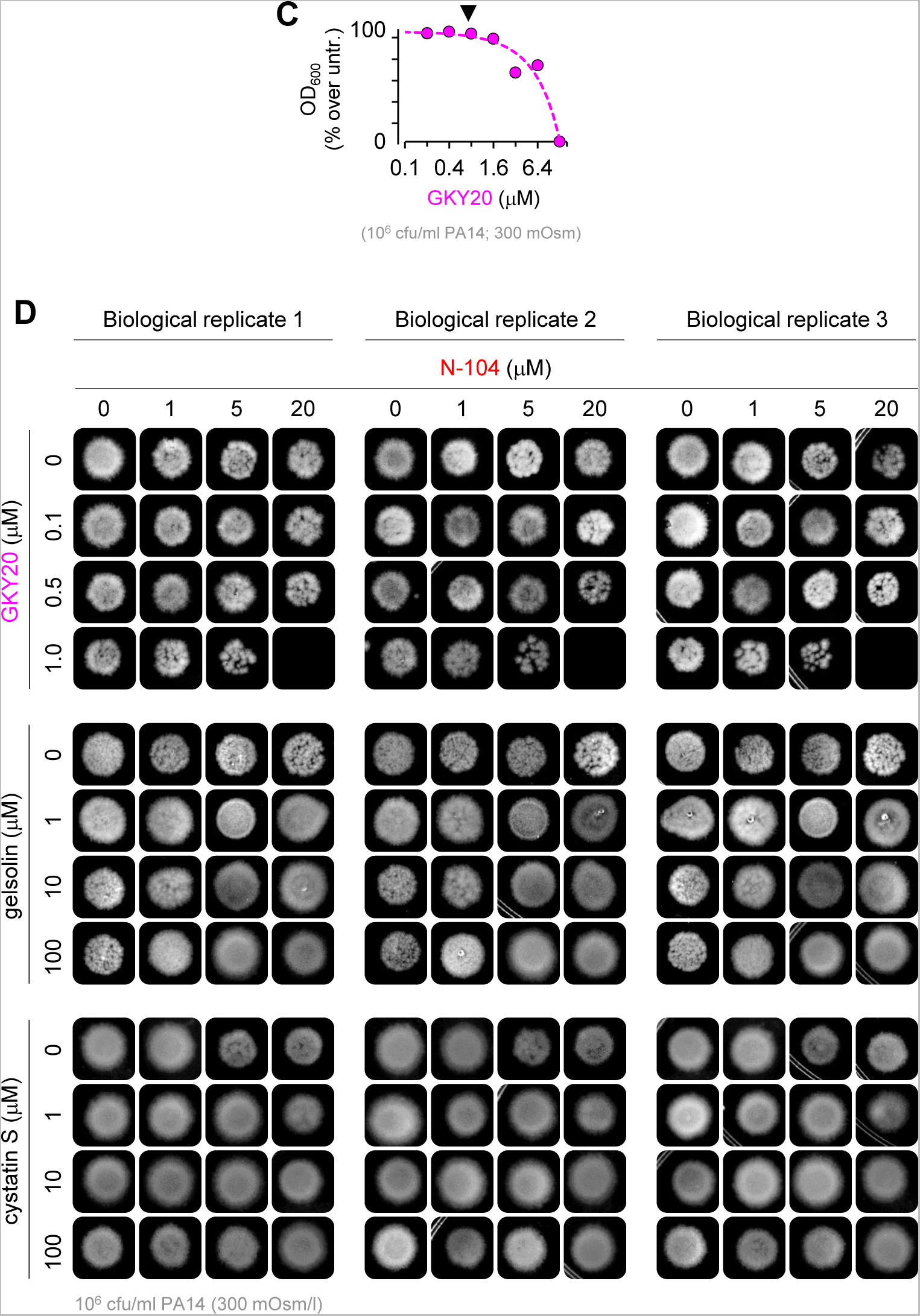
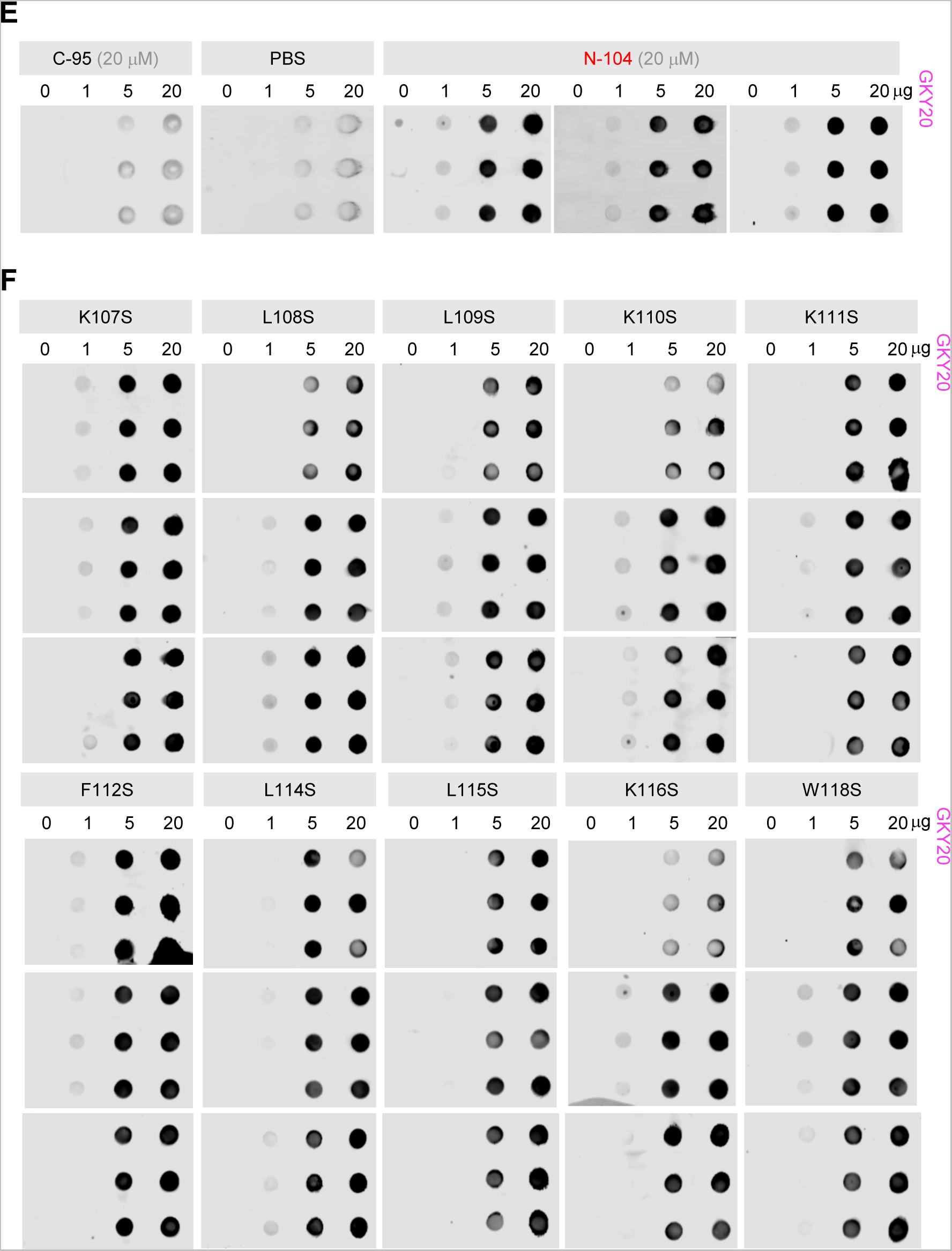
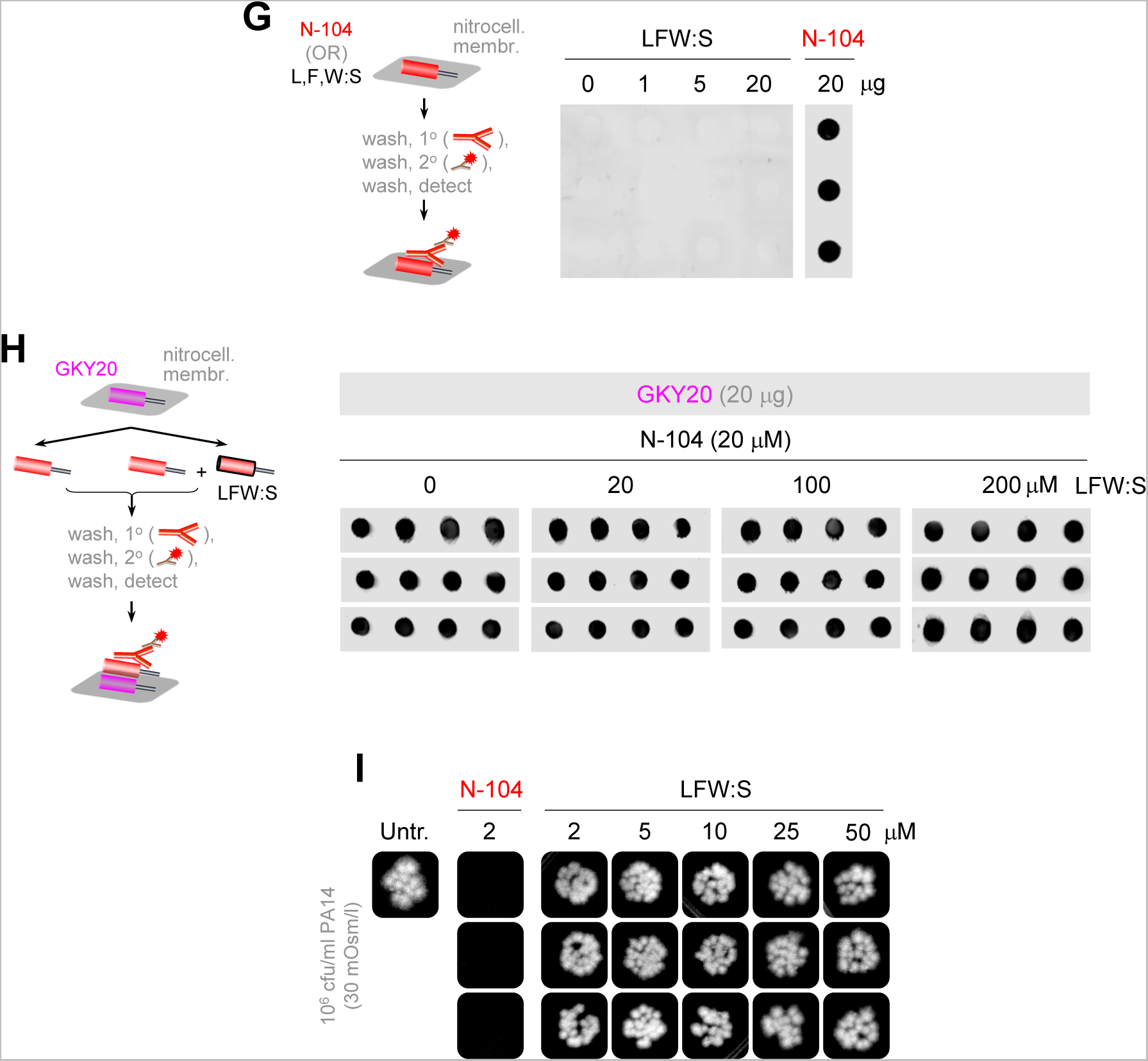
N-104 efficacy at 300 mOsm/l and synergy with thrombin GKY20 synthetic peptide, together with biological replicates related to Figure 6 and Table S5. (A) N-104 killing assay of PA14 at 300 mOsm/l. (B) N-104 column capture of thrombin from pooled human basal tears, as per absence in the final wash fraction and detection in the KCl elution fractions. None bound to the negative control C-95 column. (C) GKY20 solution killing assay of PA14. Optimization of GKY20 concentration for checkerboard assay with N-104. (D) Replicates of N-104 checkerboard assays with GKY20 or each with peptides of gelsolin and cystatin S. (E) N-104, but not C-95, binding of immobilized thrombin peptide GKY20 as respectively detected with rabbit polyclonal ab-C-term or ab-N-term anti-lacritin antibodies. (F) N-104 and N-104 analog ligation of immobilized thrombin peptide GKY20 as detected with rabbit polyclonal ab-C-term anti-lacritin antibodies. (G) Ab-C-term antibody detects N-104 but not N-104 ‘L108S/L109S/F112S/L114S/L115S/W118S’ (’LFW:S’). (H) Since ab-C-term antibody cannot detect N-104 LFW:S, the capacity of increasing amounts of LFW:S to inhibit N-104 ligation of GKY20 was assessed. (I) N-104 versus LFW:S killing assay of PA14.

**Figure S7.**
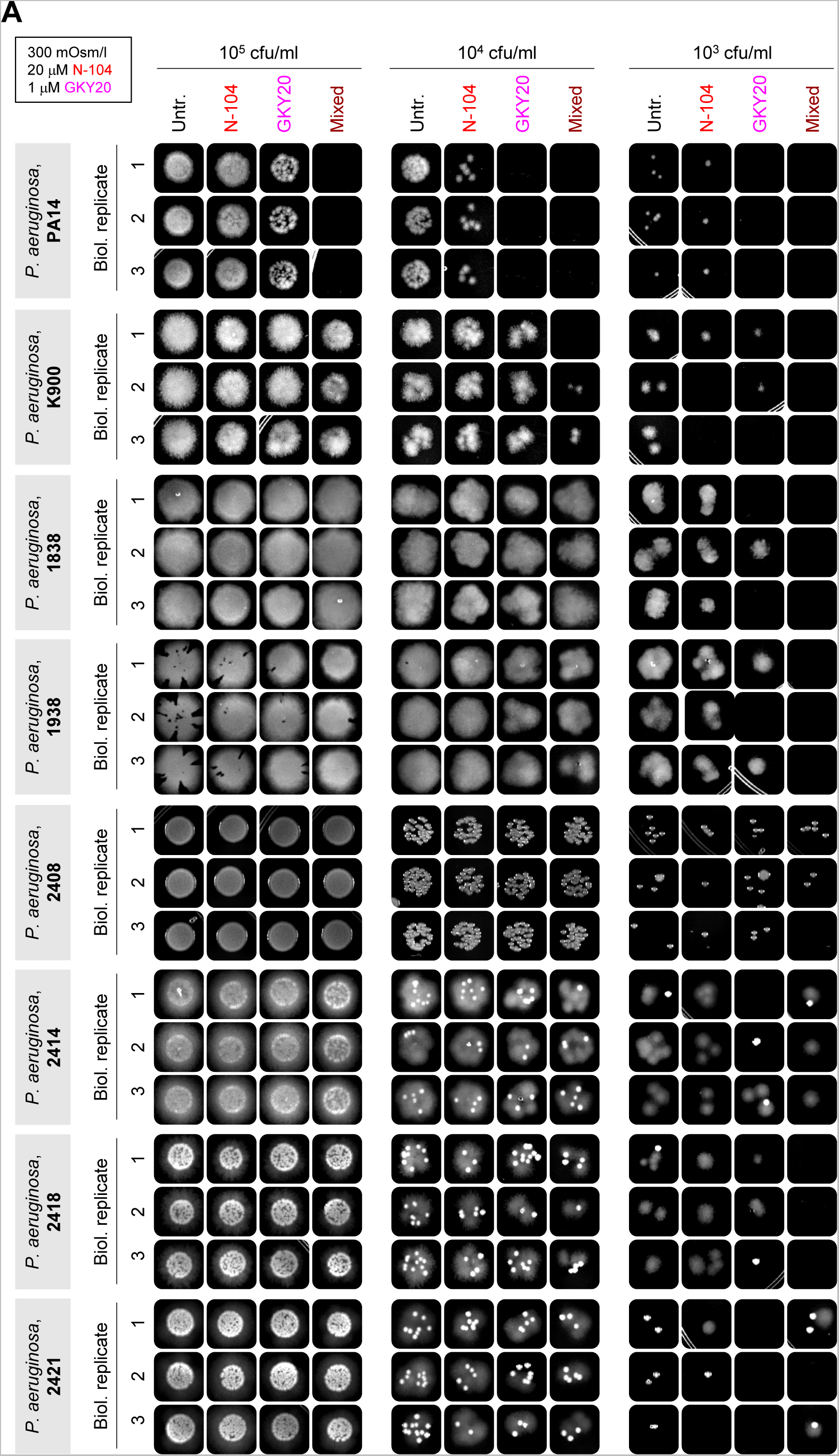
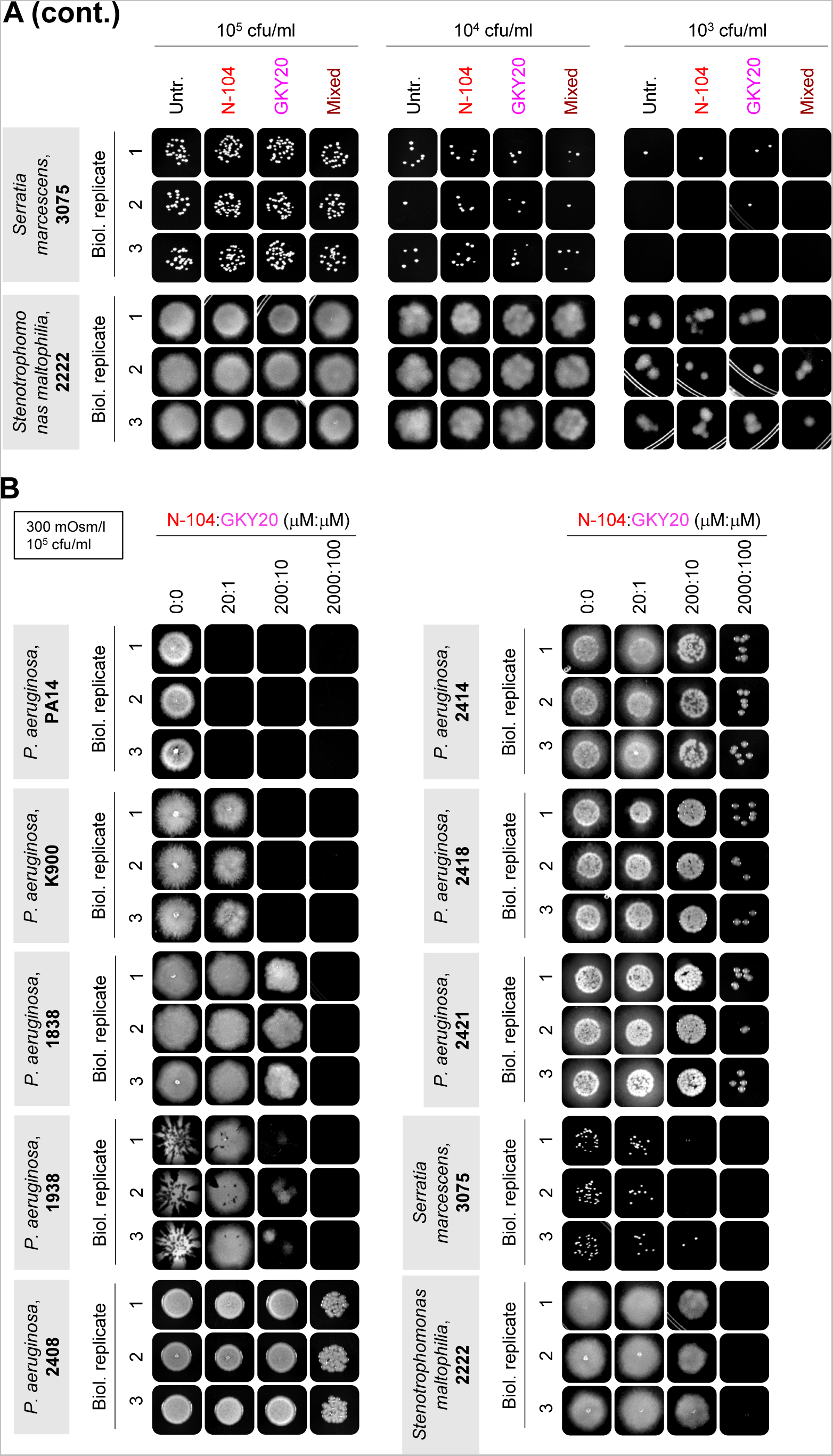
N-104 and N-104 plus GKY20 killing assays of clinical isolates versus PA14 as biological replicates related to Figure 7. (A) Comparative killing assays of PA14 and clinical isolates at increasing densities. (B) Comparative killing assays of PA14 and clinical isolates at increasing N-104:GKY20 molar concentrations.

**Figure S8.**
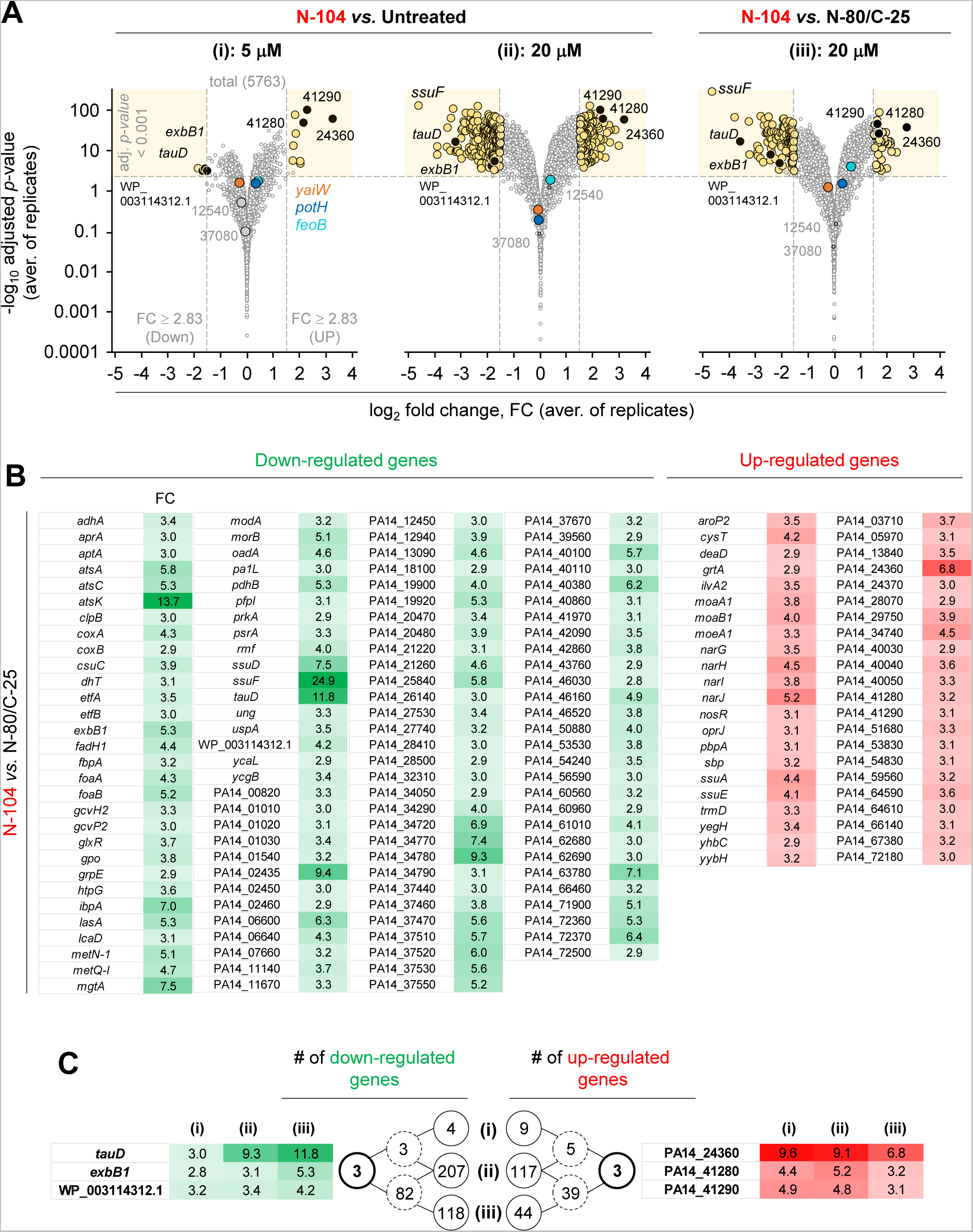
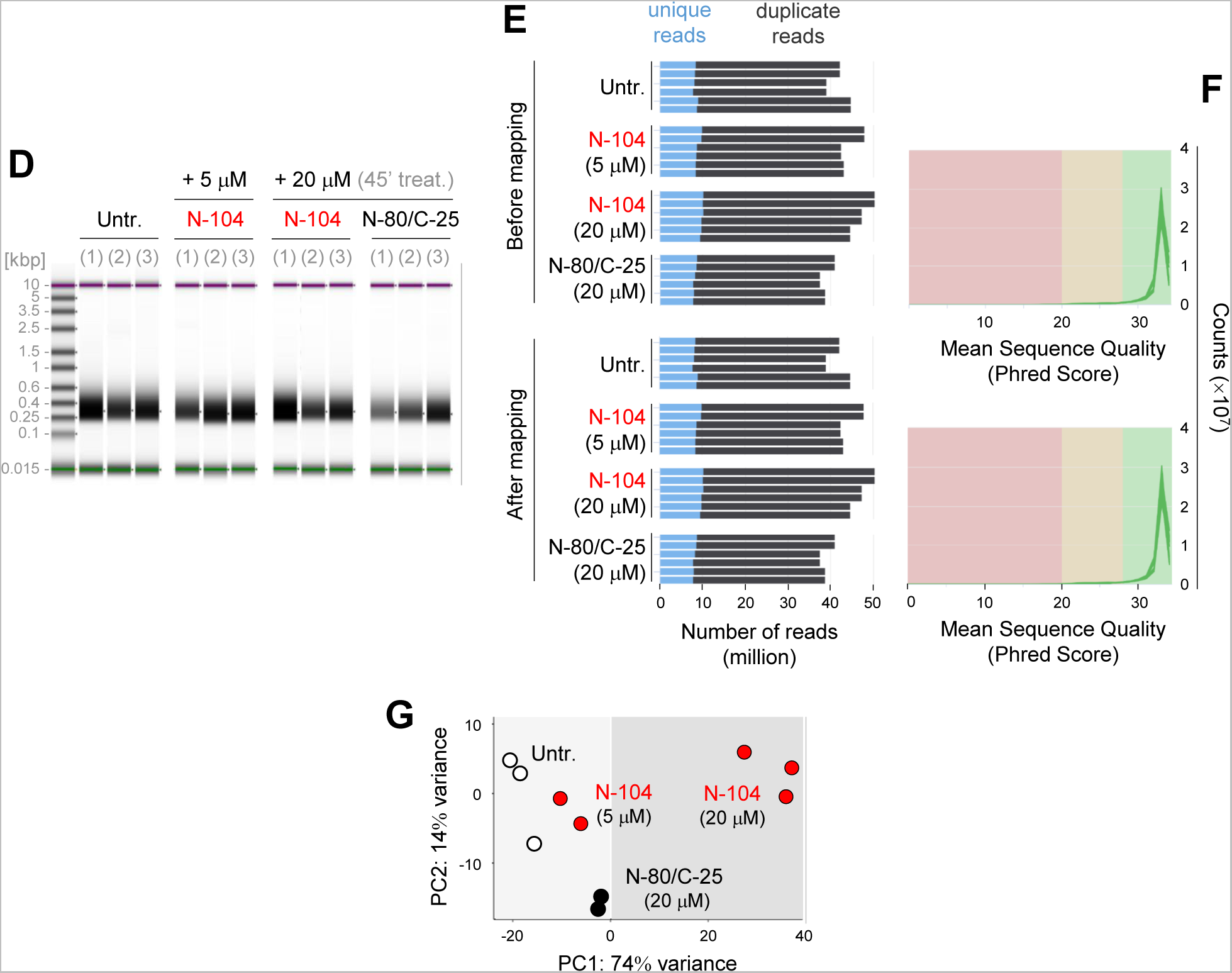
N-104 suppresses gene expression of Fe^3+^ and taurine transport, as well as that of respiratory machinery, as related to Table S6. (A) Differential PA14 gene expression after treatment with (i) 5 or (ii) 20 μM N-104 vs untreated, or (iii) with 20 μM N-104 vs 20 μM N-80/C-25 (negative control). An adjusted *p*-value < 0.001 and fold change ≥ 2.83 were considered significant. For comparison, *feoB*, *potH*, and *yaiW* gene expression is respectively highlighted in aqua, blue and orange. (B) Genes displaying significant down- or up-regulation with fold-change. (C) Shared changes among (i) - (iii). (D) mRNA quality of each biological replicate sample after rRNA removal. (E) Unique and duplicate reads before and after mapping. (F) Sequence quality before and after mapping. (G) Principal component analysis revealing disassociation of 20 µM N-104 from 20 µM N-80/C-25 (20 µM) and untreated samples.

**Figure S9.**
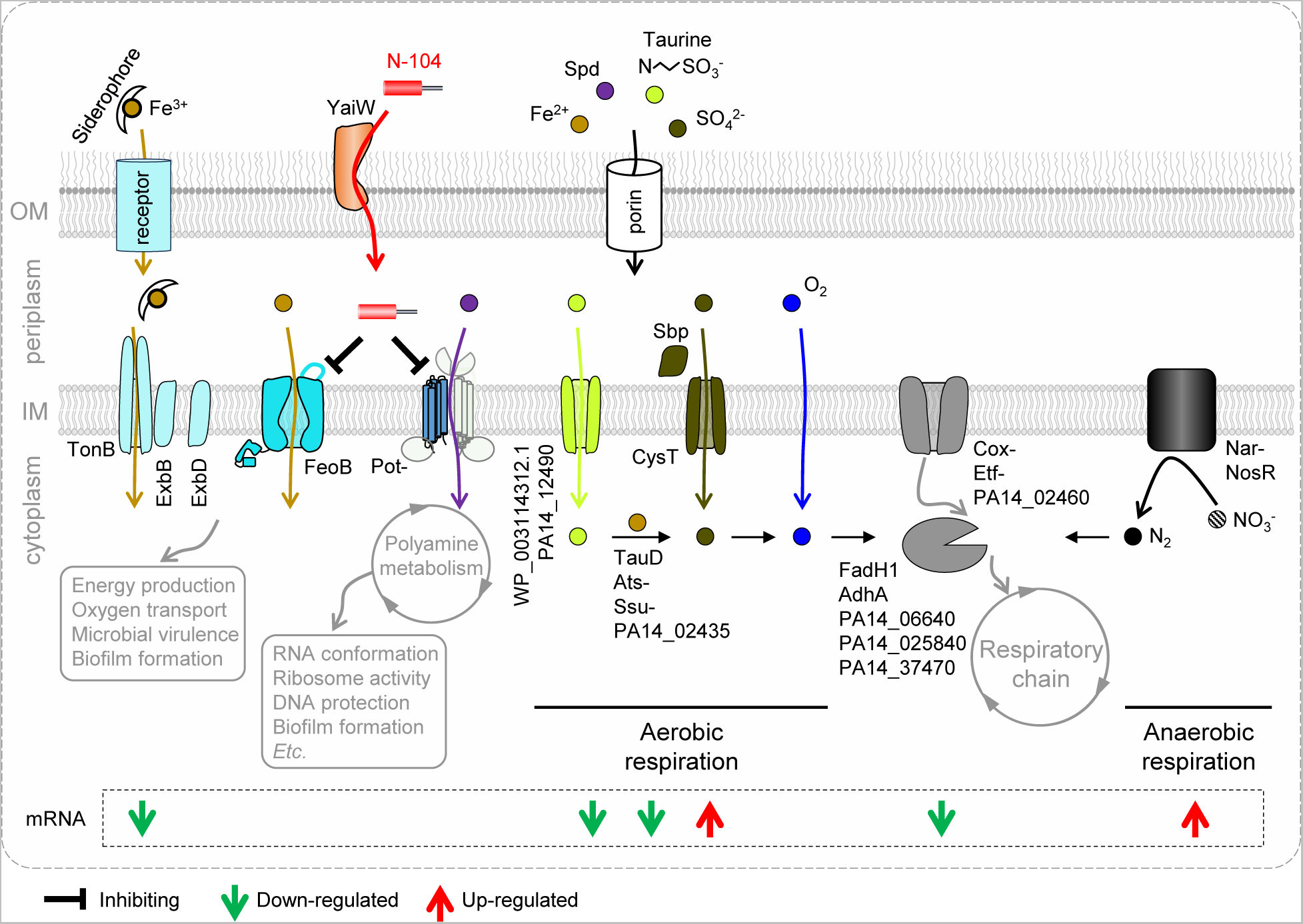
Multiple N-104 killing mechanisms. Beyond suppressing FeoB and PotH, N-104 suppresses the gene expression of Fe^3+^ and taurine transport, as well as that of respiratory machinery.

