## Supplementary material for "Targeting Iron - Respiratory Reciprocity Promotes Bacterial Death": Key Resource Table

### KEY RESOURCES TABLE

| REAGENT or RESOURCE | SOURCE | IDENTIFIER |
|---|---|---|
| Antibodies | | |
| Rabbit polyclonal anti-His-tag | Cell Signaling Technology, Inc | Cat #: 2365S |
| Rabbit polyclonal anti-lacritin C-terminus | DOI: 10.1167/iovs.12-11488 | N/A |
| Rabbit polyclonal anti-lacritin N-terminus | DOI: 10.1167/iovs.11-8729 | N/A |
| Rabbit anti-coagulation factor II/thrombin | Novus Biologicals | Cat #: NBP1-58268 |
| IRDye® 680RD donkey anti-rabbit IgG 2$^{o}$ antibody | LI-COR | Cat #: 926-68073 |
| Bacterial strains | | |
| *Escherichia coli* K12 | DOI: 10.1038/msb4100050 | Wild-type strain K12 |
| *Escherichia coli* Keio collection | | Keio collection, Kan$^{r}$ [a] |
| *Escherichia coli* BL21 (DE3) | BioLabs Inc. | Cat #: C2527I, Chl$^{r}$ [a] |
| *Escherichia coli* OP50 | Caenorhabditis Genetics Center | Wild-type strain, Str$^{r}$ [a] |
| *Pseudomonas aeruginosa* PA14 | DOI: 10.1073/pnas.0511100103 | Wild-type strain PA14, Amp$^{r}$, Kan$^{r}$, Mem$^{r}$ [a] |
| *Pseudomonas aeruginosa* PA14 Δ*feoB* | | *feoB* transposon mutant, Gen$^{r}$ [a] |
| *Pseudomonas aeruginosa* PA14 Δ*spuG* | | *spuG* transposon mutant, Gen$^{r}$ [a] |
| *Pseudomonas aeruginosa* PA14 Δ*yaiW* | | *yaiW* transposon mutant, Gen$^{r}$ [a] |
| *Pseudomonas aeruginosa* K900 | University of Pittsburgh's Charles T. Campbell Ophthalmic Microbiology Laboratory | Clinical isolate; Stain K900, Bac$^{r}$, Van$^{r}$, Cef$^{r}$, Sulf$^{r}$ [a] |
| *Pseudomonas aeruginosa* K1838 | | Clinical isolate; Strain K1838, Bac$^{r}$, Van$^{r}$, Cef$^{r}$, Sulf$^{r}$ [a] |
| *Pseudomonas aeruginosa* K1938 | | Clinical isolate; Strain K1938; Bac$^{r}$, Van$^{r}$, Cef$^{r}$, Sulf$^{r}$ [a] |
| *Pseudomonas aeruginosa* K2408 | | Clinical isolate; Strain K2408; Bac$^{r}$, Van$^{r}$, Cef$^{r}$, Sulf$^{r}$ [a] |
| *Pseudomonas aeruginosa* K2414 | | Clinical isolate; Strain K2414; Bac$^{r}$, Van$^{r}$, Cef$^{r}$, Sulf$^{r}$, Ofl$^{r}$, Mox$^{r}$ [a] |
| *Pseudomonas aeruginosa* K2418 | | Clinical isolate; Strain K2418; Bac$^{r}$, Van$^{r}$, Cef$^{r}$, Sulf$^{r}$ [a] |
| *Pseudomonas aeruginosa* K2421 | | Clinical isolate; Strain K2421, Bac$^{r}$, Van$^{r}$, Cef$^{r}$, Sulf$^{r}$ [a] |
| *Serratia marcescens* K3075 | | Clinical isolate; Strain K3075; Bac$^{r}$, Van$^{r}$, Cef$^{r}$, Sulf$^{r}$, PB$^{r}$ [a] |
| *Stenotrophomonas maltophilia* K2222 | | Clinical isolate; Strain K2222; Bac$^{r}$, Van$^{r}$, Cef$^{r}$, Gen$^{r}$, Tob$^{r}$ [a] |
| Biological samples | | |
| Basal tears from normal human eyes | This paper | N/A |
| Synthetic peptides and chemicals | | |
| N-104 (AQKLLKKFSLLKPWA) | This paper | N/A |

| | | |
|---|---|---|
| K107S (AQSLLKKFSLLKPWA) | This paper | N/A |
| L108S (AQKSLKKFSLLKPWA) | This paper | N/A |
| L109S (AQKLSKKFSLLKPWA) | This paper | N/A |
| K110S (AQKLLSKFSLLKPWA) | This paper | N/A |
| K111S (AQKLLKSFSLLKPWA) | This paper | N/A |
| F112S (AQKLLKKSSLLKPWA) | This paper | N/A |
| L114S (AQKLLKKFSSLKPWA) | This paper | N/A |
| L115S (AQKLLKKFSLSKPWA) | This paper | N/A |
| K116S (AQKLLKKFSLLSPWA) | This paper | N/A |
| W118S (AQKLLKKFSLLKPSA) | This paper | N/A |
| LFW:S (AQKSSKKSSSSSKPSA) | This paper | N/A |
| Cys(N)-N-104 (CAQKLLKKFSLLKPWA) | This paper | N/A |
| FITC-N-104 (FITC-Ahx-AQKLLKKFSLLKPWA) | This paper | N/A |
| JanFl-N-104 (JaneliaFluor-CAQKLLKKFSLLKPWA) | This paper | N/A |
| Chloroalkane-N-104 (Cl-{C11H21O3N}-N-104) | This paper | N/A |
| C-95 (EDASSDSTGADPAQEAGTSKPNEE) | This paper | N/A |
| Cys(N)-C-95 (CEDASSDSTGADPAQEAGTSKPNEE) | This paper | N/A |
| FITC-C-95 (FITC-Ahx-EDASSDSTGADPAQEAGTSKPNEE) | This paper | N/A |
| JanFl-C-95 (JaneliaFluor-CEDASSDSTGADPAQEAGTSKPNEE) | This paper | N/A |
| Chloroalkane-C-95 (Cl-{C11H21O3N}-C-95) | This paper | N/A |
| N-80/C-25 (AKAGKGMHGGVPGG) | This paper | N/A |
| GKY20 (GKYGFYTHVFRLKKWIQKVI) | This paper | N/A |
| GSN (gelsolin) (QRLFQVKGRR) | This paper | N/A |
| CST4 (cystatin S) (SSSKEENRIIPGGI) | This paper | N/A |
| Syndecan-1 pep19-30 (DNFSGSGAGAL) | This paper | N/A |
| FeoB OL1 peptide (INIGGALQP) | This paper | N/A |
| FeoB OL2 peptide (LEDSGYMARAAFVMDRLMQ) | This paper | N/A |
| FeoB OL3 peptide (GAFFGQGGA) | This paper | N/A |
| FeoB OL5 peptide (ATFAA) | This paper | N/A |
| PotH OL2 peptide (WMGILKNNGVLNNFLLWLGVIDQPLTILHTN) | This paper | N/A |
| PotH OL3 peptide (ELLGGPDSIMIGRVLWQEFFNNRDW) | This paper | N/A |
| Acetonitrile | Sigma-Aldrich | Cat #: 34998 |
| Alamar Blue | Invitrogen | Cat #: DAL1100 |
| CHAPS detergent | Fisher Scientific | Cat #: 28299 |
| α-Cyano-4-hydroxycinnamic acid | Sigma-Aldrich | Cat #: 70990 |
| Diethylpyrocarbonate | Fisher Scientific | Cat #: AC170250250 |
| Dithiolthreitol, DTT | Fisher Scientific | Cat #: BP172-25 |
| Dodecyl-β-D-maltopyranoside | Alfa Aesar | Cat #: J66869 |
| Ferrozine | Sigma-Aldrich | Cat #: 160601 |
| Formaldehyde | Polysciences Inc. | Cat #: 18814 |
| β-D-glucopyranoside | Sigma-Aldrich | Cat #: O8001 |
| JaneliaFluor 549 Maleimide | TOCRIS | Cat #: 6500 |
| Lysozyme | Fisher Scientific | Cat #: PI89833 |
| Neocuproine | Sigma-Aldrich | Cat #: N1501 |
| Ni-sepharose™ | GE Healthcare | Cat #: 17-3712-01 |

| | | |
|---|---|---|
| OptiPhase HiSafe 3 scintillation fluid | PerkinElmer | Part #: 1200.437 |
| 1-palmitoyl-2-oleoyl-sn-glycero-3-phosphocholine | Avanti Polar Lipids | Cat #: 850457P |
| 1-palmitoyl-2-oleoyl-sn-glycero-3-phospho-(1'-rac-glycerol) | Avanti Polar Lipids | Cat #: 840457 |
| 1-palmitoyl-2-oleoyl-sn-glycero-3-phosphoethanolamine | Avanti Polar Lipids | Cat #: 850757 |
| Pierce™ Control Agarose Resin | Fisher Scientific | Cat #: 26150 |
| Proteinase K | Fisher Scientific | Cat #: EO0491 |
| [1,4-$^{14}$C]putrescine | American Radiolabeled Chemicals | Cat #: ARC-0245-250 |
| Putrescine | MP Biomedicals | Cat #: 100450 |
| Rhodamine chloroalkane | Promega® | Cat #: G3221 |
| RNaseZap™ | Invitrogen | Cat #: AM9780, AM9782 |
| RNA-protect Bacteria Reagent | Qiagen | Cat #: 76506 |
| Sephadex™ G-10 | Cytiva | Cat #: 17001001 |
| [1,4-$^{14}$C]spermidine | American Radiolabeled Chemicals | Cat #: ARC-3138-50 |
| Spermidine | Acros organics | Cat #: 132740010 |
| SulfoLink® Coupling Resin | Fisher Scientific | Cat #: 20401 |
| Sytox orange | Fisher Scientific | Cat #: S11368 |
| Trifluoroacetic acid | Chem-Impex International | Cat #: 02883 |
| Trypsin EDTA | Gibco | Cat #: 25200-056 |
| Critical commercial assays | | |
| EndoFree® Plasmid Maxi Kit | Qiagen | Cat #: 12362 |
| RNeasy mini kit | Qiagen | Cat #: 74104 |
| NEBNext® rRNA Depletion Kit (Bacteria) | New England BioLabs | Cat #: E7850L |
| BCA protein assay | bioWORLD | Cat #: 20831001-1 |
| Deposited data | | |
| RNA-seq data | NCBI GEO data set | Accession ID: GSE253123 |
| Tear proteomics | MassIVE data set | Accession ID: MSV000094085 |
| PA14 proteomics | MassIVE data set | Accession ID: MSV000094086 |
| Experimental models: Cell lines | | |
| Sheep red blood cells | MP Biomedicals | Cat #: 0855876 |
| Experimental models: Organisms/strains | | |
| *Caenorhabditis elegans*: Wild-type strain N2 | Caenorhabditis Genetics Center | Wild-type strain N2 |
| Mouse | The Jackson Laboratory | C57BL/6 (female, 7 weeks old) |
| Oligonucleotides | | |
| Primers for site-directed mutagenesis experiments, see Table S4 | This paper | N/A |
| Recombinant DNA | | |
| *feoB* | DOI: 10.1016/j.pep.2014.06.012 | Addgene ID: 216753 (*P. aeruginosa*'s *feoB* in pET41-a(+) vector with T7 promoter & *lac*-operator system, C-term. 8xHis, Kan$^{r}$ [(a)]) |

| | | |
|---|---|---|
| *feoB* OL1 deletion | This paper | Addgene ID: 216741 |
| *feoB* OL2 deletion | This paper | Addgene ID: 216742 |
| *feoB* OL3 deletion | This paper | Addgene ID: 216743 |
| *feoB* OL4 deletion | This paper | Addgene ID: 216744 |
| *feoB* OL5 deletion | This paper | Addgene ID: 216745 |
| *feoB* OL2/4 substitution | This paper | Addgene ID: 216746 |
| *feoB* OL3/4 substitution | This paper | Addgene ID: 216747 |
| *feoB* OL5/4 substitution | This paper | Addgene ID: 216748 |
| *potH* | DNASU Plasmid Repository (https://dnasu.org/DNASU/Home.do) | ID: EcCD00397460 (*E. coli*'s *potH* in PCDF Bravo vector with T7 promoter & *lac*-operator system, C-term. 10xHis, Str$^{r}$ [a]) |
| *potH* OL1 deletion | This paper | Addgene ID: 216749 |
| *potH* OL2 deletion | This paper | Addgene ID: 216750 |
| *potH* OL3 deletion | This paper | Addgene ID: 216751 |
| *potH* OL3/2 substitution | This paper | Addgene ID: 216752 |
| *cyto-Halo* | DOI: 10.1021/acsinfecdis.2c00435 | HaloTag in pET21(b)+ vector with T7 promoter & lac-operator system, Amp$^{r}$ [a] |
| *peri-Halo* | | Cyto_Halo containing an amine terminal recognition sequence in pET21(b)+ vector with T7 promoter & lac-operator system, Amp$^{r}$ [a] |
| Software and algorithms | | |
| DESeq2 (version 1.38.3) | Bioconductor | https://support.bioconductor.org |
| GraphPad Prism (version 10.2.0) | | |
| IDEAS® program (version 6.2) | Amnis® system | https://www.emdmillipore.com/US/en/20150122_174404 |
| ImageJ (version 1.48v) | National Institute of Health | https://imagej.net/ij/docs/install/windows.html |
| Image Studio™ Lite (version 5.2.5) | LI-COR | https://www.licor.com/bio/image-studio-lite/ |
| NEBase Changer (version 2.4.3) | New England Biolabs Inc. | https://nebasechanger.neb.com/?# |
| SnapGene Viewer (version 5.0.4) | SnapGene | https://www.snapgene.com/support/downloads |
| Other | | |
| Acetic acid | Fisher Scientific | Cat #: A38SI-212 |
| Ammonium acetate | Fisher Scientific | Cat #: A639 |
| Ampicillin | Fisher Scientific | Cat #: BP1760-5 |
| Ascorbic acid | Sigma-Aldrich | Cat #: A4544 |
| 2,2'-bipyridyl | Sigma-Aldrich | Cat #: D216305 |
| L-cysteine | Alfa Aesar | Cat #: A10435 |
| Dimethyl sulfoxide | Fisher Scientific | Cat #: D128-500 |

| | | |
|---|---|---|
| $FeSO_4$ | Sigma-Aldrich | Cat #: 215422 |
| Fish gelatin | Sigma-Aldrich | Cat #: G7765 |
| Formic acid | Fisher Scientific | Cat #: A117-50 |
| Gentamicin | Alfa Aesar | Cat #: J62834 |
| Imidazole | Fisher Scientific | Cat #: 122025000 |
| Isopropyl-β-D-1-thiogalactopyrannoside | Fisher Scientific | Cat #: R0392 |
| Kanamycin | Fisher Scientific | Cat #: BP906-5 |
| Laemmli buffer (4x) | Bio-Rad | Cat #: 1610737 |
| LB Broth medium | Fisher Scientific | Cat #: BP9723-2 |
| Luria-Bertani agar | Fisher Scientific | Cat #: BP1425-2 |
| M9 minimal salts | Gibco | Cat #: A1374401 |
| β-mercaptoethanol | Sigma-Aldrich | Cat #: M7522 |
| Mini-PROTEIN TGX Gel | Bio-Rad | Cat #: 4561094 |
| MWCO (50 kDa) | Amicon® | Cat #: UFC505024 |
| MWCO (30 kDa) | Amicon® | Cat #: UFC503024 |
| MWCO (5 kDa) | Amicon® | Cat #: UFC900596 |
| Nitrocellulose membrane | GE Healthcare Life Sciences | Cat #: 10600001 |
| Protease inhibitor cocktail | cOmplete Mini | Cat #: 11836153001 |
| RDD buffer | Qiagen | Cat #: 79254 |
| Streptomycin | Sigma-Aldrich | Cat #: S650-1 |
| TE buffer | Invitrogen | Cat #: 12-090-015 |
| Terrific broth medium | Research Products International | Cat #: T15100-1000.0 |
| Triton X-100 | Bio-Rad | Cat #: 161-0407 |
| Tween-20 | Bio-Rad | Cat #: 1706531 |

Key:
(i) antibiotics ($^{a}$): Amp, ampicillin; Bac, bacitracin; Cef, cefazolin; Chl, chloramphenicol; Gen, gentamicin; Kan, kanamycin; Mem, meropenem; Mox, moxifloxacin; Ofl, ofloxacin; PB, polymyxin B; Str, streptomycin; Sulf, sulfasoxazole; Tob, tobramycin; Van, vancomycin;
(ii) antibiotic resistance ($^{r}$)

### SUPPLEMENTAL FILE INFORMATION

**SUPPLEMENTARY INFORMATION:**

**Formulaic analysis of Figures 3I and 3J:**

For analysis of the ChloroAlkane Penetration Assay, '$\mathrm{I}$' corresponds to the fluorescence intensity of chloroalkane-modified fluorescent dye, '$\varepsilon$' is the fluorescence signal coefficient, '$[\mathrm{A}]_0$' denotes the initial concentration of chloroalkane-modified fluorescent dye when '$[\mathrm{B}]$' equals zero. $[\mathrm{B}]$ is the concentration of Chloroalkane tagged N-104 or Chloroalkane tagged C-95, and '$\mathrm{I}_{\mathrm{base}}$' represents the background signal:

$$\mathrm{I} = \varepsilon\,([\mathrm{A}]_0 - [\mathrm{B}]) + \mathrm{I}_{\mathrm{base}}$$

Substituting $[\mathrm{A}]_0$ with $\frac{\mathrm{I}_0 - \mathrm{I}_{\mathrm{base}}}{\varepsilon}$, and subtracting $\mathrm{I}_{\mathrm{base}}$ from all data gives:

$$\mathrm{I} = \mathrm{I}_0 - \varepsilon.[\mathrm{B}]$$

To then establish the following relationship, where 'peri' and 'cyto' refer to the periplasm and cytoplasm respectively:

$$[\mathrm{B}]_{\mathrm{peri}} = \frac{\mathrm{I}_{0,\mathrm{peri}} - \mathrm{I}_{\mathrm{peri}}}{\varepsilon} \quad \text{(AND)} \quad [\mathrm{B}]_{\mathrm{cyto}} = \frac{\mathrm{I}_{0,\mathrm{cyto}} - \mathrm{I}_{\mathrm{cyto}}}{\varepsilon}$$

This facilitates graphical representation of $\frac{\mathrm{I}_{0,\mathrm{peri}} - \mathrm{I}_{\mathrm{peri}}}{\varepsilon}$ or $\frac{\mathrm{I}_{0,\mathrm{cyto}} - \mathrm{I}_{\mathrm{cyto}}}{\varepsilon}$ as a function of '$[\mathrm{B}]_{\mathrm{extr}}$', or of $[\mathrm{B}]_{\mathrm{cyto}}$ or $[\mathrm{B}]_{\mathrm{peri}}$. $[\mathrm{B}]_{\mathrm{extr}}$ is the initial treatment concentration of Chloroalkane tagged N-104 or Chloroalkane tagged C-95. $[\mathrm{B}]_{\mathrm{peri}}$ and $[\mathrm{B}]_{\mathrm{cyto}}$ are the respective periplasmic and cytoplasmic concentrations of each.
